# Evolution under Stochastic Transmission: Mutation-Rate Modifiers

**DOI:** 10.1101/2025.11.04.686407

**Authors:** Elisa Heinrich-Mora, Marcus Feldman

## Abstract

In evolutionary models of large populations, it is common to analyze the effects of cyclic or random variation in the parameters that describe selection. It is less common, however, to study how stochasticity in the genetic transmission process itself affects evolutionary outcomes. Suppose that a gene locus has alleles *A* and *a* under constant selection. This locus is linked to a modifier locus with alleles *M*_1_ and *M*_2_, which control the mutation rate from *A* to *a*. The *Reduction Principle* states that, near a mutation–selection balance where *M*_1_ is fixed with mutation rate *u*_1_, a rare allele *M*_2_ can invade if its associated rate *u*_2_ is lower than *u*_1_. This result, valid for both haploids and diploids, assumes constant mutation rates through time. We extend this framework by allowing the mutation rate associated with *M*_2_ to fluctuate randomly across generations, denoted as *u*_2,*t*_. In this stochastic setting, the condition for invasion by a new modifier allele depends not only on the resident mutation rate *u*_1_ and the mean mutation rate *u*_2_ associated with the invading allele, but also on the temporal distribution of *u*_2,*t*_, the strength of selection at the *A/a* locus, and the recombination rate between *M*_1_*/M*_2_ and *A/a*. The analysis shows how stochasticity and recombination in transmission do not simply modify the magnitude of evolutionary change predicted under deterministic assumptions. Instead, through their interaction with selection and linkage, they can generate conditions under which the direction of modifier evolution is qualitatively reversed relative to the deterministic *Reduction Principle*.

## Introduction

An important goal of evolutionary biology is to identify the forces that shape the structure and dynamics of genetic systems. Standard population–genetic models often treat aspects of transmission (e.g. mutation, recombination rates) and genetic structure (e.g. dominance, epistasis) as fixed. Empirical work, however, has shown that these parameters can themselves be subject to evolutionary modification. *Modifier theory* makes this point explicit by analyzing loci whose alleles affect the transmission and/or genetic structure at a primary locus under selection—for example, modifiers of mutation, recombination, or dominance. Invasion analysis of a rare modifier allele identifies which changes are initially favored, and provide insight into how the such parameters might evolve. It is useful to distinguish modifiers with *direct* fitness effects—those that alter viability or fertility through changes in parameters such as dominance or epistasis—from selectively *neutral* modifiers, which have no intrinsic effect on fitness [1], such as those influencing transmission parameters (e.g., mutation or recombination rates). The former change in frequency through both their direct fitness effects and associations with selected loci, whereas the latter evolve solely through such associations.

A key result from this framework is the *Reduction Principle*, which states that modifier alleles that reduce mutation rates at loci under mutation–selection balance, or that reduce recombination rates between loci with epistatic interactions that generate linkage disequilibrium, increase in frequency due to their indirect effects on the selected loci [8, 18, 24, 1, 9, 2, 7, 3, 20, 10]. The principle holds under assumptions of infinite population size, random mating, constant viability selection, and deterministic dynamics—prompting studies to examine departures from these conditions.

A substantial body of theoretical work has investigated how temporal fluctuations in selection parameters can influence long-term evolutionary outcomes. These studies treat fitness values as random variables that fluctuate across generations. Gillespie [14, 15], Cook and Hartl [6], and Karlin and Liberman [16] analyzed such scenarios, demonstrating that fluctuating election can—under specified conditions—maintain polymorphism and favor the allele with the highest geometric mean fitness across time, which is shaped by both the mean and higher moments of the fitness distribution. Related results in population growth theory show that, in randomly varying environments, multiplicative growth is governed by the geometric mean of growth factors [19]. In these analyses, stochasticity enters through selection; transmission parameters remain fixed.

*Modifier theory* has also been extended to scenarios with variable selection regimes. In these models, the fitness landscape changes over time while the fundamental assumptions of modifier theory are retained. Indirect selection on modifier alleles can arise from spatial or temporal fluctuations in selection coefficients [5, 4], and environmental variability can favor nonzero mutation or switching rates by altering the geometric mean fitness of genotypes across cycles [21]. Recent simulation studies of multigenic mutation–rate modifiers in sexual populations further demonstrate that the form of selection acting on a quantitative trait—stabilizing versus directional—can bias the evolution of mutation rates through associations between modifier alleles and individuals in the phenotypic tails [23]. Together, these findings suggest that the evolution of transmission parameters such as mutation rate depends not only on the expected magnitude of environmental change but also on its distribution and directional properties.

There is evidence that transmission parameters can vary through endogenous processes even under constant external selective conditions. In *Arabidopsis thaliana*, mutation–accumulation experiments show genome–wide heterogeneity in mutation rates that correlates with epigenetic and physical genomic features rather than with direct natural selection [22]. In *Escherichia coli*, long–term evolution experiments reveal “hypermutator” alleles—typically defects in DNA repair—whose effects on mutation rate are not strictly monotone with environmental conditions [26]. In humans, variation in recombination rate is influenced by genetic variants—most notably in *PRDM9* and *RNF212* —as well as by structural variation, with little evidence for direct effects on fitness [11]. These observations suggest that the mechanisms of transmission are themselves evolvable traits, capable of generating variation even in the absence of external change.

If endogenous processes generate fluctuations in epigenetic or physical genomic features that influence transmission parameters, a natural question arises: are genetic modifiers subject to selection not only through their mean effects, but also through their *variability* across generations? To address this, we analyzed *A. thaliana* essential and lethal genes (Appendix C) using gene–level mutation data from Monroe et al. [22]. In multivariable models, chromatin marks associated with active repair (H3K4me1, H3K4me3, H3K36ac) were linked to large reductions in mean mutation rates and, more strongly, reduction in their variance. Although formal selection tests were underpowered, regressions of a standardized Tajima’s *D* score on moments of the mutation–rate distribution showed similar tendencies: stronger purifying selection coincided with lower means and lighter tails. These patterns are consistent with selection in functionally constrained genomic regions, acting to reduce not only the average mutation rate but also its variability. Modifier alleles that permit occasional high mutation rates may be disfavored relative to those that suppress variance while maintaining a comparable mean mutation rate.

We therefore consider models of stochasticity in the *transmission* process. Rather than fixing transmission parameters while allowing selection to vary, we hold selection constant and allow transmission parameters to fluctuate across generations. The focus is the invasion of a rare modifier allele that reduces the average transmission rate while introducing intergenerational variance in that same parameter. The central question is whether indirect selection on the modifier still conforms to the *Reduction Principle* or whether variance in the transmission parameter alters this outcome.

Let *T* denote a transmission parameter with mean *µ* = *E*[*T*] and variance *σ*^2^ = Var(*T*). In each generation, the value of *T* is drawn independently from this distribution. Classical modifier theory treats *T* as constant. When *T* fluctuates, the variance *σ*^2^, even with constant *µ*, may affect the initial increase of a new modifier allele. This effect is captured by Lyapunov exponents of the linearized recursions for the modifier haplotype frequencies, which depend on the full distribution of transmission parameters. Hence invasion depends on both the average effect on transmission and higher–order statistics of the transmission process. Variability in transmission is therefore not simply an extraneous source of noise, but a factor that can alter the magnitude and direction of evolutionary outcomes. Extending modifier theory to include fluctuating transmission thus provides a general framework for analyzing how endogenous variability in transmission rates influences the evolution of genetic systems.

### Mutation Modification

Consider a large, randomly mating population with discrete generations. A single locus under constant selection carries alleles *A* and *a*; a linked, selectively neutral modifier locus carries alleles *M*_1_ and *M*_2_ that determine the forward mutation rate at the selected locus. Under *M*_1_, the mutation rate is constant at *u*_1_ across all generations, and the selected locus is at its deterministic mutation–selection balance. At this equilibrium, a rare modifier allele that reduces the constant rate (*u*_2_ *< u*_1_) is favored—the *Reduction Principle* [17, 1, 9, 2, 20]. Here we ask whether that principle holds when the modifier allele influences not only the mean mutation rate but also its variability. Specifically, *M*_2_ generates a mutation rate sequence *{u*_2,*t*_*}* with expectation *E*[*u*_2,*t*_] = *u*_2_ and variance Var(*u*_2,*t*_) = *σ*^2^ across generations, while *u*_1_ remains constant and selection at *A/a* is time-invariant. Our objective is to derive the conditions under which *M*_2_, introduced at low frequency near a mutation-selection balance with *M*_1_, can invade in haploid or diploid populations, and to elucidate how the temporal distribution of the mutation rate, together with selection strength and recombination between the modifier and selected loci, affect these invasion dynamics.

## 1 Model Set-up

A large population is considered to have two biallelic loci: a selected locus (*A/a*) under constant selection and a linked, selectively neutral modifier (*M*_1_*/M*_2_) that controls the *forward* mutation rate *A* → *a*. Back mutation *a* → *A* is absent. The recombination fraction between loci is 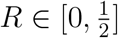, constant through time. The life cycle within each generation is

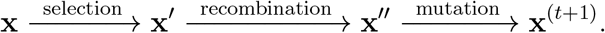

Let **x** = (*x*_1_, *x*_2_, *x*_3_, *x*_4_) denote haplotype frequencies (*AM*_1_, *aM*_1_, *AM*_2_, *aM*_2_) with Σ_*i*_ *x*_*i*_ = 1. After selection and recombination the frequencies are **x**^*′*^ and **x**^*′′*^, respectively; **x**^(*t*+1)^ denotes the state after mutation (start of generation *t*+1).

### Selection

Selection acts only at *A/a*. For haploids, viabilities are *W*_*A*_ = 1 and *W*_*a*_ = 1 − *s* with *s* ∈ (0, 1], yielding mean fitness

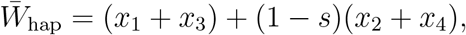

and

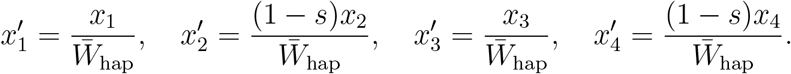

For diploids, viabilities at the selected locus are additive:

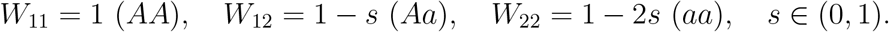

Random mating (Hardy–Weinberg before selection) and viability selection entail that the mean fitness in diploids is

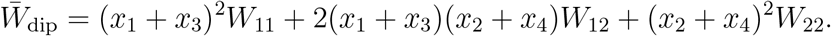

The resulting gamete frequencies, 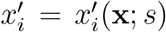 after selection are the standard two-locus frequencies with viabilities *W*_11_, *W*_12_, *W*_22_.

### Recombination

Recombination acts on **x**^*′*^ in haploids and diploids at the gamete level. With linkage disequilibrium after selection, 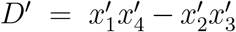, after recombination we have

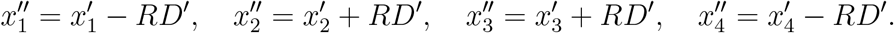

### Mutation

Forward mutation *A* → *a* is modifier dependent and operates on *A*-bearing gametes after recombination. In the haploid case, *M*_1_ and *M*_2_ produce constant per-generation rates *u*_1_, *u*_2_ ∈ [0, 1):

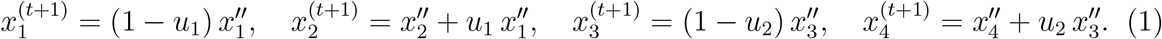

In diploids, the mutation rate depends on the modifier *genotype*, and gametes produced from *M*_1_*M*_1_, *M*_1_*M*_2_, and *M*_2_*M*_2_ zygotes have rates *u*_1_, *u*_2_, *u*_3_; the evolutionary recursions are obtained by applying these rates to the appropriate gamete contributions after selection and recombination.

### Full recursion

Substituting the parameters at each step yields:

- *Haploids:*

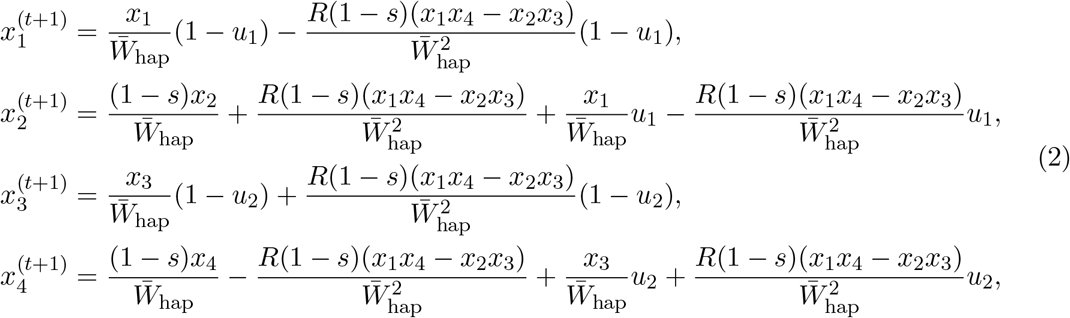

with 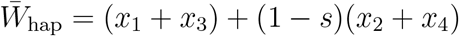.
- *Diploids* [25]:

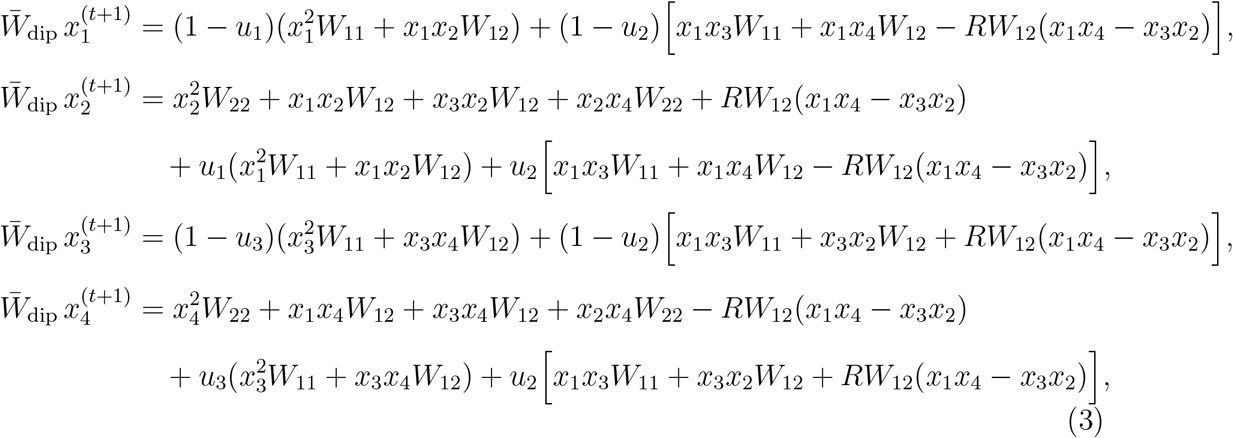

with 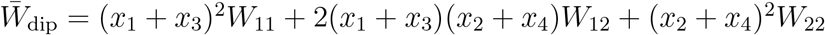.

Recursion systems (2) and (3) are the basis for the invasion analysis in which *M*_2_ is initially rare and the resident *M*_1_ is at its mutation–selection equilibrium.

## 2 Invasion Analysis of Modifier Allele *M*_2_

We ask whether a rare modifier allele *M*_2_ increases in frequency when introduced near a resident population fixed for *M*_1_. Let **x**_*t*_ = (*x*_1_, *x*_2_, *x*_3_, *x*_4_)^⊤^ denote the haplotype frequencies of *AM*_1_, *aM*_1_, *AM*_2_, and *aM*_2_ at generation *t*, and let 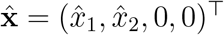 denote the resident equilibrium (mutation–selection balance) at the selected locus *A/a*. Using the life–cycle order specified in Section 1, the one–locus subsystem at equilibrium satisfies *x*_*i,t*+1_ = *x*_*i,t*_, with

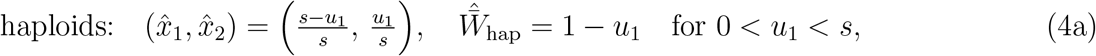

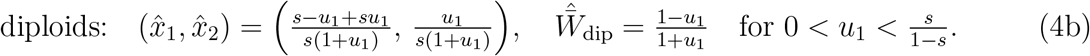

Linearizing the full two–locus recursions system at 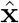 yields the Jacobian

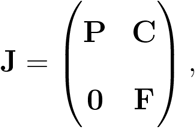

where **P** describes the resident (*AM*_1_, *aM*_1_) subsystem, **C** captures first–order coupling from residents to rare haplotypes, and **F** governs the rare subsystem associated with *M*_2_. Because **J** is block–triangular, the eigenvalues of **F** determine the local stability of the *M*_1_ equilibrium with respect to invasion by *M*_2_. Consequently, the frequencies of the rare haplotypes ***v***_*t*_ = (*x*_3,*t*_, *x*_4,*t*_)^⊤^ satisfy

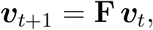

and invasion occurs when the dominant eigenvalue (Perron root) of **F**, *ρ*(**F**), exceeds unity. Substituting 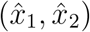 from (4) into the *x*_3,*t*+1_ and *x*_4,*t*+1_ components of the recursion systems (2) and (3) and discarding higher–order terms gives the explicit forms:

- *Haploids*.

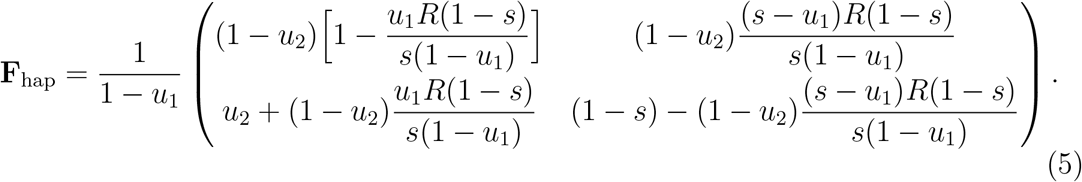
- *Diploids*.

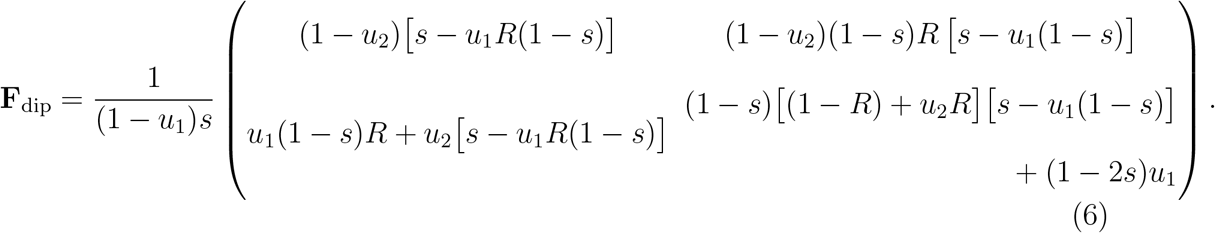

Thus the invasion dynamics depend on selection *s*, recombination *R*, and the resident mutation rate *u*_1_ through the entries of **F**. Because the modifier has no direct fitness effect, its initial change is driven by the dynamics of the *M*_2_–bearing haplotypes (*AM*_2_, *aM*_2_) near the resident haplotypes *AM*_1_, *aM*_1_. Selection at the *A/a* locus generates the associations that facilitate this growth, whereas recombination modulates these associations each generation, as reflected in the *R*–dependent terms of **F**. The condition *ρ*(**F**) *>* 1 defines the criterion for invasion.

### 2.1 Deterministic Case (*u*_2_ constant)

Near the resident mutation–selection equilibrium 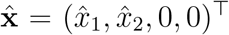 with modifier *M*_1_, and forward mutation rate *u*_1_, introduce a rare modifier *M*_2_ with constant mutation rate *u*_2_. Linearizing at 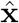 yields ***v***_*t*+1_ = **F *v***_*t*_, where ***v***_*t*_ = (*x*_3,*t*_, *x*_4,*t*_)^⊤^ collects the rare haplotype frequencies and **F** is a nonnegative 2*×*2 matrix (with positive entries in the parameter ranges considered). By Perron–Frobenius [12], **F** has a simple dominant eigenvalue *λ*_+_ = *ρ*(**F**) *>* 0, and *M*_2_ increases when *λ*_+_ *>* 1.

The characteristic polynomial is *p*(*λ*) = *λ*^2^ − *τλ* + *δ*, where *τ* = tr(**F**) and *δ* = det(**F**). Since *p*(*λ*) = (*λ* − *λ*_−_)(*λ* − *λ*_+_) with 0 *< λ*_−_ ≤ *λ*_+_, we have *p*(1) = (1 − *λ*_−_)(1 − *λ*_+_). Thus, if *λ*_−_ *<* 1 (which holds at the resident equilibrium under the admissible parameter ranges), then *p*(1) *<* 0 if *λ*_+_ *>* 1. Because *p*(1) = 1 − *τ* + *δ*, it suffices to evaluate *p*(1) at the resident equilibrium. Direct calculation gives

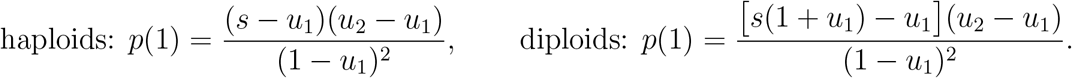

For haploids, equilibrium requires 0 *< u*_1_ *< s*, so *s* − *u*_1_ *>* 0; for the additive diploid model, 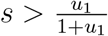, so *s*(1 + *u*_1_) − *u*_1_ *>* 0. Under these conditions, *p*(1) has the sign of (*u*_2_ − *u*_1_), and therefore

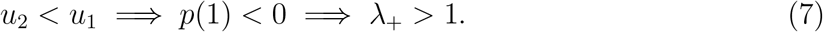

This is the *Reduction Principle*: a modifier allele that lowers the mutation rate (*u*_2_ *< u*_1_) will invade.

#### Effect of recombination on *M*_2_ invasion

Although recombination *R* does not affect the *criterion* for invasion—since *p*(1) = 1 − *τ* + *δ* is independent of *R*—it affects the *magnitude* of growth through the Perron root 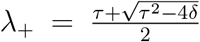. Writing the characteristic equation *p*(*λ*) = *λ*^2^ − *τλ* + *δ* = 0 and differentiating implicitly with respect to *R* gives

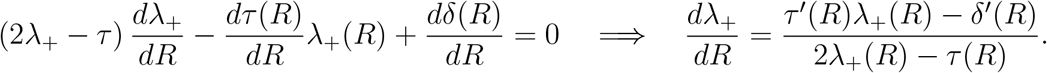

Since *p*(1) is *R*–invariant, −*τ* ^*′*^(*R*) + *δ*^*′*^(*R*) = 0, hence *δ*^*′*^(*R*) = *τ* ^*′*^(*R*) and

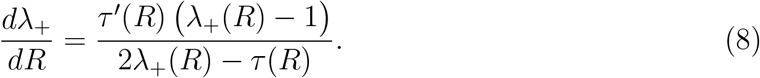

For a 2 *×* 2 matrix, 2*λ*_+_ − *τ* = *λ*_+_ − *λ*_−_ *>* 0, so sign *dλ*_+_*/dR* = sign *τ* ^*′*^(*R*) sign *λ*_+_ − 1. It remains to compute *τ* ^*′*^(*R*).

- *Haploids*. From (5),

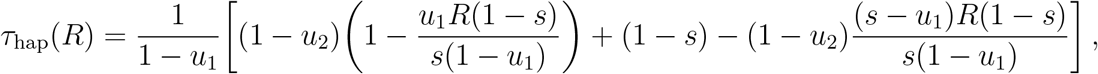

so

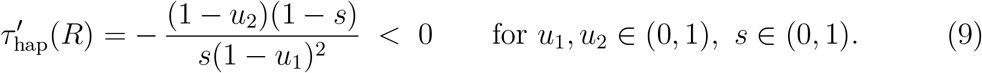
- *Diploids*. From (6), the trace is

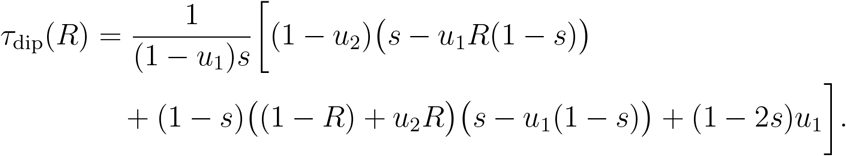

hence

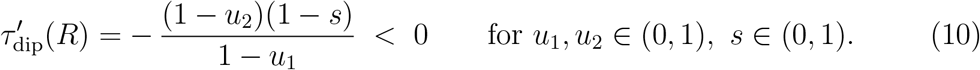 Substituting (9)–(10) into (8) yields

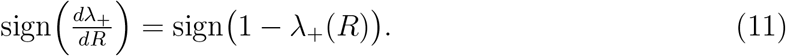

Thus, recombination shifts *λ*_+_ monotonically toward 1: it decreases *λ*_+_ when *λ*_+_ *>* 1 and increases it when *λ*_+_ *<* 1. If *u*_2_ = *u*_1_, then *λ*_+_(*R*) ≡ 1 for all *R*, consistent with (8).

### 2.2 Stochastic Mutation Rate (*u*_2,*t*_)

Assume the resident population is at mutation–selection equilibrium (Eq. (4)) with forward mutation rate *u*_1_, satisfying 0 *< u*_1_ *< s* in haploids and 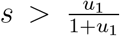 in diploids. Let the invading modifier allele *M*_2_ produce a forward mutation rate that varies independently across generations,

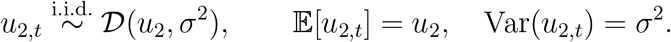

Linearization at 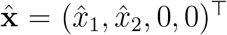 yields the rare–haplotype recursion ***v***_*t*+1_ = **F**_*t*_(*R, u*_2,*t*_) ***v***_*t*_ with ***v***_*t*_ = (*x*_3,*t*_, *x*_4,*t*_)^⊤^ and **F**_*t*_ given by (5) or (6) for the realized *u*_2,*t*_ and constant 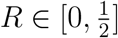.

If *E*[log∥**F**_*t*_∥] *<* ∞, the top Lyapunov exponent

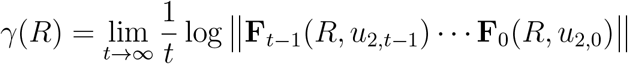

exists almost surely and is norm–independent [13]. Invasion occurs if *γ*(*R*) *>* 0 (conversely, *M*_2_ is lost if *γ*(*R*) *<* 0; when *γ*(*R*) = 0, we cannot distinguish loss from invasion to linear order).

For each *t*, decompose **F**_*t*_(*R*) as **F**_*t*_(*R*) = **A**_*t*_ + *R* **B**_*t*_, i.e. separate the *R*–independent and *R*–linear parts. Rewrite (5)–(6) as,

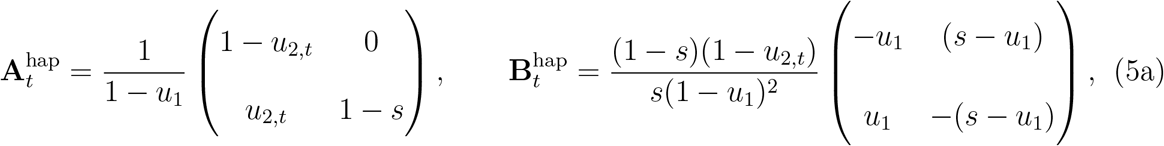

for haploids, and for diploids,

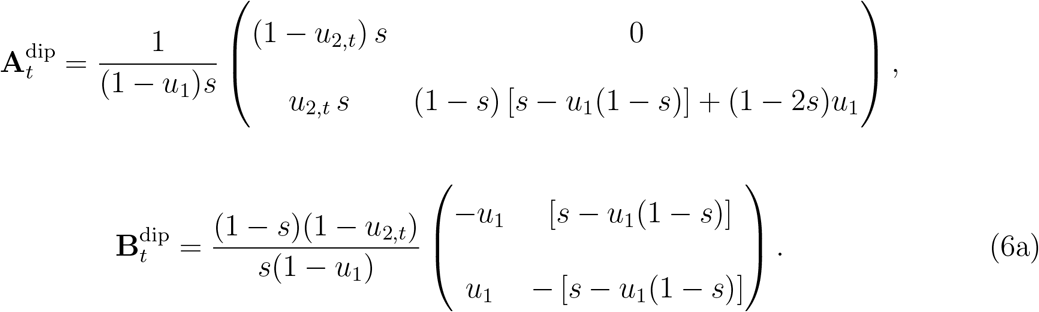

The form of **A**_*t*_ in (5a) and (6a) match **F**_*t*_(0) from the *R* = 0 analysis. The matrices **B**_*t*_ capture the same–generation linear response to recombination; in particular, their column sums are zero, reflecting that recombination redistributes haplotypes across backgrounds. Two implications follow directly:

1. In the case *R* = 0, **F**_*t*_(0) = **A**_*t*_ is lower triangular; the Lyapunov exponent reduces to the time average of the log of the dominant diagonal term.
2. For *R >* 0 and *u*_2,*t*_ *>* 0, all entries of **F**_*t*_(*R*) are strictly positive. Hence each **F**_*t*_(*R*) is primitive. If ℙ(*u*_2,*t*_ *>* 0) *>* 0, primitivity occurs infinitely often almost surely, which suffices for a unique top Lyapunov exponent [13]. *R* enters *linearly* at each generation via **B**_*t*_, but the map *R 1*→ *γ*(*R*) is *not* necessarily linear (or even monotone) because *γ*(*R*) is a limit of logs of products of non-commuting random matrices.

#### Baseline at *R* = 0

When *R* = 0, **F**_*t*_(0) matrices are lower triangular, so products remain triangular. The top Lyapunov exponent is the maximum of the time–averaged logarithms of the diagonal entries.

- For haploids,

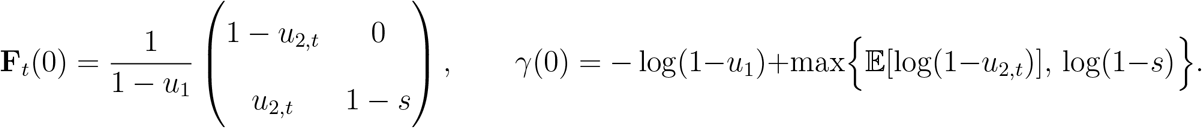
- For diploids,

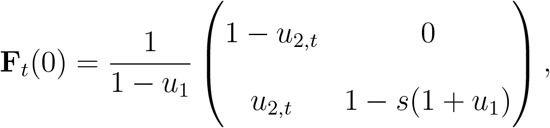

and, provided 1 − *s*(1 + *u*_1_) *>* 0,

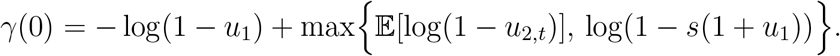

The eigenvalues of **F**_*t*_(0) are therefore

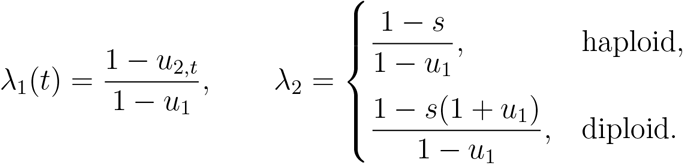

##### Regime 1 (λ_1_(*t*) term dominates)

Because the resident population is at mutation–selection balance, *λ*_2_ *<* 1, in both haploids and diploids. Hence, with complete linkage (*R* = 0), invasion can occur only through *λ*_1_(*t*). The stochastic invasion condition is

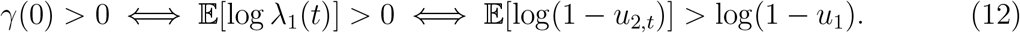

Equation (12) states that *M*_2_ increases when the expected log of its per-generation probability of transmitting allele *A*, namely (1 − *u*_2,*t*_), exceeds that of the resident. Because log(1 − *x*) is strictly concave on [0, 1), Jensen’s inequality implies

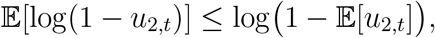

with equality only if *u*_2,*t*_ is constant. Hence temporal variance in *u*_2,*t*_ reduces *E*[log(1 − *u*_2,*t*_)] relative to a deterministic rate with the same mean, shrinking the parameter region in which invasion is possible. Increasing *u*_1_ increases *γ*(0) (since − log(1 − *u*_1_) increases), expanding the invasion region. When (12) holds, the growth rate is

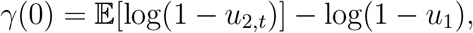

which is independent of *s* and ploidy; invasion then depends solely on the difference in long-run effective mutation rates. The *R*=0 invasion criterion can be evaluated analytically or by a single numerical integral for standard distributions; Appendix A and Table 2, summarize *E*[log(1 − *u*_2,*t*_)] and the relationship between *E*[log(1 − *u*_2,*t*_)] and log(1 − *u*_1_) for several cases. *Regime 2 (λ*_2_ *term dominates)*. If *E*[log(1 − *u*_2,*t*_)] ≤ log(1 − *u*_1_), then *γ*(0) = log *λ*_2_ *<* 0 and invasion fails. In this regime *γ*(0) is independent of *u*_2,*t*_ and its variance; only *u*_1_, *s*, and ploidy matter.

**Table 1:**
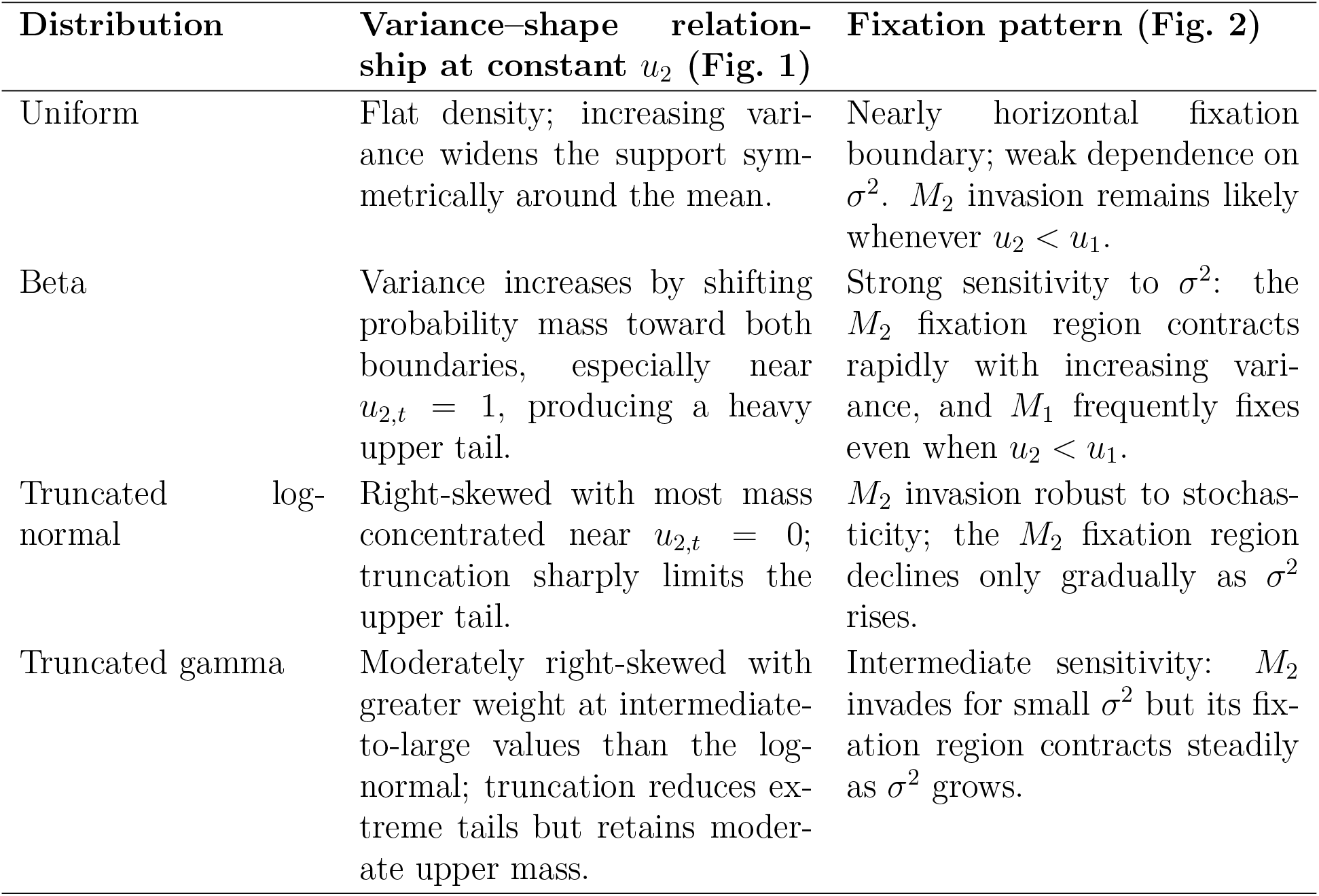
Summary of distributional effects on modifier invasion at *R* = 0. The shape of the mutation–rate distribution determines *E*[log(1 ™ *u*_2,*t*_)] and thus the long-term growth rate of the invading modifier. Distributions placing more probability mass near high mutation values (*u*_2,*t*_ → 1) yield smaller geometric means and therefore hinder invasion, even when the arithmetic mean *u*_2_ is held constant. Conversely, distributions concentrated at low *u*_2,*t*_ maintain larger *E*[log(1 − *u*_2,*t*_)] and favor invasion.

**Table 2:**
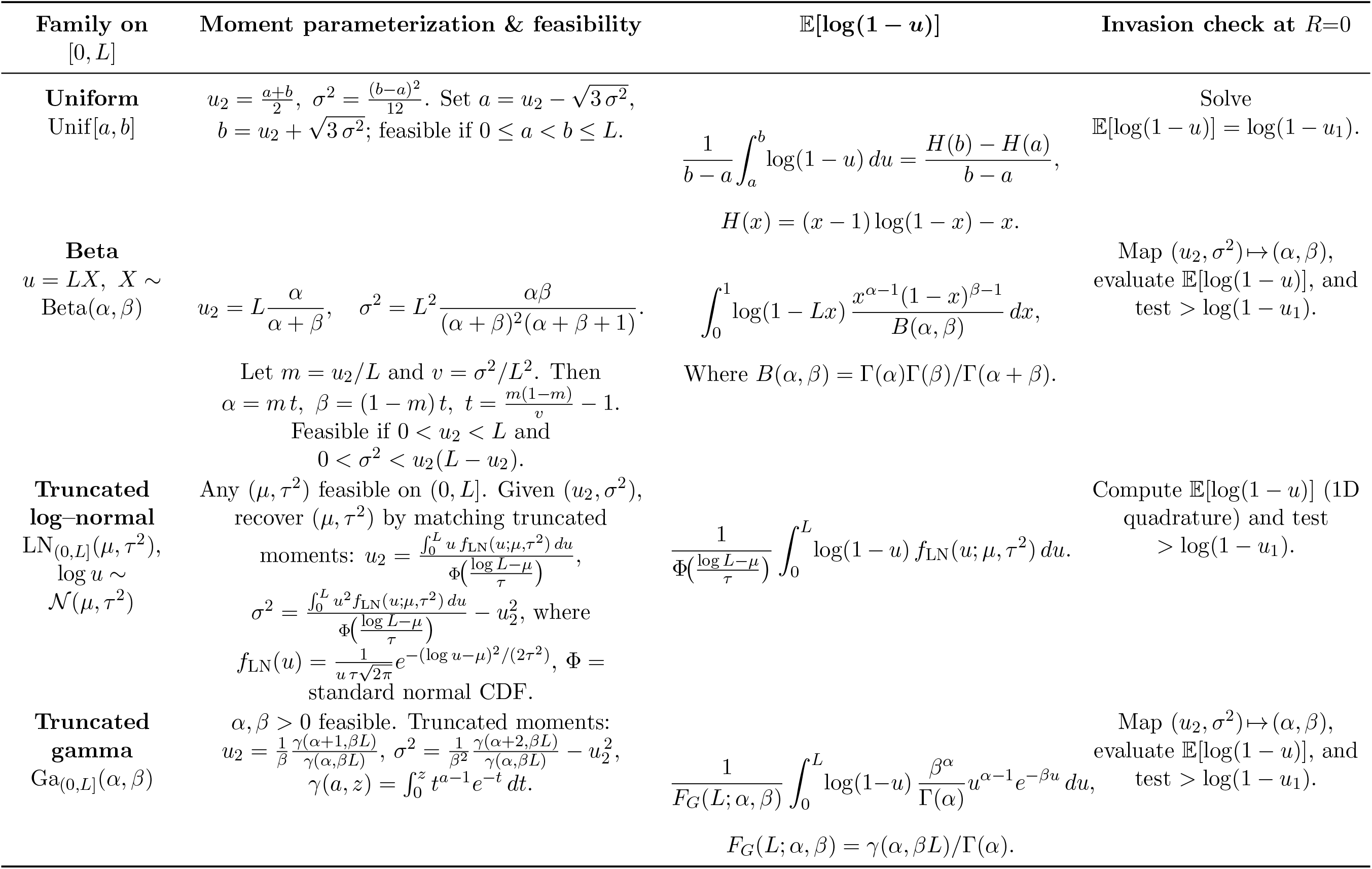
Summary of *E*[log(1 ™ *u*_2,*t*_)] and the invasion test *E*[log(1 ™ *u*_2,*t*_)] *>* log(1 ™ *u*_1_) at *R* = 0 for four distribution families on [0, *L*] (in simulations *L* = 1). Means and variances refer to the distribution on [0, *L*]. Closed forms are available for some special cases (e.g., Beta with *L*=1); otherwise a single one–dimensional quadrature suffices.

- *Haploids: λ*_2_ = (1 − *s*)*/*(1 − *u*_1_). For fixed *u*_1_, ∂*λ*_2_*/*∂*s* = −1*/*(1 − *u*_1_) *<* 0, so increasing *s* decreases *λ*_2_ and makes log *λ*_2_ more negative. For fixed *s <* 1, ∂*λ*_2_*/*∂*u*_1_ = (1 − *s*)*/*(1 − *u*_1_)^2^ *>* 0, so increasing *u*_1_ increases *λ*_2_ (moving log *λ*_2_ toward 0).
- *Diploids: λ*_2_ = *{*1 − *s*(1 + *u*_1_)*}/*(1 − *u*_1_), assumed positive (i.e., 1 − *s*(1 + *u*_1_) *>* 0) so that log *λ*_2_ is defined. For fixed *u*_1_, ∂*λ*_2_*/*∂*s* = −(1 + *u*_1_)*/*(1 − *u*_1_) *<* 0, so increasing *s* decreases *λ*_2_. Also,

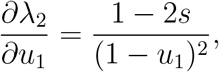 *λ*_2_ increases with *u*_1_ when 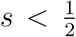 and decreases with *u*_1_ when 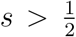, and is locally insensitive to *u*_1_ at 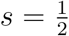 (subject to 1 − *s*(1 + *u*_1_) *>* 0). Accordingly, log *λ*_2_ becomes less negative or more negative in these cases.

In summary, when the *λ*_1_ term dominates, invasion is determined by differences in long-run effective mutation rates: it is promoted by higher *u*_1_ but hindered by temporal variance in *u*_2,*t*_. When the *λ*_2_ term dominates, invasion cannot occur, and the negative Lyapunov exponent *γ*(0) = log *λ*_2_ *<* 0 quantifies the strength of this constraint on the growth rate of *M*_2_’s frequency. Stronger selection decreases *λ*_2_, making *γ*(0) more negative and thus reinforcing the constraint. The effect of the resident mutation rate *u*_1_ is monotonic in haploids but depends on *s* in diploids, decreasing *λ*_2_ when 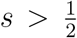 and increasing it when 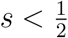.

#### Simulation Results

We examine how temporal variation in the mutation rate *u*_2_ alters invasion when its mean is held constant and *R* = 0. The rare modifier allele generates an i.i.d. sequence *u*_2,*t*_ ∼ *D*(*u*_2_, *σ*^2^) with mean *E*[*u*_2,*t*_] = *u*_2_ and variance Var(*u*_2,*t*_) = *σ*^2^, supported between 0 and 1. Four distributions are considered for *D*—uniform, beta, truncated log-normal, and truncated gamma—with parameters fitted to match (*u*_2_, *σ*^2^) exactly (Appendix A).

In the deterministic case (*u*_2,*t*_ ≡ *u*_2_), the modifier invades if *u*_2_ *< u*_1_. With stochastic variation in *u*_2_ and complete linkage (*R* = 0), the leading Lyapunov exponent satisfies

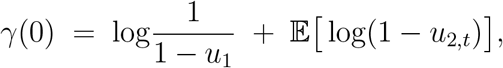

so invasion occurs if *E*[log(1 − *u*_2,*t*_)] *>* log(1 − *u*_1_) (Eq. (12)). Because log(1 − *x*) is strictly concave on [0, 1),

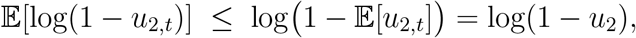

with strict inequality when *σ*^2^ *>* 0 (by Jensen’s inequality). Temporal variability in *u*_2,*t*_ therefore lowers *γ*(0) relative to the deterministic case, with the reduction driven mainly by infrequent large values of *u*_2,*t*_, which strongly affect the logarithmic mean. Figure 1 shows that, for a constant mean mutation rate *E*[*u*_2,*t*_] = *u*_2_ = 0.06, increasing the variance *σ*^2^ changes the shape of *u*_2,*t*_ under different distributional assumptions.

**Figure 1:**
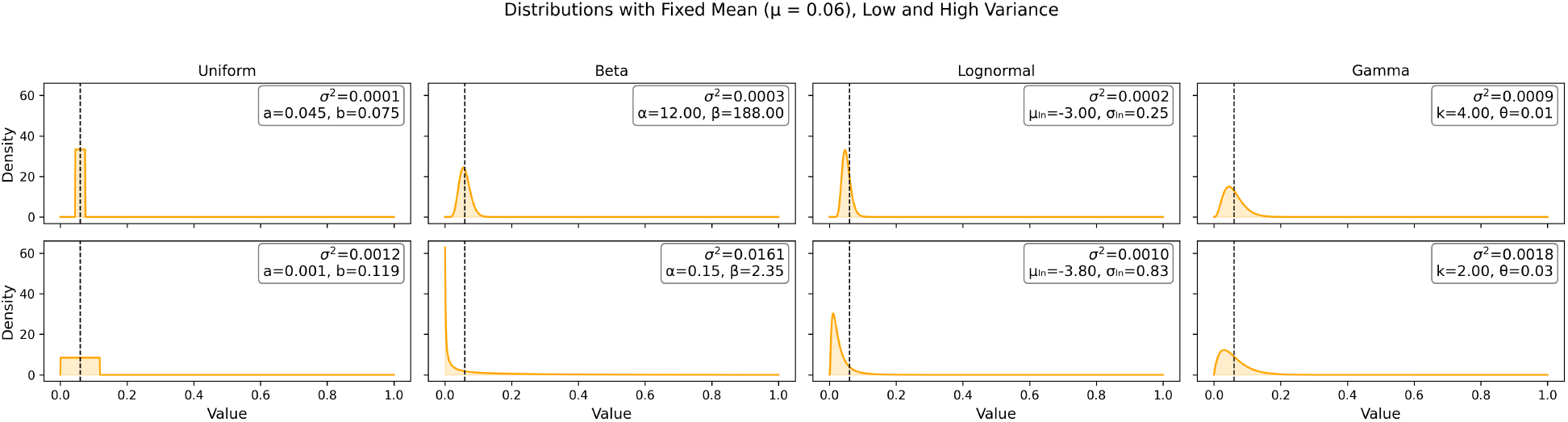
Mutation–rate probability densities on [0, 1] with fixed mean. *u*_2_ = 0.06. Columns show distributions (uniform, beta, truncated log-normal, truncated gamma); rows correspond to “low” (top) and “high” (bottom) variance parameterizations. Dashed vertical line: common mean. Parameters used: uniform [*a, b*] = [0.045, 0.075] (low), [0.001, 0.119] (high); beta (*α, β*) = (12.0, 188.0) (low), (0.15, 2.35) (high); log-normal (*µ*_ln_, *σ*_ln_) = (−3.0, 0.25) (low), (−3.8, 0.83) (high); gamma (*k, θ*) = (4.0, 0.015) (low), (2.0, 0.030) (high). Log-normal and gamma densities are truncated and renormalized on [0, 1]; realized variances (annotated in each panel) are computed numerically.

The distributions are parameterized to have the same mean but different higher moments. The uniform expands symmetrically with variance, while the beta shifts probability mass toward both boundaries—especially near *u*_2,*t*_ = 1—as *σ*^2^ increases. The truncated log-normal and gamma distributions are right-skewed; the former retains a relatively thin tail after truncation, whereas the latter has a heavier upper tail with comparable variance. These differences in shape determine how much weight is placed on infrequent, large *u*_2,*t*_ values that disproportionately reduce *E*[log(1 − *u*_2,*t*_)] making invasion less likely.

For each simulation, the population is initialized at the resident equilibrium under *M*_1_ (haploid example: *u*_1_ ≈ 0.048796), set *s* = 0.2 and *R* = 0, and the full two-locus recursion is iterated for 5,000 generations. Fixation is declared if (*x*_3_ + *x*_4_)_*t*_ ≥ 0.999 (modifier *M*_2_) or (*x*_1_ + *x*_2_)_*t*_ ≥ 0.999 (resident *M*_1_); otherwise the outcome is “no fixation.” For each distributional family *D* we generate an i.i.d. sequence *u*_2,*t*_ ∼ *D*(*u*_2_, *σ*^2^) with prescribed mean *u*_2_ and variance *σ*^2^.

Parameterization is as follows. (i) *uniform:* choose [*a, b*] with 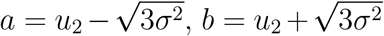, and *σ*^2^ = (*b* − *a*)^2^*/*12. (ii) *beta:* set *α* = *u*_2_*m, β* = (1 − *u*_2_)*m* with *m* = *u*_2_(1 − *u*_2_)*/σ*^2^ − 1 (feasible for *σ*^2^ *< u*_2_(1−*u*_2_)); this matches (*u*_2_, *σ*^2^) exactly on [0, 1]. (iii) *truncated log-normal* and (iv) *truncated gamma:* choose (*µ*_ln_, *σ*_ln_) or (*k, θ*) so that the [0, 1]-truncated density has mean *u*_2_ and variance *σ*^2^; parameters are obtained by a two-dimensional root-finding routine, and the realized *σ*^2^ is computed by numerical quadrature. The feasible variance range [min *σ*^2^, max *σ*^2^] depends on the family: beta admits the largest *σ*^2^ (probability near boundaries), uniform is limited by interval width, and truncation restricts log-normal and gamma to have moderate *σ*^2^.

Figure 2 summarizes simulation outcomes in terms of (*u*_2_, *σ*^2^) for each *D*. Boundaries align with the geometric-mean criterion at *R* = 0: distributions placing little mass at high mutation (uniform; truncated log-normal) maintain larger *E*[log(1 − *u*_2,*t*_)] and favor invasion, whereas distributions allowing frequent large *u*_2,*t*_ (beta; high-variance truncated gamma) reduce the logarithmic mean and disfavor invasion—even when the arithmetic mean *u*_2_ *< u*_1_. Quantitative comparisons across distributions are provided in Appendix A.1, Table 3, which reports how the invasion boundary 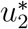 varies with constant variance for different distributions. These results illustrate how the location of probability mass relative to *u*_1_ affects invasion through its effect on *E*[log(1 − *u*_2,*t*_)].

**Table 3:** Largest mean 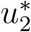 (at fixed *σ*^2^) consistent with *γ* = 0, together with the *u*_1_ cdf and upper–tail penalty.

| Distribution on $[0, 1]$ | $u_2^*$ | $\mathbf{C}(u_1)$ | $\mathcal{T}(u_1)$ |
| --- | --- | --- | --- |
| Uniform | 0.09427879 | 0.507983 | 0.068280 |
| Beta | 0.09448161 | 0.559617 | 0.061754 |
| Truncated log-normal | 0.09432226 | 0.591052 | 0.057932 |
| Truncated gamma | 0.09427113 | 0.567150 | 0.060784 |

**Figure 2:**
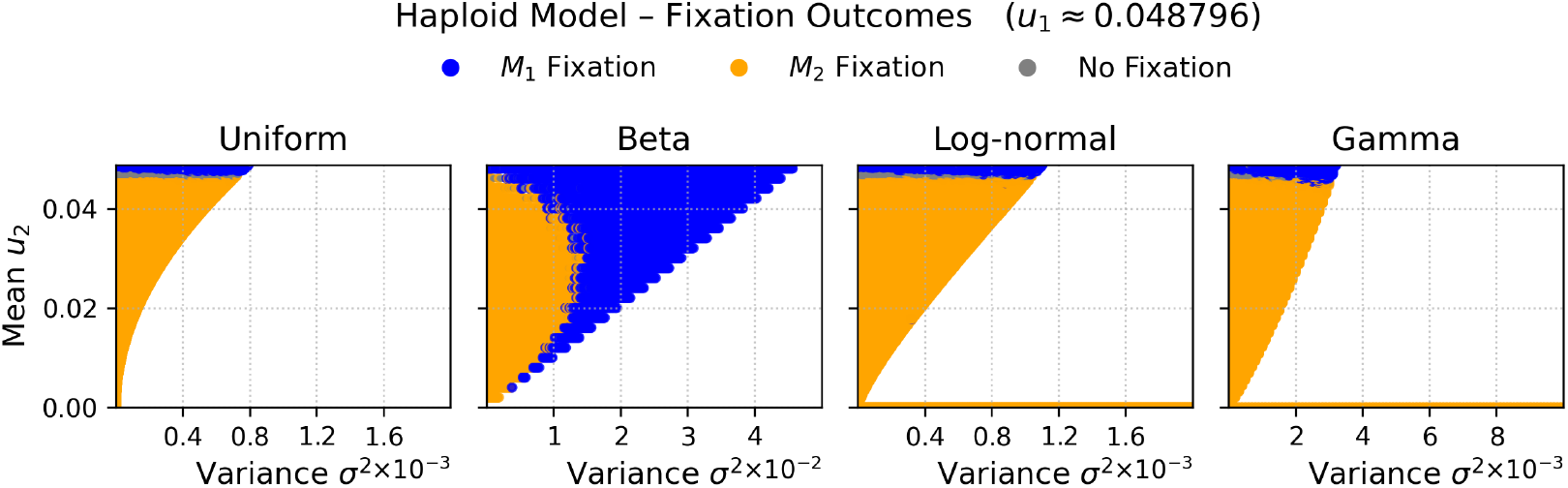
Fixation outcomes for alleles *M*_1_ and *M*_2_ across (*u*_2_, *σ*^2^) at *R* = 0 (haploids). Settings: 5,000 generations; *M*_2_ initially rare; resident *M*_1_ fixed at *u*_1_ = 0.048796; *s* = 0.2. Columns correspond to distributional families (uniform, beta, truncated log-normal, truncated gamma). The variance axis in each panel reflects the feasible range at constant *u*_2_: uniform *σ*^2^ ∈ [0, 2 *×* 10^−3^], beta [0, 5 *×* 10^−2^], truncated log-normal [0, 2 *×* 10^−3^], truncated gamma [0, 10^−2^] (panel limits used in the simulations). Parameterization for each sampled point: uniform 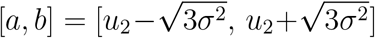 clipped to [0, 1]; beta (*α, β*) = (*u*_2_*m*, (1−*u*_2_)*m*) with *m* = *u*_2_(1 ™ *u*_2_)*/σ*^2^ ™ 1; truncated log-normal and truncated gamma fitted so their truncated moments equal (*u*_2_, *σ*^2^) (realized *σ*^2^ obtained by numerical quadrature). Colors denote outcomes: *M*_1_ fixation (blue), *M*_2_ fixation (orange), no fixation within 5,000 generations (gray).

These results highlight two key insights. First, invasion depends on *E*[log(1 − *u*_2,*t*_)] rather than on *E*[*u*_2,*t*_]. For a constant mean of *u*_2,*t*_, greater variance *σ*^2^ decreases *E*[log(1 − *u*_2,*t*_)]. More generally, when two distributions with the same mean are ordered by convexity (i.e., one is more variable in the sense of second-order stochastic dominance), the expectation of the concave function log(1 − *x*) is smaller for the more variable distribution. Consequently, modifier alleles with identical mean mutation rates may differ in their invasion outcomes solely due to differences in temporal variance. That is, if 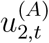 and 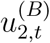 are two i.i.d. sequences with 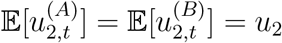 and

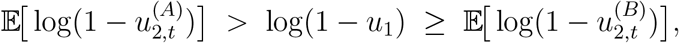

then 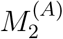 invades at *R* = 0 while 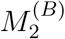 does not, despite identical means.

#### Stochastic mutation rate with Recombination 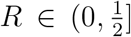

For *R >* 0, **F**_*t*_(*R*) has positive off–diagonal entries whenever *u*_2,*t*_ *>* 0; products of **F**_*t*_(*R*) do not commute, and no closed form for *γ*(*R*) is available. However, the qualitative effect of *R* derives from the linear structure of (5a) and (6a): recombination has no direct fitness effect at the modifier alleles, it mediates the indirect response by redistributing the associations between *M*_2_ and the selected background that are generated each generation by selection and mutation. In deterministic settings (constant *u*_2_), this association brings the dominant eigenvalue of the linearized near 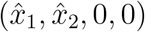 towards 1, changing the *magnitude* but not the *sign* of initial frequency change. With stochastic *u*_2,*t*_, multiplicative averaging emphasizes the geometric mean of per–generation growth factors, and the distribution of *u*_2,*t*_ interacts with *s* and *u*_1_, allowing recombination to *reverse* the sign of *γ*(*R*) as *R* increases—an effect not seen in deterministic models.

#### Numerical analysis

To assess the effect of recombination with stochastically varying mutation induced by *M*, let 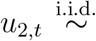 Beta(*α, β*) with mean *u*_2_ = 0.04 and variance *σ*^2^ ∈ *{*0, 0.25, 0.5, 0.75, 0.95*} ×u*_2_(1 − *u*_2_), where *u*_2_(1 − *u*_2_) = 0.0384. The factor *u*_2_(1 − *u*_2_) is the maximum possible variance for a variable on [0, 1] with mean *u*_2_ and scaling by *u*_2_(1 − *u*_2_) expresses *σ*^2^ as a fraction of this upper bound for comparability. For each *σ*^2^, we draw one sequence *{u*_2,*t*_*}*, and reuse it for both ploidies to isolate ploidy effects. For *R* on a grid in 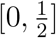, we iterate ***v***_*t*+1_ = **F**_*t*_(*R*) ***v***_*t*_ with ℓ^1^-normalization at each step; by the Furstenberg–Kesten theorem, the time average of the log rescalings converges almost surely to the top Lyapunov exponent *γ*(*R*) [13]. Unless stated otherwise, *u*_1_ = 0.05 and *s* ∈ *{*0.20, 0.06*}*. Figure 3 plots *γ*(*R*) for both ploidies; the left and right endpoints of each curve correspond to *R* = 0 and 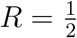, respectively.

**Figure 3:**
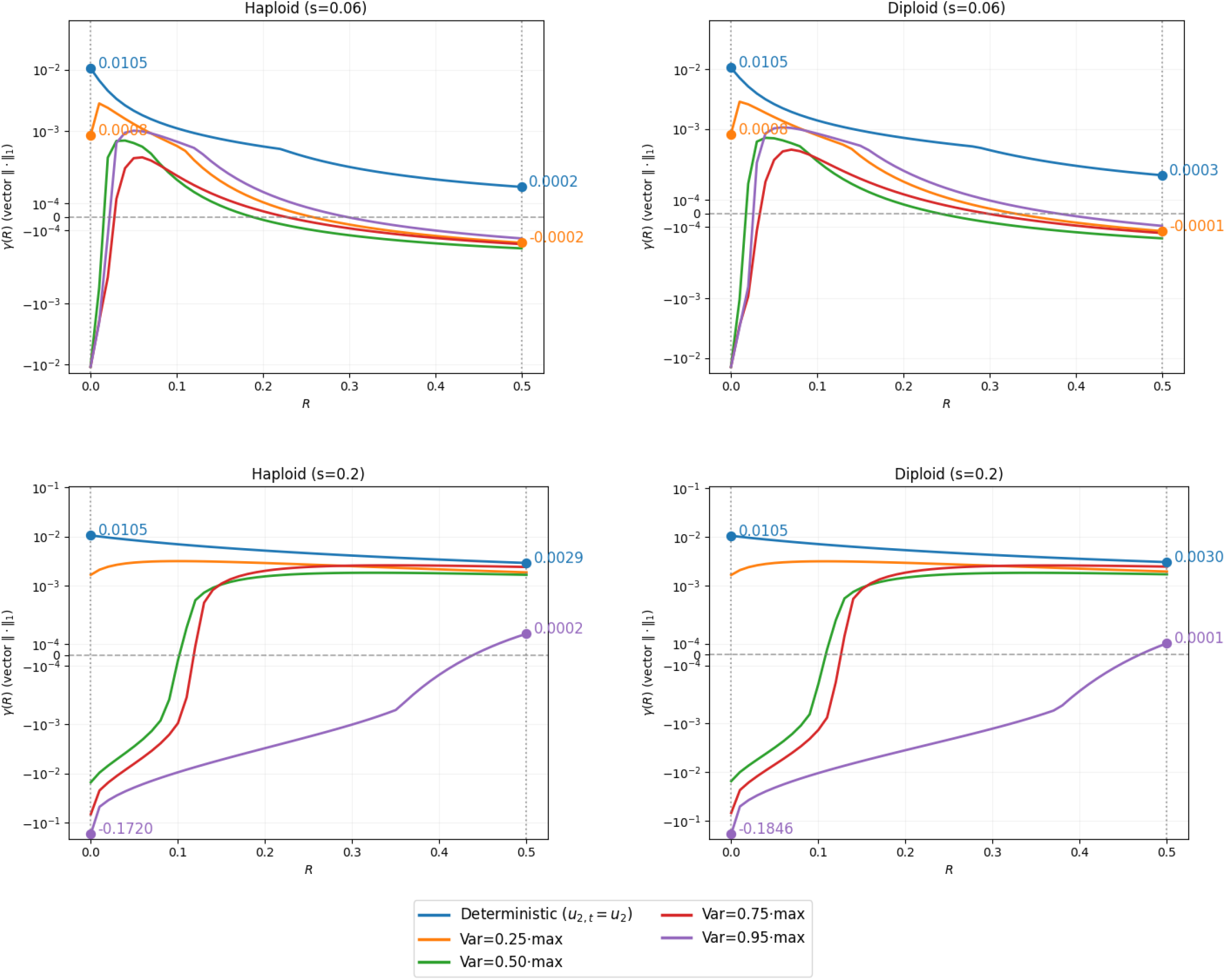
Top Lyapunov exponent *γ*(*R*) for the two-locus modifier system. Panels are arranged in a 2 × 2 grid: rows indicate selection strength *s* ∈ {0.06, 0.20}; columns indicate ploidy (haploid vs. diploid). For each (*s*, ploidy), curves correspond to variances line (*σ*^2^ = 0, labeled *u*_2,*t*_ = *u*_2_) uses *γ*(*R*) = log *ρ* **F**(*u*_2_, *R*). Stochastic curves plot the *σ*^2^ ∈ *{*0, 0.25, 0.50, 0.75, 0.95*} × u*_2_(1 − *u*_2_) with *u*_2_ = 0. 04. The deterministic base-sample Lyapunov exponent from normalized matrix products using the vector ℓ^1^ norm at each iteration (i.e., ∥ · ∥_1_ re-scaling). Annotated points display *γ*(0) and 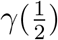 only for selected variance levels: in the weak-selection panels (*s* = 0.06), annotations are shown for *σ*^2^ = 0 and *σ*^2^ = 0.25 *u*_2_(1 − *u*_2_); in the strong-selection panels (*s* = 0.20), for *σ*^2^ = 0 and *σ*^2^ = 0.95 *u*_2_(1 − *u*_2_).

- *Weak selection (s* = 0.06). With low variance (0.25 *× u*_2_(1 − *u*_2_)), *γ*(0) *>* 0 while 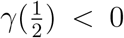 (e.g., haploid: 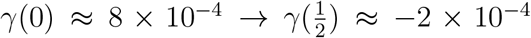; diploid: 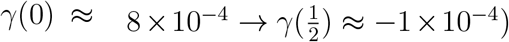). Thus if recombination is frequent enough, it suppresses invasion.
- *Strong selection (s* = 0.20). With high variance (0.95 *× u*_2_(1 − *u*_2_)), *γ*(0) *<* 0 while 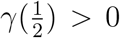 in both ploidies (e.g., haploid: 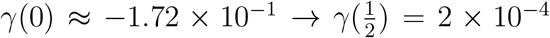; diploid: 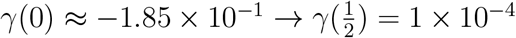). Thus sufficiently frequent recombination allows invasion.

Fig. 3 shows that sign changes in *γ*(*R*) with *R* occur only when variability in *u*_2,*t*_ is present; with *σ*^2^ = 0 (deterministic *u*_2,*t*_ ≡ *u*_2_), *R* changes the magnitude but not the sign of *γ*(*R*) on 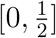.

##### Recombination cutoff

Define the cutoff 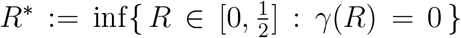, the smallest recombination fraction at which the long–run growth rate of *M*_2_ changes invasion direction (*γ*(*R*) ≶ 0). This quantity identifies the minimum recombination needed to reverse the direction of invasion for a given parameter set. We estimate *R*^*^ by evaluating *γ*(*R*) on a fine grid in 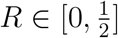 using a common realization of *{u*_2,*t*_*}* for all *R* (variance reduction via common random numbers). We then bracket a sign change, selecting an interval [*R*_*L*_, *R*_*U*_] with *γ*(*R*_*L*_)*γ*(*R*_*U*_) *<* 0, and solve *γ*(*R*) = 0 on [*R*_*L*_, *R*_*U*_] by a one–dimensional root finder (bisection or Brent). Tolerances are |*γ*(*R*^*^)| *<* 10^−5^ or interval width *<* 10^−3^. Results are stable to numerical error across independent replicates to numerical error.

- *Weak selection example*. For *s* = 0.06 and a beta distribution for *u*_2,*t*_ calibrated to mean *u*_2_ and variance *σ*^2^ = 0.25 *× u*_2_(1 − *u*_2_), we obtai

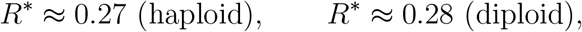

consistent with the zero crossing near *R* ∈ [0.25, 0.30] in Fig. 3. In this setting, moderate recombination reverses the sign of *γ*(*R*) because it disrupts associations between *M*_2_ and high–mutation backgrounds that disproportionately reduces the geometric mean growth rate. When the variance is increased (e.g., 0.50*×u*_2_(1−*u*_2_) and 0.75*×u*_2_(1−*u*_2_) for the same beta distribution), *γ*(*R*) can become non-monotone in *R*. Intuitively, for small *R*, recombination primarily breaks associations formed in generations with larger *u*_2,*t*_, raising *γ*(*R*); at higher *R*, it also breaks beneficial associations formed during low–*u*_2,*t*_ generations, lowering *γ*(*R*). This can yield two zero crossings in *R*; by definition, *R*^*^ denotes the smaller root. The presence and locations of these roots depend on (*s, u*_1_) and the full distribution of *u*_2,*t*_ (not just its mean and variance).
- *Strong selection example*. For *s* = 0.20 and high variance *σ*^2^ = 0.95 *× u*_2_(1 − *u*_2_),

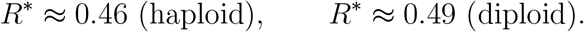 Here, reversal requires recombination to be close to free 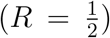: strong selection amplifies the cost of low-fitness associations, so more frequent reshuffling is needed to offset them. In these high-variance cases, *γ*(*R*) typically increases with *R* for small *R* (breaking high–*u*_2,*t*_ associations) and may decrease for larger *R* once the loss of favorable associations during low–*u*_2,*t*_ episodes dominates.

The above variance multipliers were chosen because, among the tested set *{*0, 0.25, 0.5, 0.75, 0.95*}*, they were the lowest and highest values for which *γ*(*R*) changed sign under weak and strong selection, respectively.

##### Parameter effects

For *R >* 0, recombination affects associations between the modifier and the selected background without directly changing modifier fitness. Consequently, the sensitivity of *γ*(*R*) to *R* is governed by (i) how selection strength *s* and the resident mutation rate *u*_1_ are associated at each generation, and (ii) how temporal variability in *u*_2,*t*_ enters through geometric (multiplicative) averaging. Throughout, we use the linear decomposition **F**_*t*_(*R*) = **A**_*t*_ + *R* **B**_*t*_ with

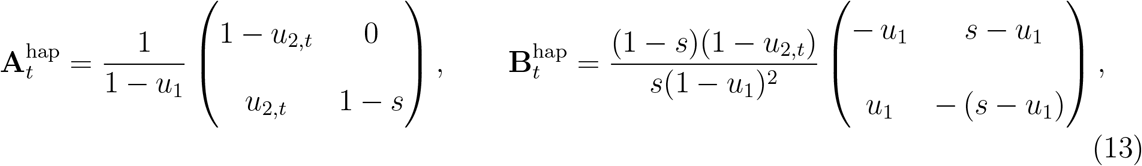

for haploids, and

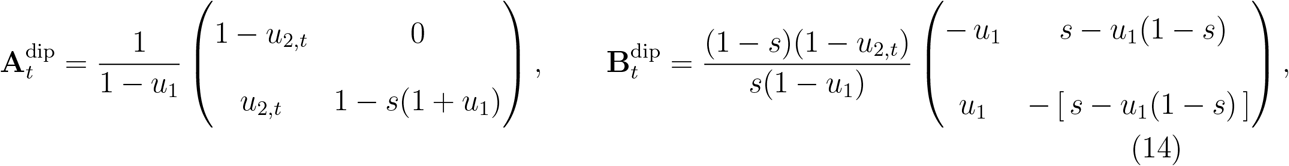

for diploids. These make explicit that *R* enters *linearly* in each generation via **B**_*t*_. However, *γ*(*R*) itself is not generally a linear (or monotone) function of *R* because it depends on products of non-commuting random matrices.

- *Selection strength s*. In both ploidies, **B**_*t*_ scales as (1 − *s*)*/s* times a matrix whose off–diagonal elements increase linearly with *s* (specifically *s* − *u*_1_ in haploids and *s* − *u*_1_(1 − *s*) in diploids). Because d[(1 − *s*)*/s*]*/*d*s* = −1*/s*^2^ *<* 0, increasing *s* reduces the overall multiplicative factor governing the per–generation sensitivity to *R*, while simultaneously amplifying certain matrix entries that mediate background exchange. The resulting effect of *s* on the long–run growth rate *γ*(*R*) is therefore non-monotonic: stronger selection diminishes the effect of recombination yet enhances direct coupling between backgrounds. At *R* = 0, where **F**_*t*_ = **A**_*t*_ is lower–triangular, the eigenvalue *λ*_2_ equals 1 − *s* (haploid) or 1 − *s*(1 + *u*_1_) (diploid), with ∂*λ*_2_*/*∂*s <* 0, so log *λ*_2_ decreases strictly with *s*. For *R >* 0, however, *γ*(*R*) depends on products of non-commuting random matrices, and its response to increasing *s* cannot be determined in general.
- *Resident mutation rate u*_1_. Assume *u*_1_ ∈ [0, 1) so that expectations of logs exist. At *R* = 0, the product is lower–triangular and *γ*(0) = *E*[log(1 − *u*_2,*t*_)] − log(1 − *u*_1_), hence

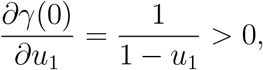

so increasing *u*_1_ increases *γ*(0). For *R >* 0, *u*_1_ enters **B**_*t*_ through global factors (1−*u*_1_)^−2^ (haploid) or (1 − *u*_1_)^−1^ (diploid), which amplify recombination’s per–generation effect as *u*_1_ increases, and through off–diagonals *s* − *u*_1_ (haploid) and *s* − *u*_1_(1 − *s*) (diploid), which *decrease* with *u*_1_; these opposing influences, combined with temporal covariances between *u*_2,*t*_ and selection–generated modifier–background associations, entail that the effect of *u*_1_ on *γ*(*R*) for *R >* 0 has no general sign and is parameter–dependent.
- *Temporal variability of u*_2,*t*_. For constant mean *u*_2_, concavity of log(1 − *x*) on [0, 1) gives *E*[log(1 − *u*_2,*t*_)] ≤ log 1 − *E*[*u*_2,*t*_], with strict inequality when Var(*u*_2,*t*_) *>* 0. Thus higher variance lowers the *R*=0 component of *γ*(*R*). When *R >* 0, *u*_2,*t*_ enters **B**_*t*_ through the factor (1 − *u*_2,*t*_), so higher temporal variance increases fluctuations in the per–generation growth rate and modifies how recombination influences the geometric mean. The overall influence of variability on *γ*(*R*) depends on the joint dynamics of *u*_2,*t*_ and the modifier–background associations and cannot be determined in general.
- *Comparing ploidies*. The recombination prefactor is (1−*u*_1_)^−2^ in haploids and (1−*u*_1_)^−1^ in diploids, so—holding other terms fixed—the single–step *R*–perturbation is larger in haploids. However, the *s*–dependent off–diagonal terms also differ (e.g., *s* − *u*_1_ vs. *s* − *u*_1_(1 − *s*)), and *λ*_2_ differs between ploidies at *R* = 0. Consequently, there is no uniform ordering by ploidy; differences in *γ*(*R*) arise from the interaction of these factors with the distribution of *u*_2,*t*_.

Stochastic transmission replaces arithmetic by geometric averaging of per–generation growth factors, so invasion depends on the *full distribution* of *u*_2,*t*_ rather than only its mean. At *R* = 0, the condition *γ*(0) *>* 0 is equivalent to *E*[log(1 − *u*_2,*t*_)] *>* log(1 − *u*_1_); by concavity of log(1 − *x*), greater variance in *u*_2,*t*_ strictly lowers *E*[log(1 − *u*_2,*t*_)] at constant mean, reducing *γ*(0). For *R >* 0, recombination has no direct fitness effect at the modifier, but it affects the time–varying associations between *M*_1_*/M*_1_ and *A/a* generated each generation by selection and mutation. Because *γ*(*R*) is a Lyapunov exponent of products of random matrices, its dependence on *R* need not be linear or monotone: recombination can increase *γ*(*R*) when it disproportionately breaks associations formed during high–*u*_2,*t*_ generations, and decrease *γ*(*R*) when it breaks associations formed during low–*u*_2,*t*_ generations. The joint effects of the distribution of *u*_2,*t*_ (particularly its variance), the selection coefficient *s*, and the resident mutation rate *u*_1_ can therefore induce sign reversals of *γ*(*R*) that are absent in deterministic models with constant *u*_2_. In sum, stochastic mutation at the modifier creates regimes in which recombination alters not only the *magnitude* of the growth rate but also the *direction* of invasion.

## Discussion

Most analyses of genetic evolution in large populations introduce stochasticity through fluctuating selection while holding transmission parameters fixed. In this study, selection is constant and randomness enters through the transmission process itself. A neutral modifier locus (*M*_1_*/M*_2_) changes the forward mutation rate at a linked, selected locus (*A/a*): the resident *M*_1_ has constant rate *u*_1_, whereas the invader *M*_2_ has a generation–to–generation rate *u*_2,*t*_ ∈ [0, 1) with mean *u*_2_ and variance *σ*^2^. We ask how *σ*^2^ *>* 0 alters *M*_2_ invasion. In the deterministic setting (*u*_2,*t*_ ≡ *u*_2_), both haploid and diploid models yield an invasion condition that is independent of *R*: the modifier *M*_2_ increases when rare if *u*_2_ *< u*_1_, consistent with the *Reduction Principle* (Eq. (7)). Recombination shifts the dominant eigenvalue of the linearized system toward unity, reducing the magnitude of indirect selection without altering its sign; hence, recombination affects the rate but not the direction of invasion. In this case, the direction of modifier evolution is statistically aligned with the equilibrium structure of the resident system.

With stochastic mutation (Var(*u*_2,*t*_) *>* 0) and complete linkage (*R* = 0), the invasion recursion is lower triangular and the criterion becomes geometric (Eq. (12)):

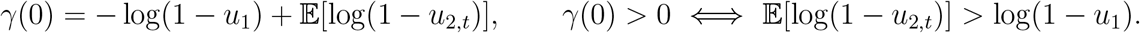

Because log(1 − *x*) is strictly concave on [0, 1), *E*[log(1 − *u*_2,*t*_)] ≤ log(1 − *u*_2_), with strict inequality when *σ*^2^ *>* 0; randomness in *u*_2,*t*_ therefore reduces *γ*(0) relative to the deterministic case. Occasional large *u*_2,*t*_ values near 1 can substantially reduce *E*[log(1−*u*_2,*t*_)]. Simulations at *R* = 0 with uniform, beta, truncated log–normal, and truncated gamma families support this: with constant *u*_2_, heavier upper tails yield smaller geometric means and lower invasion probabilities.

Variability at *R* = 0 also modifies the deterministic *Reduction Principle*. When *u*_2,*t*_ ≡ *u*_2_, any reduction (*u*_2_ *< u*_1_) gives *γ*(0) = log {(1 − *u*_2_)*/*(1 − *u*_1_)) *>* 0. If *u*_2,*t*_ fluctuates with small variance *σ*^2^, a Taylor expansion yields

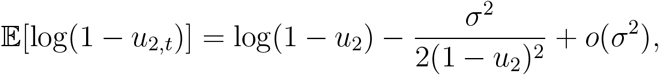

so invasion requires

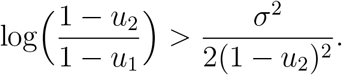

Thus, for sufficiently large *σ*^2^, invasion can fail even when *E*[*u*_2,*t*_] *< u*_1_. Variability in transmission can therefore reduce—or overturn—the deterministic *Reduction Principle*. For 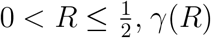 is continuous in *R* under the regularity conditions assumed for the product of random matrices. At *R* = 0, the geometric–mean condition defines the baseline, with *γ*(0) decreasing monotonically in Var(*u*_2,*t*_) with constant mean. When *R >* 0, recombination reshuffles haplotypes, averaging fitness effects over the distribution of associations produced each generation by selection and mutation. This averaging can either increase or decrease the Lyapunov exponent: *γ*(*R*) increases when recombination disproportionately breaks associations formed during high–*u*_2,*t*_ generations (which decreases multiplicative growth), and decreases when it breaks associations formed during low–*u*_2,*t*_ generations. Consequently, with Var(*u*_2,*t*_) *>* 0 the direction of modifier evolution is not necessarily aligned with the deterministic equilibrium and can be reversed by recombination —a phenomenon absent when *u*_2,*t*_ is deterministic. The direction of this stochastic reversal depends on *u*_1_, *s*, ploidy, and the distribution of *u*_2,*t*_; modifier alleles with the same mean *u*_2_ can therefore differ in invasion solely because temporal variability in *u*_2,*t*_ induces different responses to recombination. Two conclusions follow. First, the *Reduction Principle* emerges as the deterministic limit: with constant *u*_2_, the invasion condition depends only on the ordering of *u*_2_ and *u*_1_; under stochastic *u*_2,*t*_, the relevant quantity is the geometric mean of (1 − *u*_2,*t*_), which can diverge markedly from predictions based on *u*_2_. Second, recombination neither uniformly facilitates nor uniformly hinders invasion; its effect depends jointly on *u*_1_, *s*, ploidy, and the distribution of *u*_2,*t*_. Mean mutation rates alone are therefore insufficient to predict outcomes: modifiers alleles with identical *u*_2_ can differ in invasion outcome purely because of differences in temporal variance or tail behavior, through their impact on E[log(1 − *u*_2,*t*_)] and on products of the *R*–dependent **F**_*t*_ matrices.

Several extensions follow naturally. Introducing weak temporal correlation in *u*_2,*t*_ might affect whether clustering of high–mutation generations changes invasion probabilities and how this interacts with small *R >* 0. Allowing recombination itself to vary randomly 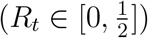, possibly jointly with *u*_2,*t*_, would generalize the framework to products of matrices with time–varying *R*_*t*_; the effect on *γ* would depend on the joint distribution of (*u*_2,*t*_, *R*_*t*_) and their covariation with selection–generated associations. *Empirically*, these results motivate estimating not only *E*[*u*_2,*t*_] and Var(*u*_2,*t*_) but also *E*[log(1 − *u*_2,*t*_)] and the temporal covariance between *u*_2,*t*_ and measures of linkage or background fitness, using pedigree–based recombination maps, mutation–accumulation lines, or experimental evolution data.

In summary, stochastic transmission replaces the arithmetic–mean criterion of deterministic models with a geometric–mean criterion. Rare but extreme realizations of *u*_2,*t*_ disproportionately reduce the expected logarithmic growth rate, while recombination determines how this variability is averaged across genetic backgrounds. Invasion therefore depends jointly on the distribution of *u*_2,*t*_, the recombination rate, selection strength, and ploidy. Stochastic variation in *u*_2,*t*_ can reverse the deterministic *Reduction Principle*, allowing recombination to change not only the *magnitude* but also the *direction* of modifier invasion.

## Supporting information

Full Project Package

## A Estimation of Invasion Regions from Lyapunov Exponents: Examples

With *R* = 0, the invasion matrices are lower triangular for both ploidies, so the top Lyapunov exponent equals the larger of the two diagonal log–growth rates. Fix *u*_1_ ∈ (0, 1) and a rare modifier allele *M*_2_ with i.i.d. mutation rates *u*_2,*t*_ ∈ [0, *L*] (*L* ≤ 1), mean *u*_2_ and variance *σ*^2^.

Then

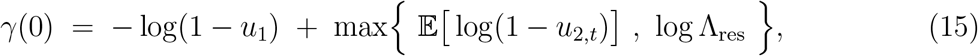

where

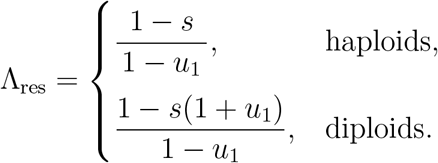

Under the resident–equilibrium constraints (*u*_1_ *< s* in haploids; *u*_1_ *< s/*(1 − *s*) in diploids), log Λ_res_ *<* log(1 − *u*_1_), and (15) yields the invasion criterion

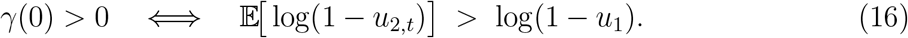

For any distribution *D* on [0, *L*] with density *f* (*u*), the expected log term is

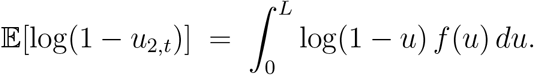

This quantity can be evaluated directly once the distribution is parameterized by its mean *u*_2_ and variance *σ*^2^, either in closed form or by one–dimensional numerical integration. The invasion boundary then follows from solving

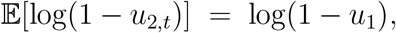

subject to the feasible support of *D*. Table 2 summarizes analytical and numerical expressions for *E*[log(1 − *u*_2,*t*_)] and the corresponding invasion criterion across the four distribution families considered.

### A.1 Sensitivity of *E*[log(1 − *u*_2,*t*_)] to Distributional Shape

Fix the variance *σ*^2^ ≈ 0.00159, support (0, 1), and resident mutation rate *u*_1_ ≈ 0.0954. For each distribution family, let 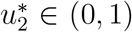 denote the *largest* mean such that, under complete linkage (*R* = 0),

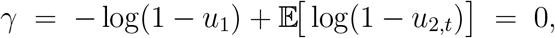

so that 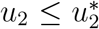 implies invasion at *R* = 0. The function log(1 − *u*_2,*t*_) is strictly decreasing and strictly concave on (0, 1), with derivatives (1 − *u*_2,*t*_)^−1^, (1 − *u*_2,*t*_)^−2^, and (1 − *u*_2,*t*_)^−3^ increasing as *u*_2,*t*_ ↑ 1. Consequently, *E*[log(1 − *u*_2,*t*_)] is especially sensitive to probability mass near the upper boundary. To quantify how probability mass relative to *u*_1_ influences *E*[log(1 − *u*_2,*t*_)], define

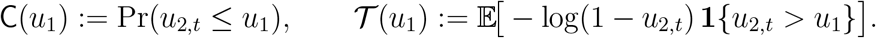

Here, **1***{·}* denotes the indicator function, equal to 1 when the condition inside the bracket is true and 0 otherwise. The quantity *T*(*u*_1_) measures the expected contribution to − log(1 − *u*_2,*t*_) from realizations where *u*_2,*t*_ *> u*_1_, thus capturing the influence of the right tail of the mutation–rate distribution on the overall mean. Larger C(*u*_1_) (greater probability mass at or below *u*_1_) increases E[log(1 − *u*_2,*t*_)], whereas larger *T* (*u*_1_) (heavier right tail above *u*_1_) decreases it. For the fixed *σ*^2^ and *u*_1_ given above, Table 3 reports 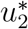 together with C(*u*_1_) and *T*(*u*_1_) for the four distribution families. The 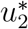 ordering in Table 3 (beta *>* truncated log–normal *>* uniform *>* truncated gamma) follows from the combined effect of C(*u*_1_) and *T*(*u*_1_) together with where mass lies below *u*_1_. The beta case has the largest 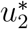 by pairing relatively high C with moderate *T*. The truncated log–normal has the highest C and the lowest *T* but yields a slightly smaller 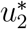, reflecting diminishing returns from placing additional probability far below *u*_1_. The uniform allocates more mass above *u*_1_, raising *T* and lowering 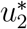. The truncated gamma places comparatively more mass in the upper mid-range than the log–normal, increasing *T* enough to produce the smallest 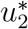. Although the numerical differences are small (on the order of 2 *×* 10^−4^), they are systematic: for fixed variance, shifting probability from the right tail toward (and just below) *u*_1_ increases *E*[log(1 − *u*_2,*t*_)] and thus permits a larger 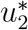, while shifting probability toward the boundary *u* ≈ 1 has the opposite effect.

### A.2 Accuracy of Lyapunov Approximations

We compare the Lyapunov invasion criterion with stochastic recursions (5,000 generations) for four mutation–rate distributions on [0, *L*] (uniform, beta, truncated log–normal, truncated gamma). With resident *u*_1_ ≈ 0.048796 and fixed variance *σ*^2^, the *R*=0 invasion boundary is the mean 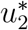 solving

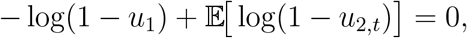

where *u*_2,*t*_ is drawn from the chosen distribution fitted to *E* [*u*_2,*t*_] = *u*_2_ and Var(*u*_2,*t*_) = *σ*^2^ (closed form fitting for uniform and beta; moment inversion for truncated distributions). Because log(1 − *x*) is strictly decreasing on [0, 1), the left-hand side is strictly decreasing in *u*_2_, so any root 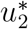 is unique. Numerically, we restrict *u*_2_ to the feasible mean range of each distribution and solve by Brent’s method (tolerance 10^−10^). Expectations are evaluated exactly for the uniform case and, when available, in closed form for beta on [0, 1]; otherwise we use one–dimensional adaptive quadrature with an endpoint split near *L* = 1 to handle the logarithmic singularity.

Figure 4 shows close agreement between simulated fixation outcomes and the Lyapunov threshold. The prediction is slightly conservative: *u*_2_ values below the curve lead to reliable invasion. Differences across distributions arise from how fixed variance reallocates probability mass relative to *u*_1_ and toward the upper boundary 1, where log(1 − *u*_2,*t*_) is highly sensitive. When most probability lies below *u*_1_ and away from 1 (e.g., truncated log–normal with a thin upper tail, or uniform once its interval includes *u*_1_), *E*[log(1 − *u*_2,*t*_)] changes little with *σ*^2^, yielding an almost horizontal boundary in (*u*_2_, *σ*^2^).

**Figure 4:**
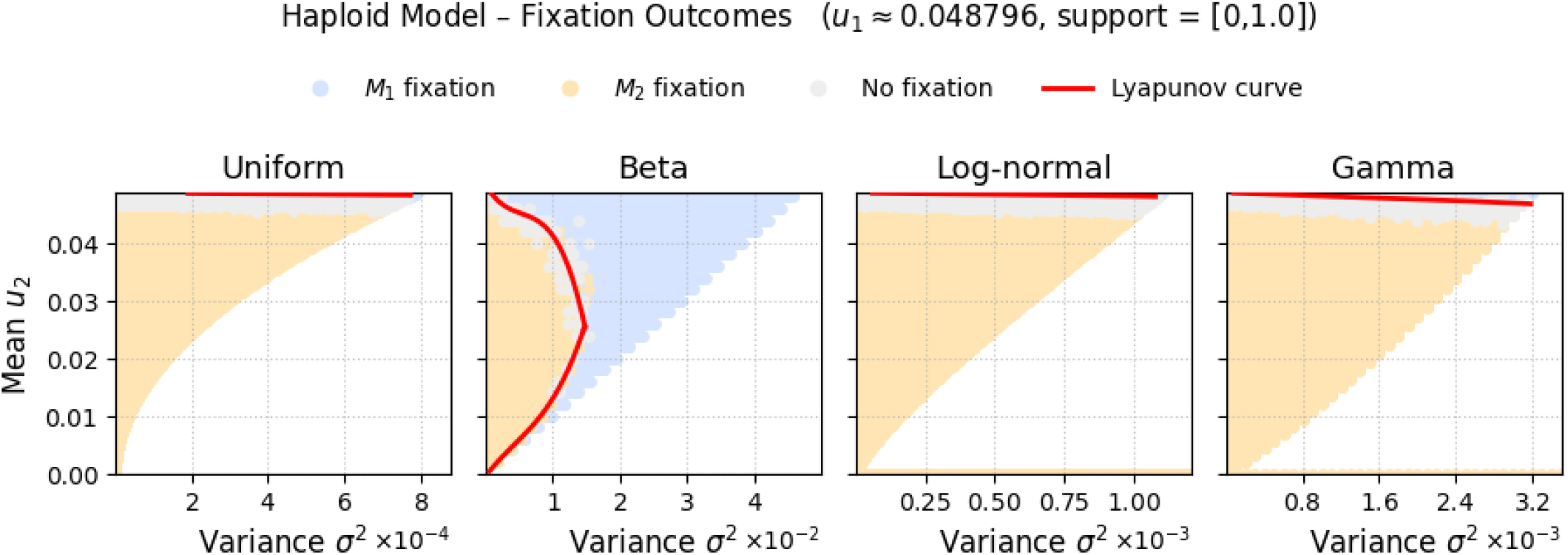
Fixation outcomes versus Lyapunov thresholds. Each panel: one mutation–rate distribution. Settings: 5,000 generations, *R* = 0, *s* = 0.2, *u*_1_ ≈ 0.048796. Colors: resident fixation (blue), modifier fixation (orange), no fixation within time range (gray). Red curve: Lyapunov prediction 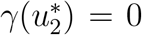. The predicted boundaries (red) closely tracks the observed transition for all families.

In contrast, distributions that shift mass into the right tail as variance increases (beta, truncated gamma) produce curved Lyapunov boundaries (red): larger *σ*^2^ raises the upper–tail contribution, lowers *E*[log(1 − *u*_2,*t*_)], and reduces 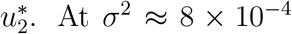, for example, the beta distribution gives 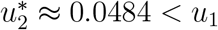, consistent with a modest right tail. The uniform is infeasible at this variance (its interval cannot match the target mean and contain *u*_1_). The truncated log–normal and truncated gamma yield 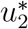 close to 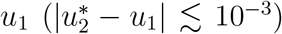, with the gamma slightly lower due to heavier midrange weight. In all cases, shifting probability from below *u*_1_ toward 1 decreases *E*[log(1 − *u*_2,*t*_)] and moves the boundary downward, as predicted by the Lyapunov condition.

## B Recombination effect on Invasion Condition Across Models

Table 4 summarizes invasion conditions for a rare modifier *M*_2_ under deterministic and stochastic transmission, distinguishing *R* = 0 from 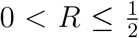. It indicates when recombination can alter the *direction* of selection on *M*_2_—that is, whether varying *R* can switch the outcome between invasion and non–invasion.

**Table 4:** Invasion conditions and recombination across deterministic and stochastic transmission. Deterministic transmission: the *Reduction Principle* holds for all *R*; recombination alters the magnitude but not the sign of initial growth. Stochastic transmission: at *R*=0, invasion depends on the geometric mean *E* [log(1 − *u*_2,*t*_)]; temporal variability can overturn the deterministic prediction even without recombination. For *R >* 0, *γ*(*R*) may change sign as *R* varies, producing reversals that do not occur when Var(*u*_2,*t*_)=0.

|  | Haploid |  | Diploid (additive) |  |
| --- | --- | --- | --- | --- |
|  | Deterministic | Stochastic | Deterministic | Stochastic |
| $R = 0$ | $\lambda_+ > 1 \iff u_2 < u_1$<br>(*Reduction Principle*;<br>(7)). | $\gamma(0) > 0 \iff$<br>$\mathbb{E}[\log(1 - u_{2,t})] >$<br>$\log(1 - u_1)$ ((12));<br>variability strictly<br>lowers $\mathbb{E}[\log(1 - u_{2,t})]$<br>at constant mean, so<br>invasion may fail even<br>if $u_2 < u_1$. | $\lambda_+ > 1 \iff u_2 < u_1$<br>(*Reduction Principle*;<br>(7)). | $\gamma(0) > 0 \iff$<br>$\mathbb{E}[\log(1 - u_{2,t})] >$<br>$\log(1 - u_1)$ ((12));<br>same conclusion. |
| $0 < R \leq \frac{1}{2}$ | Sign independent of<br>$R$; $R$ pulls $\lambda_+$ toward<br>1 (magnitude only). | $\gamma(R)$ exists (under<br>standard<br>moment/positivity<br>conditions) and can<br>change sign with $R$;<br>*reversals observed in*<br>*both directions.* | Sign independent of<br>$R$; $R$ pulls $\lambda_+$ toward<br>1 (magnitude only). | Same qualitative<br>behavior as haploids;<br>reversals possible,<br>with the $R$-response<br>typically smaller for<br>comparable<br>parameters. |

### Deterministic transmission (u_2,t_ ≡ u_2_)

For both ploidies, the invasion threshold is independent of *R*: *M*_2_ increases when rare if *u*_2_ *< u*_1_ (the *Reduction Principle*; Eq. (7)). For 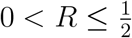, recombination moves the dominant eigenvalue toward 1 in magnitude but cannot change its sign. Thus, under deterministic transmission, recombination affects the *rate* but not the *direction* of invasion.

### Stochastic transmission (Var(u_2,t_) > 0)

With complete linkage (*R* = 0), both haploid and diploid models satisfy the geometric–mean criterion

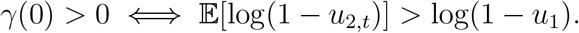

Because log(1 − *x*) is strictly concave on [0, 1), Jensen’s inequality implies

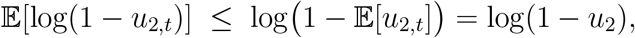

with strict inequality whenever Var(*u*_2,*t*_) *>* 0 and ℙ(*u*_2,*t*_ ∈ (0, 1)) *>* 0. Temporal variability therefore lowers the effective (geometric–mean) growth rate and can prevent invasion even when *u*_2_ *< u*_1_. Hence, the deterministic *Reduction Principle* may fail under stochastic transmission.

For *R >* 0, products of noncommuting random matrices preclude a closed–form expression for *γ*(*R*). Recombination has no direct fitness effect but affects associations between the modifier and the selected background that are created each generation by selection and mutation. Depending on how *R* modulates these associations across the realized distribution of *u*_2,*t*_, *γ*(*R*) can increase or decrease with *R*, allowing *reversals in invasion direction* under a fixed parameter set—an effect absent when Var(*u*_2,*t*_) = 0.

## C Empirical Analysis - *Arabidopsis thaliana*

We examine transmission–related determinants of mutation for genes annotated as essential or lethal in *A. thaliana*. These loci are expected to experience consistently strong purifying selection, reducing confounding from fluctuating selective regimes. Within this constrained functional class, residual variation in mutation rates should more plausibly reflect transmission mechanisms—chromatin configuration, local DNA accessibility, and DNA–repair efficacy—rather than changing selection pressures [22].

### Predictors of Mutation Rate

Following Monroe et al. [22], the response for gene *i* is the per–base mutation rate

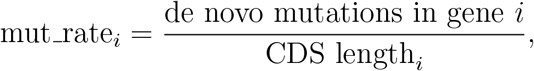

modeled as a function of sequence composition (GC content), cytosine methylation (CG/CHG/CHH), chromatin accessibility (ATAC–seq), and histone marks (H3K4me1/2/3, H3K27ac, H3K14ac, H3K27me1, H3K36ac/3, H3K56ac, H3K9ac/me1/me2, H3K23ac). Variables reported on a 0–100 scale were rescaled to [0, 1]; all predictors were standardized to *z*–scores. Thus, in linear models, coefficients represent the change in *expected* per–base mutation rate associated with a +1 SD change in a predictor, conditional on others at their means. Gene–level observations are treated as independent; heteroskedasticity and mean–variance coupling with gene length are addressed below.

We constructed a stable predictor set in three steps: *(i) Multicollinearity control:* we iteratively removed predictors with variance–inflation factor (VIF) *>* 10, a conventional threshold that guards against unstable coefficient estimation. *(ii) Multiple–testing control across models:* for each predictor, we computed Benjamini–Hochberg FDR *q*–values within each model and averaged these *q*–values across models to obtain a cross–model significance ranking. *(iii) Model diversity:* we fit three multivariable specifications to the same predictors:

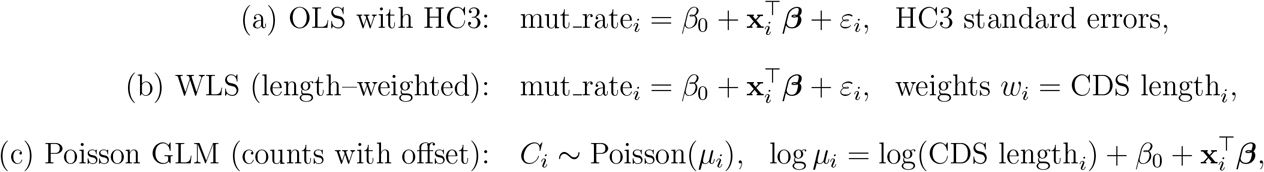

where *C*_*i*_ denotes the raw mutation count. In (c), exp(*β*_*j*_) is a rate ratio per +1 SD in predictor *j*, holding others fixed; in (a,b), *β*_*j*_ is a marginal slope for the per–base rate. HC3 ensures valid Wald tests under heteroskedasticity (given independence), while WLS stabilizes variance when the error variance scales with gene length.

To focus inference on effects that are stable across specifications and adjusted for multiplicity, we defined the *predictor set* by retaining the five predictors with the smallest mean BH–*q* across the three models (a pre–specified cap to promote parsimony and curb residual collinearity). The resulting set was *{*H3K4me1, H3K4me3, H3K36ac, GC content pct, CG pct*}*. We then refit the OLS model on this reduced set (after a second VIF screen to verify VIF ≤ 10 for all retained predictors) and summarized partial effects by varying one predictor at a time while fixing others at their standardized means (Fig. 5). This sequence—screen for collinearity, control FDR across complementary models, then refit on the selected subset—yields coefficient estimates and uncertainty that are interpretable, comparably scaled, and robust to reasonable changes in distributional assumptions.

**Figure 5:**
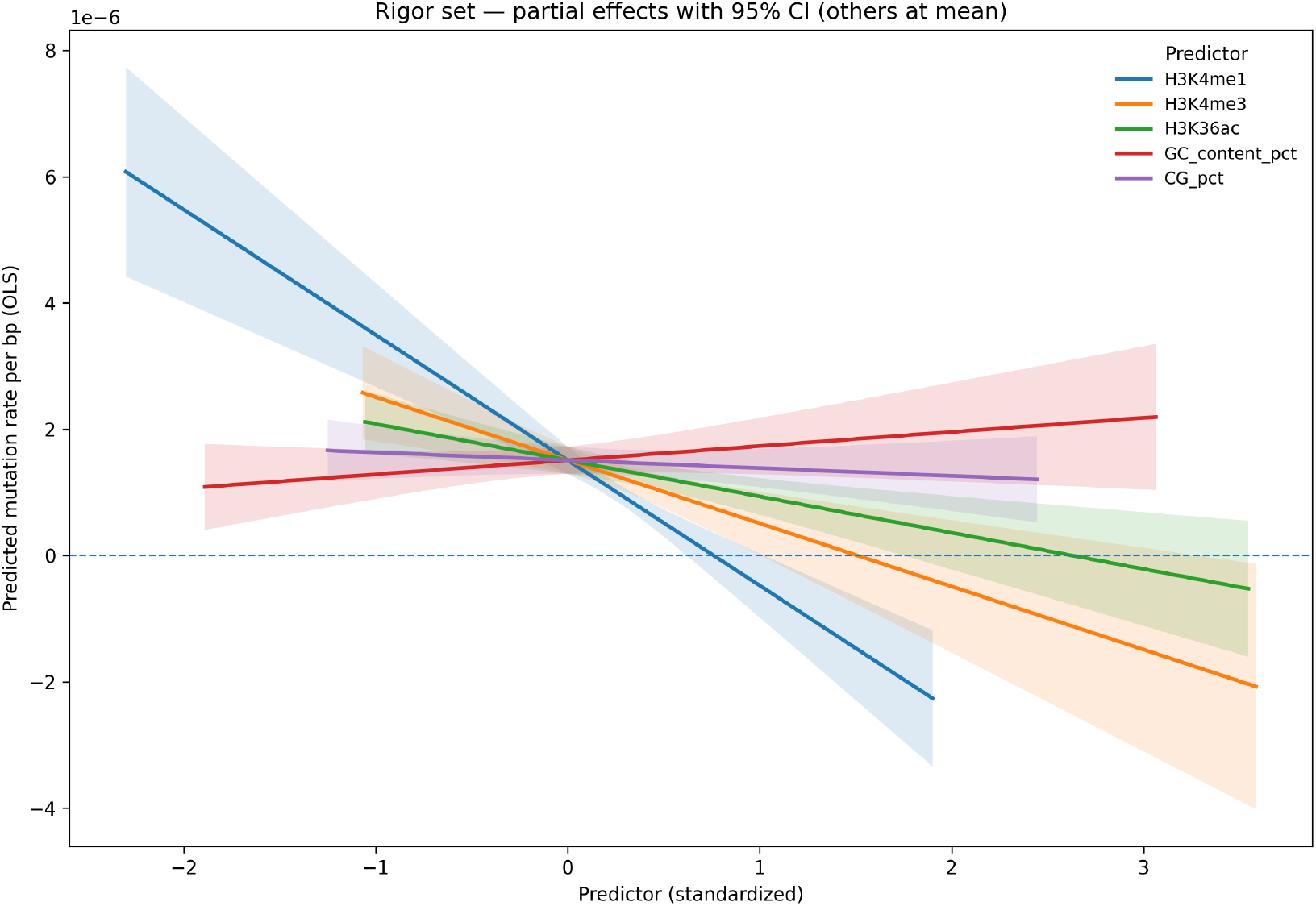
Rigor set—partial effects with 95% CI. OLS (HC3) partial–effect curves for the five rigor–set predictors. Each curve varies one standardized predictor across its 1st–99th percentile while holding the others at their means (0). Shaded bands are 95% confidence intervals; coefficients and FDR *q*–values for the same model are reported in the text.

In the joint model, three histone marks—H3K4me1, H3K4me3, and H3K36ac—showed strong negative associations with per–base mutation rate:

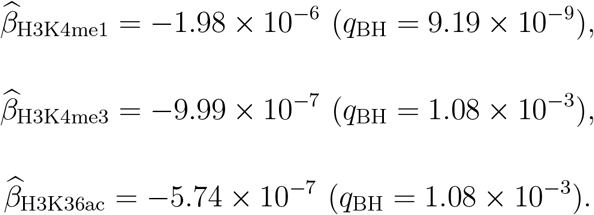

GC content showed a small, non–significant positive effect 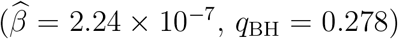, while CG methylation had a small, non–significant negative effect 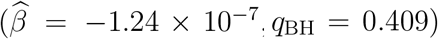. The signs and relative magnitudes of all coefficients were consistent across the WLS and Poisson–GLM models used for sensitivity analysis. The negative coefficients for H3K4me1, H3K4me3, and H3K36ac indicate that regions marked by these histone modifications are associated with lower mutation rates, consistent with enhanced D NA repair activity in transcriptionally active chromatin. In contrast, the weak methylation effect likely reflects reduction of its apparent influence on cecorrelated chromatin features are included [22].

Two caveats apply. First, restricting analysis to essential or lethal genes reduces variation in several predictors (e.g., GC content, chromatin accessibility), lowering statistical power and thereby reducing detectable marginal effects. Second, residual correlations among histone marks remain even after VIF pruning. Coefficients should therefore be interpreted as *partial associations conditional on the other included predictors*, not as independent causal effects.

### Predictors and Mutation Moments

To examine distributional properties in addition to the mean, we extended the regression to higher moments using the same standardized predictor set (*i*.*e*., each predictor *X*_*j*_ was centered and scaled as 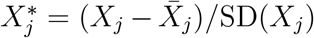 to have mean 0 and unit variance: H3K4me1, H3K4me3, H3K36ac, GC content, CG methylation). After fitting the mean model, we modeled dispersion and shape from the residuals: (i) a log–variance regression on squared residuals (reporting percent change in residual variance per +1 SD in a predictor), (ii) a skewness regression on cubed standardized residuals, and (iii) a kurtosis regression on the fourth standardized residual moment centered at 3. All specifications used heteroskedasticity–robust covariance and Benjamini–Hochberg adjustment. The variance models indicate strong stabilizing associations for active chromatin marks:

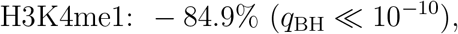

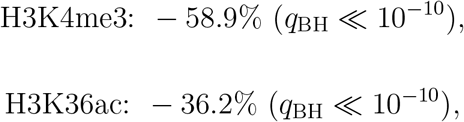

consistent with their negative mean effects (Fig. 5) and with reduced variance across genes (Fig. 6). In contrast, GC content shows a positive variance association (+38.7%, *q*_BH_ *<* 10^−11^), indicating greater mutation–rate variability in GC–rich regions. CG methylation exhibits a modest stabilizing association (−18.2%, *q*_BH_ ≈ 5 *×* 10^−6^).

**Figure 6:**
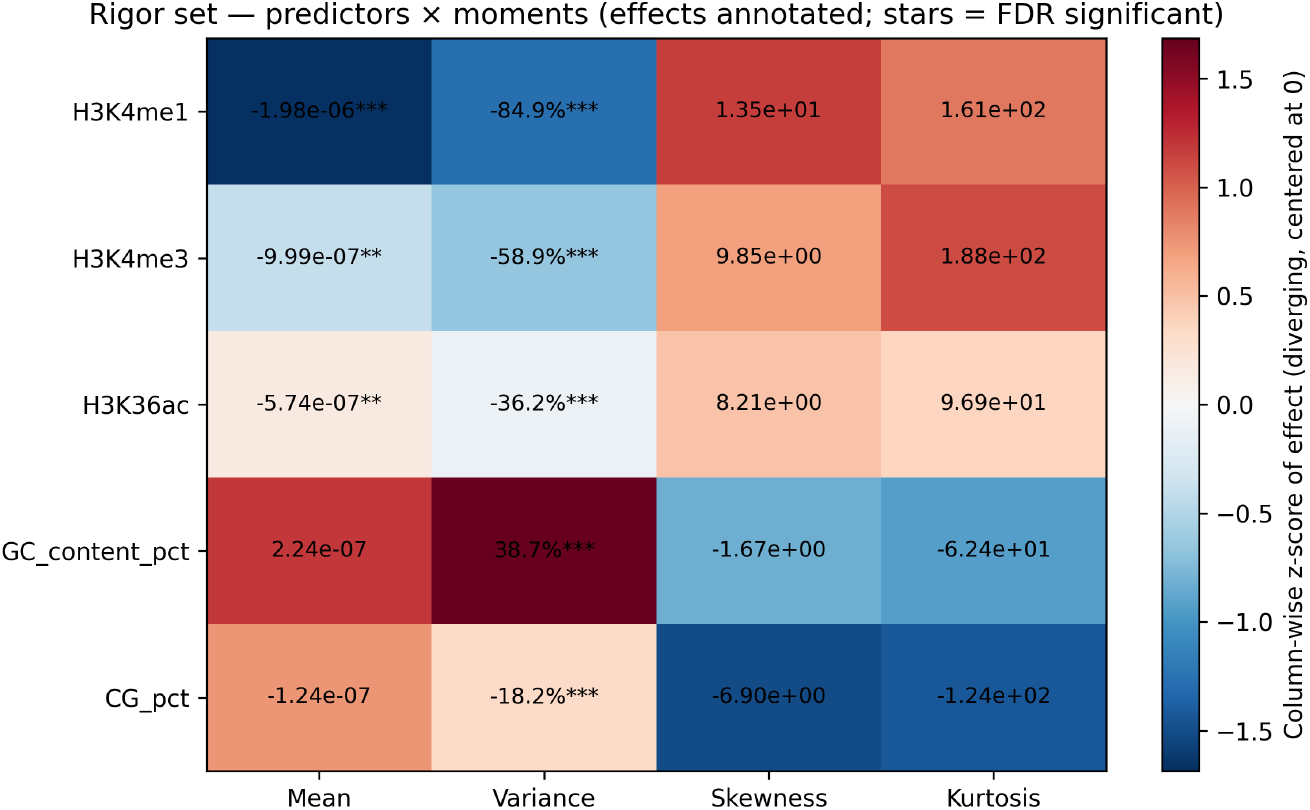
Predictors *×* moments of the per–bp mutation rate (rigor set, essential/lethal genes). Cells report regression coefficients for the mean (per–bp mutation rate), variance (percent change in residual variance per +1 SD), skewness, and kurtosis. Stars denote FDR significance. The color scale indicates column–wise standardized effect size (diverging, centered at 0).

For skewness and kurtosis, estimates are imprecise and not significant after FDR correction (all *q*_BH_ *>* 0.8). Point estimates suggest that GC content and CG methylation, while associated with higher mean and (for GC) higher variance, are simultaneously associated with lower skewness and kurtosis, whereas H3K4me1/3/36ac display the opposite qualitative pattern. Given the uncertainty of higher–moment regressions, these patterns should be interpreted as descriptive. They are nevertheless consistent with a potential trade–off in which predictors that increase overall dispersion (mean/variance) may correspondingly reduce tail weight (skewness/kurtosis), and vice versa.

Overall, H3K4me1, H3K4me3, and H3K36ac show a coherent stabilizing profile—lower mean and substantially lower variance—within these essential/lethal genes, while GC content increases variability despite a weak mean effect. This moment–based analysis complements the mean regression by indicating that epigenomic context is associated not only with the *level* of mutation but also with its *dispersion* and, tentatively, its tail shape across genes.

### Trends Across Tajima’s D Deciles

We examined whether a proxy for purifying selection covaries with (i) the epigenomic predictor set and (ii) the first four moments of the per–base mutation–rate distribution. For each essential/lethal gene *i*, let *D*_*i*_ denote Tajima’s *D*. With 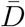 and sd(*D*) the sample mean and standard deviation over the analyzed genes, define a standardized selection score

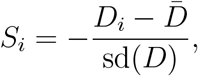

so larger *S*_*i*_ indicates more negative *D*_*i*_ (stronger putative purifying selection). All predictors were *z*–scored, and inference used heteroskedasticity–robust covariance with Benjamini–Hochberg FDR control.

To assess associations with the epigenomic predictors, we regressed each predictor *X*_*j*_ (H3K4me1, H3K4me3, H3K36ac, GC content pct, CG pct) on *S* while controlling for the other four predictors. The coefficient *γ*_*j*_ measures the change in *X*_*j*_ (SD units) per +1 SD increase in *S*. Only GC content pct increased significantly with *S* (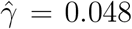, 95% CI [0.015, 0.081], *q* = 0.020). Partial slopes for the remaining predictors were small and non–significant: H3K4me1 (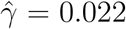, *q* = 0.142), CG pct (0.017, *q* = 0.432), H3K4me3 (0.010, *q* = 0.432), H3K36ac (−0.009, *q* = 0.494). Raw Spearman correlations between *S* and each predictor were near zero (e.g., GC content pct *ρ* ≈ 0.04), indicating that the GC trend arises only after conditioning on the other marks.

We next asked whether *S* predicts the mean and higher moments of the per–base mutation rate after adjusting for the predictor set. Specifically, we fit (i) a mean model for the mutation rate, (ii) a log–variance regression on squared residuals, (iii) a regression for skewness based on cubed standardized residuals, and (iv) a regression for kurtosis based on the fourth standardized residual moment centered at 3. Estimated effects of *S* per +1 SD were uniformly negative—mean 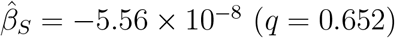, variance Δ = −5.2% (*q* = 0.185), skewness 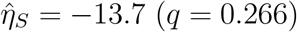, kurtosis 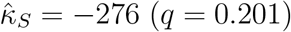—but none met the FDR threshold. The GC content result indicates that, within essential/lethal genes, stronger purifying selection is associated with modestly higher GC even after conditioning on chromatin marks. Possible mechanisms include GC–biased gene conversion or codon–usage constraints; the analysis is correlative and does not distinguish among them. For mutation moments, the negative point estimates are directionally consistent with lower means and reduced dispersion under stronger selection, but wide robust intervals indicate limited precision. Tajima’s *D* aggregates effects of demography and linked selection and is noisy for low–diversity, constraint–rich loci. Together with the essential/lethal restriction—which compresses both epigenomic and diversity variation—these factors limit power. Accordingly, all results should be interpreted as partial associations rather than causal effects (Fig. 7).

**Figure 7:**
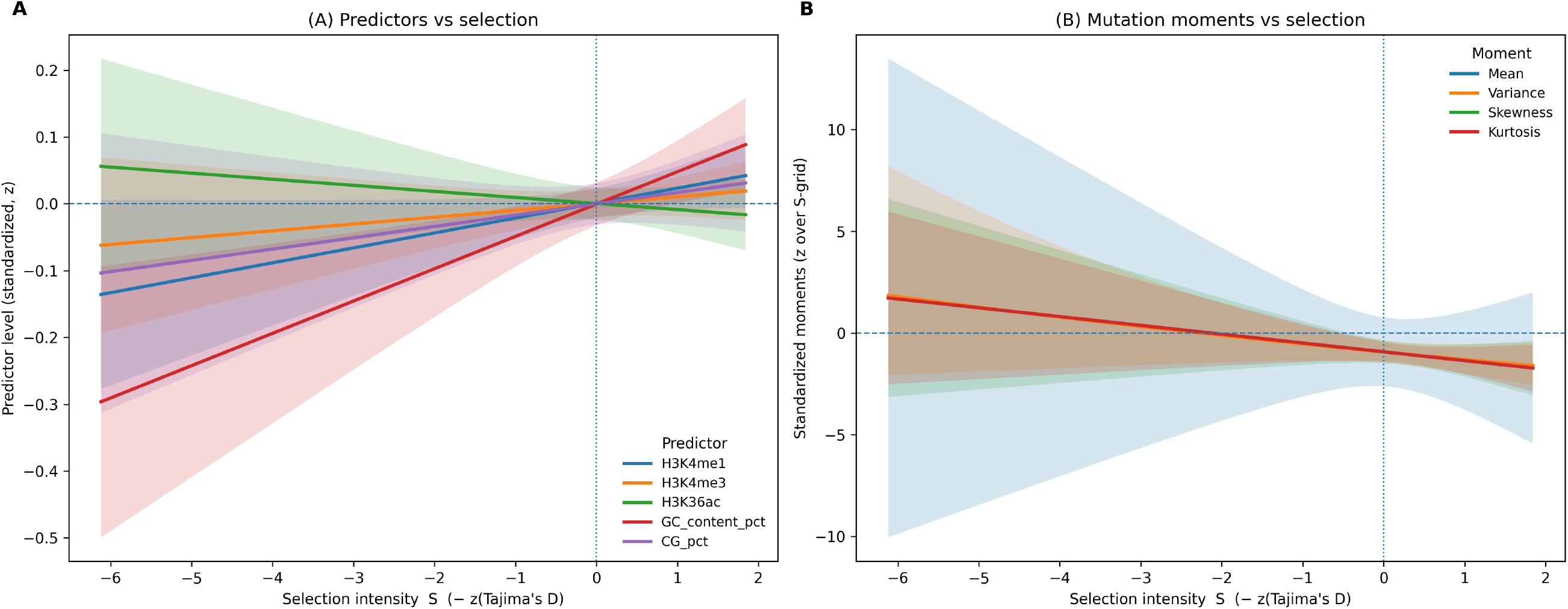
Selection intensity (*S* = −*z*(Tajima’s *D*)) versus predictors and mutation moments in essential/lethal genes. *Left (A):* Adjusted trends for each rigor predictor from *X*_*j*_ ∼ *S* + *X*_−*j*_ with robust 95% CIs; predictors are standardized (*z*; i.e., expressed in standard deviation units).. *Right (B):* Predicted mean, variance (reported as % change relative to *S*=0 and then standardized for display), skewness, and kurtosis from the moment models with robust 95% CIs; rigor predictors are held at their means. Larger *S* denotes stronger purifying selection.

### Interpretations

All analyses are restricted to essential/lethal genes, which reduces confounding from fluctuating selection but also compresses variation in predictors and outcomes, thereby lowering power. Results are therefore interpreted as partial associations, not causal effects.

In the joint linear model with standardized predictors, variance–inflation–factor (VIF) screening, and heteroskedasticity–robust inference, three histone marks linked to active chromatin show negative associations with the per–base mutation rate: H3K4me1 (−1.98 *×* 10^−6^ per +1 SD; *q* = 9.2*×*10^−9^), H3K4me3 (−9.99*×*10^−7^; *q* ≈ 1.1*×*10^−3^), and H3K36ac (−5.74*×*10^−7^; *q* ≈ 1.1 *×* 10^−3^). GC% and CG% exhibit small, non–significant mean effects (*q* ≈ 0.28 and *q* ≈ 0.41). Here, *q* denotes the Benjamini–Hochberg false–discovery–rate–adjusted *p*–value used for multiple–testing control. Thus, conditional on other marks, active chromatin is associated with lower average mutation rates, whereas base composition and CG methylation are not.

Extending the framework to residual dispersion reveals strong associations with variance. Per +1 SD in the predictor, residual variance decreases by 84.9% for H3K4me1, 58.9% for H3K4me3, and 36.2% for H3K36ac (all *q* ≪ 10^−10^), whereas GC% increases variance by 38.7% (*q* ≈ 1.3 *×* 10^−12^) and CG% reduces it by 18.2% (*q* ≈ 4.9 *×* 10^−6^). These results indicate that epigenomic context is associated not only with the mean level of mutation but also with the degree of *between–gene variability* (i.e., heterogeneity in mutation rates across genes). Regressions for skewness and kurtosis of variance–standardized residuals yield imprecise estimates (all *q* ≈ 0.82–0.90). Point estimates suggest opposite directions for active marks versus CG methylation, but the lack of FDR significance precludes inference about asymmetry or tail thickness.

Using *S* = −*z*(*D*) as a standardized proxy for stronger purifying selection (more negative Tajima’s *D*), we regressed each predictor on *S* while adjusting for the remainder. Only GC% increases with *S* (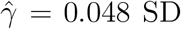 SD per +1 SD in *S*; 95% CI [0.015, 0.081]; *q* = 0.020), where 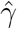 denotes the standardized partial regression slope. H3K4me1, H3K4me3, H3K36ac, and CG% show small, non–significant partial slopes (*q* ∈ [0.14, 0.49]). When the mean and higher moments of the mutation rate are modeled as functions of *S* (conditional on the chromatin predictors), estimated slopes are uniformly negative—mean −5.6 *×* 10^−8^ (*q* = 0.65), variance −5.2% (*q* = 0.19), skewness −13.7 (*q* = 0.27), kurtosis −276 (*q* = 0.20)—but none meet the FDR threshold.

Two implications follow. First, H3K4me1, H3K4me3, and H3K36ac are jointly associated with lower mutation means and substantially lower variance, whereas GC% increases variance despite a weak mean effect. Second, although estimates for each predictor versus *S* are imprecise, their consistently negative signs are compatible with stronger long–term purifying selection being associated with lower average mutation rates and reduced dispersion. Given limited power in the essential/lethal subset and the composite nature of Tajima’s *D*, these patterns should be viewed as suggestive and subject to confirmation in broader gene sets.

