## Supplementary material for "Evolution under Stochastic Transmission: Mutation-Rate Modifiers": Full Project Package: main.tex

\documentclass[12pt]{article}

% --------------------------------------------------
% Packages
% --------------------------------------------------
\usepackage[utf8]{inputenc}

\usepackage{amsmath,amssymb,amsthm,mathtools}
\usepackage{bm}
\usepackage{subcaption}
\usepackage{graphicx}
\usepackage{array,booktabs,tabularx,multirow}
\usepackage{pdflscape}
\usepackage{enumitem}
\usepackage{appendix}
\usepackage{tikz}
\usetikzlibrary{matrix,positioning}
\usepackage{natbib}
\usepackage{setspace}
\usepackage{tocloft}
\usepackage[hidelinks]{hyperref} 
\allowdisplaybreaks

\DeclareUnicodeCharacter{2234}{\therefore}
\DeclareUnicodeCharacter{202F}{~}

% Column types
\newcolumntype{P}[1]{>{\raggedright\arraybackslash}p{#1}}
\newcolumntype{Y}{>{\raggedright\arraybackslash}X}

% Theorem-like environments
\newtheorem{lm}{Lemma}
\newtheorem{remark}{Remark}
\newtheorem{proposition}{Proposition}
\newtheorem{corollary}{Corollary}

% Margins: 1in all around (typical TPB/Elsevier review style)
\usepackage[margin=1in]{geometry}

% Title/author handling
\usepackage{authblk}

% TOC / lists tweaks 
\renewcommand{\cftdotsep}{\cftnodots}
\cftpagenumbersoff{figure}
\cftpagenumbersoff{table}

% Simple keywords macro for article class
\newcommand{\keywords}[1]{%
  \vspace{0.5em}%
  \noindent\textbf{Keywords: }#1
}

% --------------------------------------------------
% Title / authors / affiliations
% --------------------------------------------------
\title{Evolution under Stochastic Transmission:\\ Mutation-Rate Modifiers}

\author[1]{Elisa Heinrich-Mora}

\affil[1]{Department of Biology, Stanford University, Stanford, CA, USA}

\date{}

% --------------------------------------------------
\begin{document}
% --------------------------------------------------

% -------- TITLE PAGE --------
\maketitle
\thispagestyle{plain} 
\clearpage           

% -------- ABSTRACT PAGE --------
\begin{abstract}
In evolutionary models of large populations, it is common to analyze the effects of cyclic or random variation in the parameters that describe selection. It is less common, however, to study how stochasticity in the genetic transmission process itself affects evolutionary outcomes. Suppose that a gene locus has alleles $A$ and $a$ under constant selection. This locus is linked to a modifier locus with alleles $M_1$ and $M_2$, which control the mutation rate from $A$ to $a$. The \textit{Reduction Principle} states that, near a mutation--selection balance where $M_1$ is fixed with mutation rate $u_1$, a rare allele $M_2$ can invade if its associated rate $u_2$ is lower than $u_1$. This result, valid for both haploids and diploids, assumes constant mutation rates through time. We extend this framework by allowing the mutation rate associated with $M_2$ to fluctuate randomly across generations, denoted as $u_{2,t}$. In this stochastic setting, the condition for invasion by a new modifier allele depends not only on the resident mutation rate $u_1$ and the mean mutation rate $u_2$ associated with the invading allele, but also on the temporal distribution of $u_{2,t}$, the strength of selection at the $A/a$ locus, and the recombination rate between $M_1/M_2$ and $A/a$. The analysis shows how stochasticity and recombination in transmission do not simply modify the magnitude of evolutionary change predicted under deterministic assumptions. Instead, through their interaction with selection and linkage, they can generate conditions under which the direction of modifier evolution is qualitatively reversed relative to the deterministic \textit{Reduction Principle}.
\end{abstract}

\keywords{mutation-rate modifiers; stochastic transmission; mutation--selection balance; reduction principle; recombination; evolutionary genetics}

\clearpage 

\newpage
\doublespacing
\section*{Introduction}
\noindent An important goal of evolutionary biology is to identify the forces that shape the structure and dynamics of genetic systems. Standard population–genetic models often treat aspects of transmission (e.g. mutation, recombination rates) and genetic structure (e.g. dominance, epistasis) as fixed. Empirical work, however, has shown that these parameters can themselves be subject to evolutionary modification. \textit{Modifier theory} makes this point explicit by analyzing loci whose alleles affect the transmission and/or genetic structure at a primary locus under selection—for example, modifiers of mutation, recombination, or dominance. Invasion analysis of a rare modifier allele identifies which changes are initially favored, and provide insight into how the such parameters might evolve. It is useful to distinguish modifiers with \emph{direct} fitness effects—those that alter viability or fertility through changes in parameters such as dominance or epistasis—from selectively \emph{neutral} modifiers, which have no intrinsic effect on fitness \citep{altenberg1984}, such as those influencing transmission parameters (e.g., mutation or recombination rates). The former change in frequency through both their direct fitness effects and associations with selected loci, whereas the latter evolve solely through such associations. 

\noindent A key result from this framework is the \emph{Reduction Principle}, which states that modifier alleles that reduce mutation rates at loci under mutation–selection balance, or that reduce recombination rates between loci with epistatic interactions that generate linkage disequilibrium, increase in frequency due to their indirect effects on the selected loci \citep{feldman1972selection, karlin1974towards, teague1976result, altenberg1984, feldman1986evolutionary, altenberg2017unified, feldman1976krakauer, balkau1973selection, liberman1986modifiers, feldman1996population}. The principle holds under assumptions of infinite population size, random mating, constant viability selection, and deterministic dynamics—prompting studies to examine departures from these conditions.

\noindent A substantial body of theoretical work has investigated how temporal fluctuations in selection parameters can influence long-term evolutionary outcomes. These studies treat fitness values as random variables that fluctuate across generations.  Gillespie \citep{gillespie1972, gillespie1973}, Cook and Hartl \citep{cookHartl1974}, and Karlin and Liberman \citep{karlinLiberman1975} analyzed such scenarios, demonstrating that fluctuating selection can—under specified conditions—maintain polymorphism and favor the allele with the highest geometric mean fitness across time, which is shaped by both the mean and higher moments of the fitness distribution. Related results in population growth theory show that, in randomly varying environments, multiplicative growth is governed by the geometric mean of growth factors \citep{lewontin1969population}. In these analyses, stochasticity enters through selection; transmission parameters remain fixed.

\noindent \textit{Modifier theory} has also been extended to scenarios with variable selection regimes. In these models, the fitness landscape changes over time while the fundamental assumptions of modifier theory are retained. Indirect selection on modifier alleles can arise from spatial or temporal fluctuations in selection coefficients \citep{charlesworth1976recombination, carja2014evolution}, and environmental variability can favor nonzero mutation or switching rates by altering the geometric mean fitness of genotypes across cycles \citep{Liberman2011}. Recent simulation studies of multigenic mutation–rate modifiers in sexual populations further demonstrate that the form of selection acting on a quantitative trait—stabilizing versus directional—can bias the evolution of mutation rates through associations between modifier alleles and individuals in the phenotypic tails \citep{rozhok2021silico}. Together, these findings suggest that the evolution of transmission parameters such as mutation rate depends not only on the expected magnitude of environmental change but also on its distribution and directional properties.

\noindent There is evidence that transmission parameters can vary through endogenous processes even under constant external selective conditions. In \textit{Arabidopsis thaliana}, mutation–accumulation experiments show genome–wide heterogeneity in mutation rates that correlates with epigenetic and physical genomic features rather than with direct natural selection \citep{monroe2022}. In \textit{Escherichia coli}, long–term evolution experiments reveal “hypermutator” alleles—typically defects in DNA repair—whose effects on mutation rate are not strictly monotone with environmental conditions \citep{wei2022}. In humans, variation in recombination rate is influenced by genetic variants—most notably in \textit{PRDM9} and \textit{RNF212}—as well as by structural variation, with little evidence for direct effects on fitness \citep{FledelAlon2011}. These observations suggest that the mechanisms of transmission are themselves evolvable traits, capable of generating variation even in the absence of external change.

\noindent If endogenous processes generate fluctuations in epigenetic or physical genomic features that influence transmission parameters, a natural question arises: are genetic modifiers subject to selection not only through their mean effects, but also through their \emph{variability} across generations? To address this, we analyzed \textit{A.~thaliana} essential and lethal genes (Appendix~\ref{appendix:empirical}) using gene–level mutation data from Monroe et al. \cite{monroe2022}. In multivariable models, chromatin marks associated with active repair (H3K4me1, H3K4me3, H3K36ac) were linked to large reductions in mean mutation rates and, more strongly, reduction in their variance. Although formal selection tests were underpowered, regressions of a standardized Tajima’s $D$ score on moments of the mutation–rate distribution showed similar tendencies: stronger purifying selection coincided with lower means and lighter tails. These patterns are consistent with selection in functionally constrained genomic regions, acting to reduce not only the average mutation rate but also its variability. Modifier alleles that permit occasional high mutation rates may be disfavored relative to those that suppress variance while maintaining a comparable mean mutation rate.

\noindent We therefore consider models of stochasticity in the \emph{transmission} process. Rather than fixing transmission parameters while allowing selection to vary, we hold selection constant and allow transmission parameters to fluctuate across generations. The focus is the invasion of a rare modifier allele that reduces the average transmission rate while introducing inter-generational variance in that same parameter. The central question is whether indirect selection on the modifier still conforms to the \textit{Reduction Principle} or whether variance in the transmission parameter alters this outcome.

\noindent Let $T$ denote a transmission parameter with mean $\mu=\mathbb{E}[T]$ and variance $\sigma^2=\mathrm{Var}(T)$. In each generation, the value of $T$ is drawn independently from this distribution. Classical modifier theory treats $T$ as constant. When $T$ fluctuates, the variance $\sigma^2$, even with constant $\mu$, may affect the initial increase of a new modifier allele. This effect is captured by Lyapunov exponents of the linearized recursions for the modifier haplotype frequencies, which depend on the full distribution of transmission parameters. Hence invasion depends on both the average effect on transmission and higher–order statistics of the transmission process. Variability in transmission is therefore not simply an extraneous source of noise, but a factor that can alter the magnitude and direction of evolutionary outcomes. Extending modifier theory to include fluctuating transmission thus provides a general framework for analyzing how endogenous variability in transmission rates influences the evolution of genetic systems. 

\section*{Mutation Modification}\label{sec: mutation}
\noindent Consider a large, randomly mating population with discrete generations. A single locus under constant selection carries alleles $A$ and $a$; a linked, selectively neutral modifier locus carries alleles $M_1$ and $M_2$ that determine the forward mutation rate at the selected locus. Under $M_1$, the mutation rate is constant at $u_1$ across all generations, and the selected locus is at its deterministic mutation–selection balance. At this equilibrium, a rare modifier allele that reduces the constant rate ($u_2<u_1$) is favored—the \textit{Reduction Principle} \citep{karlin1974random, altenberg1984, feldman1986evolutionary, altenberg2017unified,liberman1986modifiers}. 

\noindent Here we ask whether that principle holds when the modifier allele influences not only the mean mutation rate but also its variability. Specifically, $M_2$ generates a mutation rate sequence $\{u_{2,t}\}$ with expectation $\mathbb{E}[u_{2,t}]=u_2$ and variance $\mathrm{Var}(u_{2,t})=\sigma^2$ across generations, while $u_1$ remains constant and selection at $A/a$ is time-invariant. Our objective is to derive the conditions under which $M_2$, introduced at low frequency near a mutation-selection balance with $M_1$, can invade in haploid or diploid populations, and to elucidate how the temporal distribution of the mutation rate, together with selection strength and recombination between the modifier and selected loci, affect these invasion dynamics.

\section{Model Set-up} \label{sec: model_setup_unified}
\noindent A large population is considered to have two biallelic loci: a selected locus ($A/a$) under constant selection and a linked, selectively neutral modifier ($M_1/M_2$) that controls the \emph{forward} mutation rate $A\to a$. Back mutation $a\to A$ is absent. The recombination fraction between loci is $R\in[0,\tfrac12]$, constant through time. The life cycle within each generation is
\[
\mathbf x \xrightarrow{\ \text{selection}\ }\mathbf x'
\xrightarrow{\ \text{recombination}\ }\mathbf x''
\xrightarrow{\ \text{mutation}\ }\mathbf x^{(t+1)}.
\]
\noindent Let $\mathbf x=(x_1,x_2,x_3,x_4)$ denote haplotype frequencies $(AM_1,aM_1,AM_2,aM_2)$ with $\sum_i x_i=1$. After selection and recombination the frequencies are $\mathbf x'$ and $\mathbf x''$, respectively; $\mathbf x^{(t+1)}$ denotes the state after mutation (start of generation $t{+}1$).

\paragraph{Selection.} Selection acts only at $A/a$. For haploids, viabilities are $W_A=1$ and $W_a=1-s$ with $s\in(0,1]$, yielding mean fitness
\[
\bar W_{\mathrm{hap}}=(x_1+x_3)+(1-s)(x_2+x_4),
\]
and 
\[
x_1'=\frac{x_1}{\bar W_{\mathrm{hap}}},\quad
x_2'=\frac{(1-s)x_2}{\bar W_{\mathrm{hap}}},\quad
x_3'=\frac{x_3}{\bar W_{\mathrm{hap}}},\quad
x_4'=\frac{(1-s)x_4}{\bar W_{\mathrm{hap}}}.
\]
\noindent For diploids, viabilities at the selected locus are additive:
\[
W_{11}=1\ (AA),\quad W_{12}=1-s\ (Aa),\quad W_{22}=1-2s\ (aa),\quad s\in(0,1).
\]
\noindent Random mating (Hardy–Weinberg before selection) and viability selection entail that the mean fitness in diploids is
\[
\bar W_{\mathrm{dip}}=(x_1+x_3)^2 W_{11}+2(x_1+x_3)(x_2+x_4)W_{12}+(x_2+x_4)^2 W_{22}.
\]
\noindent The resulting gamete frequencies, $x_i'=x_i'(\mathbf x; s)$ after selection are the standard two-locus frequencies with viabilities $W_{11},W_{12},W_{22}$.

\paragraph{Recombination.} Recombination acts on $\mathbf x'$ in haploids and diploids at the gamete level. With linkage disequilibrium after selection, \(D' \;=\; x_1' x_4' - x_2' x_3',\) after recombination we have
\[
x_1''=x_1'-R D',\quad
x_2''=x_2'+R D',\quad
x_3''=x_3'+R D',\quad
x_4''=x_4'-R D'.
\]
\paragraph{Mutation.} Forward mutation $A\to a$ is modifier dependent and operates on $A$-bearing gametes after recombination. In the haploid case, $M_1$ and $M_2$ produce constant per-generation rates $u_1,u_2\in[0,1)$:
\begin{equation}\label{eq:hap_mutation}
x_1^{(t+1)}=(1-u_1)\,x_1'',\quad
x_2^{(t+1)}=x_2''+u_1\,x_1'',\quad
x_3^{(t+1)}=(1-u_2)\,x_3'',\quad
x_4^{(t+1)}=x_4''+u_2\,x_3''.
\end{equation}
\noindent In diploids, the mutation rate depends on the modifier \emph{genotype}, and gametes produced from $M_1M_1$, $M_1M_2$, and $M_2M_2$ zygotes have rates $u_1,u_2,u_3$; the evolutionary recursions are obtained by applying these rates to the appropriate gamete contributions after selection and recombination.

\paragraph{Full recursion.} Substituting the parameters at each step yields:
\begin{itemize}
    \item \emph{Haploids:}
    \begin{equation}
    \label{eq:hap_recursion}
    \small
    \begin{aligned}
    x_1^{(t+1)} 
    &= \frac{x_1}{\bar{W}_{\mathrm{hap}}} (1 - u_1) - 
    \frac{R(1 - s) (x_1 x_4 - x_2 x_3)}{\bar{W}_{\mathrm{hap}}^2} (1 - u_1),\\
    x_2^{(t+1)} 
    &= \frac{(1 - s)x_2}{\bar{W}_{\mathrm{hap}}} + \frac{R(1 - s)(x_1 x_4 - x_2 x_3)}{\bar{W}_{\mathrm{hap}}^2} + 
    \frac{x_1}{\bar{W}_{\mathrm{hap}}} u_1 - \frac{R(1 - s)(x_1 x_4 - x_2 x_3)}{\bar{W}_{\mathrm{hap}}^2} u_1,\\
    x_3^{(t+1)} 
    &= \frac{x_3}{\bar{W}_{\mathrm{hap}}} (1 - u_2) + \frac{R(1 - s)(x_1 x_4 - x_2 x_3)}{\bar{W}_{\mathrm{hap}}^2} (1 - u_2),\\
    x_4^{(t+1)} 
    &= \frac{(1 - s)x_4}{\bar{W}_{\mathrm{hap}}} - \frac{R(1 - s)(x_1 x_4 - x_2 x_3)}{\bar{W}_{\mathrm{hap}}^2} + 
    \frac{x_3}{\bar{W}_{\mathrm{hap}}} u_2 + \frac{R(1 - s)(x_1 x_4 - x_2 x_3)}{\bar{W}_{\mathrm{hap}}^2} u_2,
    \end{aligned}
    \end{equation}
    with  $\bar W_{\mathrm{hap}}=(x_1+x_3)+(1-s)(x_2+x_4)$.
    
    \item \emph{Diploids} \cite{TWOMEY1990320}:
    \begin{equation}
    \label{eq:dip_recursion}
    \begin{aligned}
    \bar{W}_{\mathrm{dip}}\, x_1^{(t+1)} &= (1 - u_1)(x_1^2 W_{11} + x_1 x_2 W_{12}) 
    + (1 - u_2)\Big[x_1 x_3 W_{11} + x_1 x_4 W_{12} - R W_{12}(x_1 x_4 - x_3 x_2)\Big], \\
    \bar{W}_{\mathrm{dip}}\, x_2^{(t+1)} &= x_2^2 W_{22} + x_1 x_2 W_{12} + x_3 x_2 W_{12} + x_2 x_4 W_{22} 
    + R W_{12}(x_1 x_4 - x_3 x_2) \\
    &\quad + u_1(x_1^2 W_{11} + x_1 x_2 W_{12}) 
    + u_2\Big[x_1 x_3 W_{11} + x_1 x_4 W_{12} - R W_{12}(x_1 x_4 - x_3 x_2)\Big], \\
    \bar{W}_{\mathrm{dip}}\, x_3^{(t+1)} &= (1 - u_3)(x_3^2 W_{11} + x_3 x_4 W_{12}) 
    + (1 - u_2)\Big[x_1 x_3 W_{11} + x_3 x_2 W_{12} 
    + R W_{12}(x_1 x_4 - x_3 x_2)\Big], \\
    \bar{W}_{\mathrm{dip}}\, x_4^{(t+1)} &= x_4^2 W_{22} + x_1 x_4 W_{12} + x_3 x_4 W_{12} + x_2 x_4 W_{22} 
    - R W_{12}(x_1 x_4 - x_3 x_2) \\
    &\quad + u_3(x_3^2 W_{11} + x_3 x_4 W_{12}) 
    + u_2\Big[x_1 x_3 W_{11} + x_3 x_2 W_{12} + R W_{12}(x_1 x_4 - x_3 x_2)\Big],
    \end{aligned}
    \end{equation}
    with $\bar W_{\mathrm{dip}}=(x_1+x_3)^2 W_{11}+2(x_1+x_3)(x_2+x_4)W_{12}+(x_2+x_4)^2 W_{22}$.
    \end{itemize}

\noindent Recursion systems \eqref{eq:hap_recursion} and \eqref{eq:dip_recursion} are the basis for the invasion analysis in which $M_2$ is initially rare and the resident $M_1$ is at its mutation–selection equilibrium.

\section{Invasion Analysis of Modifier Allele $M_2$}\label{sec:invasion_unified}
\noindent We ask whether a rare modifier allele $M_2$ increases in frequency when introduced near a resident population fixed for $M_1$. Let $\mathbf x_t=(x_1,x_2,x_3,x_4)^\top$ denote the haplotype frequencies of $AM_1$, $aM_1$, $AM_2$, and $aM_2$ at generation $t$, and let $\hat{\mathbf x}=(\hat x_1,\hat x_2,0,0)^\top$ denote the resident equilibrium (mutation–selection balance) at the selected locus $A/a$. Using the life–cycle order specified in Section~\ref{sec: model_setup_unified}, the one–locus subsystem at equilibrium satisfies $x_{i,t+1}=x_{i,t}$, with
\begin{subequations}\label{eq:mut_sel_balance}
\begin{align}
\text{haploids:}\quad
(\hat x_1,\hat x_2)&=\Big(\tfrac{s-u_1}{s},\,\tfrac{u_1}{s}\Big),\quad 
\hat{\bar W}_{\mathrm{hap}}=1-u_1
\quad\text{for }0<u_1<s, \label{eq:4a}\\[4pt]
\text{diploids:}\quad
(\hat x_1,\hat x_2)&=\Big(\tfrac{s-u_1+su_1}{s(1+u_1)},\,\tfrac{u_1}{s(1+u_1)}\Big),\quad 
\hat{\bar W}_{\mathrm{dip}}=\tfrac{1-u_1}{1+u_1}
\quad\text{for }0<u_1<\tfrac{s}{1-s}. \label{eq:4b}
\end{align}
\end{subequations}
\noindent Linearizing the full two–locus recursions system at $\hat{\mathbf x}$ yields the Jacobian
\[
\mathbf J=\begin{pmatrix}\mathbf P & \mathbf C\\[3pt] \mathbf 0 & \mathbf F\end{pmatrix},
\]
where $\mathbf P$ describes the resident $(AM_1,aM_1)$ subsystem, $\mathbf C$ captures first–order coupling from residents to rare haplotypes, and $\mathbf F$ governs the rare subsystem associated with $M_2$. Because $\mathbf J$ is block–triangular, the eigenvalues of $\mathbf F$ determine the local stability of the $M_1$ equilibrium with respect to invasion by $M_2$. Consequently, the frequencies of the rare haplotypes $\bm v_t=(x_{3,t},x_{4,t})^\top$ satisfy
\[
\bm v_{t+1}=\mathbf F\,\bm v_t,
\]
and invasion occurs when the dominant eigenvalue (Perron root) of $\mathbf F$, $\rho(\mathbf F)$, exceeds unity.

\noindent Substituting $(\hat{x}_1,\hat{x}_2)$ from \eqref{eq:mut_sel_balance} into the $x_{3,t+1}$ and $x_{4,t+1}$ components of the recursion systems \eqref{eq:hap_recursion} and \eqref{eq:dip_recursion} and discarding higher–order terms gives the explicit forms:
\begin{itemize}
    \item \textit{Haploids.}
\begin{equation}\label{eq:F_hap}
    \mathbf F_{\mathrm{hap}}=\frac{1}{1-u_1}
    \begin{pmatrix}
    (1-u_2)\Big[1-\dfrac{u_1 R(1-s)}{s(1-u_1)}\Big] &
    (1-u_2)\dfrac{(s-u_1) R(1-s)}{s(1-u_1)}\\[6pt]
    u_2+(1-u_2)\dfrac{u_1 R(1-s)}{s(1-u_1)} &
    (1-s)-(1-u_2)\dfrac{(s-u_1) R(1-s)}{s(1-u_1)}
    \end{pmatrix}.
\end{equation}

    \item \textit{Diploids.}
    \begingroup
    \setlength{\arraycolsep}{3pt}
    \begin{equation}\label{eq:F_dip}
      \mathbf F_{\mathrm{dip}}
      = \frac{1}{(1-u_1)s}
      \begin{pmatrix}
        (1-u_2)\bigl[s - u_1R(1-s)\bigr]
          &
        (1-u_2)(1-s)R\,\bigl[s - u_1(1-s)\bigr]
        \\[4pt]
        u_1(1-s)R + u_2\bigl[s - u_1R(1-s)\bigr]
          &
        \begin{aligned}
          (1-s)\bigl[(1-R)+u_2R\bigr]\bigl[s - u_1(1-s)\bigr]\\
          {}+ (1-2s)u_1
        \end{aligned}
      \end{pmatrix}.
    \end{equation}
    \endgroup
    
\end{itemize}
\noindent Thus the invasion dynamics depend on selection $s$, recombination $R$, and the resident mutation rate $u_1$ through the entries of $\mathbf F$. Because the modifier has no direct fitness effect, its initial change is driven by the dynamics of the $M_2$–bearing haplotypes $(AM_2,aM_2)$ near the resident haplotypes $AM_1, aM_1$. Selection at the $A/a$ locus generates the associations that facilitate this growth, whereas recombination modulates these associations each generation, as reflected in the $R$–dependent terms of $\mathbf F$. The condition $\rho(\mathbf F)>1$ defines the criterion for invasion.

\subsection{\textbf{Deterministic Case ($u_2$ constant).}} \label{sec:invasion_deterministic_unified}
\noindent Near the resident mutation–selection equilibrium $\hat{\mathbf x}=(\hat x_1,\hat x_2,0,0)^\top$ with modifier $M_1$, and forward mutation rate $u_1$, introduce a rare modifier $M_2$ with constant mutation rate $u_2$. Linearizing at $\hat{\mathbf x}$ yields $\bm v_{t+1}=\mathbf F\,\bm v_t$, where $\bm v_t=(x_{3,t},x_{4,t})^\top$ collects the rare haplotype frequencies and $\mathbf F$ is a nonnegative $2\times2$ matrix (with positive entries in the parameter ranges considered). By Perron–Frobenius \cite{frobenius1908_1909}, $\mathbf F$ has a simple dominant eigenvalue $\lambda_+=\rho(\mathbf F)>0$, and $M_2$ increases when $\lambda_+>1$.

\noindent The characteristic polynomial is $p(\lambda)=\lambda^2-\tau\lambda+\delta$, where $\tau=\mathrm{tr}(\mathbf F)$ and $\delta=\det(\mathbf F)$. Since $p(\lambda)=(\lambda-\lambda_-)(\lambda-\lambda_+)$ with $0<\lambda_-\le\lambda_+$, we have $p(1)=(1-\lambda_-)(1-\lambda_+)$. Thus, if $\lambda_-<1$ (which holds at the resident equilibrium under the admissible parameter ranges), then $p(1)<0$ if $\lambda_+>1$. Because $p(1)=1-\tau+\delta$, it suffices to evaluate $p(1)$ at the resident equilibrium. Direct calculation gives
\[
\text{haploids: } p(1)=\frac{(s-u_1)(u_2-u_1)}{(1-u_1)^2}, \qquad
\text{diploids: } p(1)=\frac{\big[s(1+u_1)-u_1\big](u_2-u_1)}{(1-u_1)^2}.
\]
\noindent For haploids, equilibrium requires $0<u_1<s$, so $s-u_1>0$; for the additive diploid model, $s>\tfrac{u_1}{1+u_1}$, so $s(1+u_1)-u_1>0$. Under these conditions, $p(1)$ has the sign of $(u_2-u_1)$, and therefore
\begin{equation} \label{eq:deterministic_reduction_principle}
    u_2<u_1 \;\Longrightarrow\; p(1)<0 \;\Longrightarrow\; \lambda_+>1.
\end{equation}
\noindent This is the \emph{Reduction Principle}: a modifier allele that lowers the mutation rate ($u_2<u_1$) will invade.

\noindent \paragraph{Effect of recombination on $M_2$ invasion.} Although recombination $R$ does not affect the \emph{criterion} for invasion—since $p(1)=1-\tau+\delta$ is independent of $R$—it affects the \emph{magnitude} of growth through the Perron root $\lambda_+=\tfrac{\tau+\sqrt{\tau^2-4\delta}}{2}$. Writing the characteristic equation $p(\lambda)=\lambda^2-\tau\lambda+\delta=0$ and differentiating implicitly with respect to $R$ gives
\[
(2\lambda_+-\tau)\,\frac{d\lambda_+}{dR}-\frac{d \tau(R)}{dR}\lambda_+(R)+\frac{d\delta(R)}{dR}=0
\quad\Longrightarrow\quad
\frac{d\lambda_+}{dR}=\frac{\tau'(R)\lambda_+(R)-\delta'(R)}{2\lambda_+(R)-\tau(R)}.
\]
\noindent Since $p(1)$ is $R$–invariant, $-\tau'(R)+\delta'(R)=0$, hence $\delta'(R)=\tau'(R)$ and
\begin{equation}\label{eq:pull_to_one_general_explicit_refined}
\frac{d\lambda_+}{dR}=\frac{\tau'(R)\,\big(\lambda_+(R)-1\big)}{2\lambda_+(R)-\tau(R)}.
\end{equation}
\noindent For a $2\times2$ matrix, $2\lambda_+-\tau=\lambda_+-\lambda_->0$, so $\operatorname{sign}\!\big(d\lambda_+/dR\big)=\operatorname{sign}\!\big(\tau'(R)\big)\operatorname{sign}\!\big(\lambda_+-1\big)$. It remains to compute $\tau'(R)$.
\begin{itemize}
    \item \emph{Haploids.} From \eqref{eq:F_hap},
    \[
    \tau_{\mathrm{hap}}(R)=\frac{1}{1-u_1}\!\left[(1-u_2)\!\left(1-\frac{u_1R(1-s)}{s(1-u_1)}\right)+(1-s)-(1-u_2)\frac{(s-u_1)R(1-s)}{s(1-u_1)}\right],
    \]
    so
    \begin{equation}\label{eq:tauprime_hap}
    \tau'_{\mathrm{hap}}(R)=-\,\frac{(1-u_2)(1-s)}{s(1-u_1)^2}\;<\;0
    \qquad\text{for }u_1,u_2\in(0,1),\ s\in(0,1).
    \end{equation}
    
    \item \textit{Diploids.} From \eqref{eq:F_dip}, the trace is
    \[
    \begin{aligned}
      \tau_{\mathrm{dip}}(R)
      &= \frac{1}{(1-u_1)s} \biggl[
          (1-u_2)\bigl(s - u_1 R(1-s)\bigr) \\
      &\quad
          + (1-s)\bigl((1-R) + u_2 R\bigr)\bigl(s - u_1(1-s)\bigr)
          + (1-2s)u_1
        \biggr].
    \end{aligned}
    \]
    hence
    \begin{equation}\label{eq:tauprime_dip}
    \tau'_{\mathrm{dip}}(R)=-\,\frac{(1-u_2)(1-s)}{\,1-u_1\,}\;<\;0
    \qquad\text{for }u_1,u_2\in(0,1),\ s\in(0,1).
    \end{equation}
\end{itemize}

\noindent Substituting \eqref{eq:tauprime_hap}–\eqref{eq:tauprime_dip} into \eqref{eq:pull_to_one_general_explicit_refined} yields
\begin{equation}\label{eq:pull_to_one_sign_final_refined}
\operatorname{sign}\!\Big(\tfrac{d\lambda_+}{dR}\Big)=\operatorname{sign}\!\big(1-\lambda_+(R)\big).
\end{equation}
Thus, recombination shifts $\lambda_+$ monotonically toward $1$: it decreases $\lambda_+$ when $\lambda_+>1$ and increases it when $\lambda_+<1$. If $u_2=u_1$, then $\lambda_+(R)\equiv 1$ for all $R$, consistent with \eqref{eq:pull_to_one_general_explicit_refined}.


\subsection{\textbf{Stochastic Mutation Rate ($u_{2, t}$).}} \label{sec:invasion_stochastic_unified}
\noindent Assume the resident population is at mutation–selection equilibrium (Eq.~\eqref{eq:mut_sel_balance}) with forward mutation rate $u_1$, satisfying $0<u_1<s$ in haploids and $s>\tfrac{u_1}{1+u_1}$ in diploids. Let the invading modifier allele $M_2$ produce a forward mutation rate that varies independently across generations,
\[
u_{2,t}\stackrel{\text{i.i.d.}}{\sim}\mathcal D(u_2,\sigma^2),\qquad 
\mathbb E[u_{2,t}]=u_2,\quad \mathrm{Var}(u_{2,t})=\sigma^2.
\]
\noindent Linearization at $\hat{\mathbf x}=(\hat x_1,\hat x_2,0,0)^\top$ yields the rare–haplotype recursion $\bm v_{t+1}=\mathbf F_t(R,u_{2,t})\,\bm v_t$ with $\bm v_t=(x_{3,t},x_{4,t})^\top$ and $\mathbf F_t$ given by \eqref{eq:F_hap} or \eqref{eq:F_dip} for the realized $u_{2,t}$ and constant $R\in[0,\tfrac12]$. If $\mathbb E[\log\!\|\mathbf F_t\|]<\infty$, the top Lyapunov exponent
\[
\gamma(R)=\lim_{t\to\infty}\frac{1}{t}\log\big\|\mathbf F_{t-1}(R,u_{2,t-1})\cdots \mathbf F_0(R,u_{2,0})\big\|
\]
exists almost surely and is norm–independent \citep{furstenberg1960}. Invasion occurs if $\gamma(R)>0$ (conversely, $M_2$ is lost if $\gamma(R)<0$; when $\gamma(R)=0$, we cannot distinguish loss from invasion to linear order).

\noindent For each $t$, decompose $\mathbf F_t(R)$ as $\mathbf F_t(R)=\mathbf A_t+R\,\mathbf B_t$, i.e. separate the $R$–independent and $R$–linear parts. Rewrite \eqref{eq:F_hap}–\eqref{eq:F_dip} as,
\begin{equation}\tag{\ref{eq:F_hap}a} \label{eq: A_hap}
\mathbf A_t^{\mathrm{hap}}=\frac{1}{1-u_1}
\begin{pmatrix}
1-u_{2,t} & 0\\[3pt]
u_{2,t} & 1-s
\end{pmatrix},\qquad
\mathbf B_t^{\mathrm{hap}}=\frac{(1-s)(1-u_{2,t})}{s(1-u_1)^2}
\begin{pmatrix}
-u_1 & (s-u_1)\\[8pt]
u_1 & -(s-u_1)
\end{pmatrix},
\end{equation}
for haploids, and for diploids,
\[
\mathbf A_t^{\mathrm{dip}}=\frac{1}{(1-u_1)s}
\begin{pmatrix}
(1-u_{2,t})\,s & 0\\[3pt]
u_{2,t}\,s & (1-s)\,[s-u_1(1-s)]+(1-2s)u_1
\end{pmatrix},
\]
\begin{equation}\tag{\ref{eq:F_dip}a} \label{eq: B_dip}
\mathbf B_t^{\mathrm{dip}}=\frac{(1-s)(1-u_{2,t})}{s(1-u_1)}
\begin{pmatrix}
-u_1 & \,[s-u_1(1-s)]\\[6pt]
u_1 & -\,[s-u_1(1-s)]
\end{pmatrix}.
\end{equation}

\noindent The form of $\mathbf A_t$ in \eqref{eq: A_hap} and \eqref{eq: B_dip} match $\mathbf F_t(0)$ from the $R=0$ analysis. The matrices $\mathbf B_t$ capture the same–generation linear response to recombination; in particular, their column sums are zero, reflecting that recombination redistributes haplotypes across backgrounds. Two implications follow directly:
\begin{enumerate}
\item In the case $R=0$, $\mathbf F_t(0)=\mathbf A_t$ is lower triangular; the Lyapunov exponent reduces to the time average of the log of the dominant diagonal term.
\item For $R>0$ and $u_{2,t}>0$, all entries of $\mathbf F_t(R)$ are strictly positive. Hence each $\mathbf F_t(R)$ is primitive. If $\mathbb P(u_{2,t}>0)>0$, primitivity occurs infinitely often almost surely, which suffices for a unique top Lyapunov exponent \cite{furstenberg1960}. $R$ enters \emph{linearly} at each generation via $\mathbf B_t$, but the map $R\mapsto\gamma(R)$ is \emph{not} necessarily linear (or even monotone) because $\gamma(R)$ is a limit of logs of products of non-commuting random matrices.
\end{enumerate}

\noindent \paragraph{\textbf{Baseline at $R=0$.}}  When $R=0$, $\mathbf F_t(0)$ matrices are lower triangular, so products remain triangular. The top Lyapunov exponent is the maximum of the time–averaged logarithms of the diagonal entries.  
\begin{itemize}
    \item For haploids,
    
    $$\mathbf F_t(0)=\frac{1}{1-u_1}
    \begin{pmatrix}
    1-u_{2,t} & 0\\[3pt]
    u_{2,t} & 1-s
    \end{pmatrix},$$
    $$\gamma(0)=-\log(1-u_1)+\max\!\Big\{\mathbb E[\log(1-u_{2,t})],\,\log(1-s)\Big\}.$$
    
    \item For diploids,
    \[
    \mathbf F_t(0)=\frac{1}{1-u_1}
    \begin{pmatrix}
    1-u_{2,t} & 0\\[3pt]
    u_{2,t} & 1-s(1+u_1)
    \end{pmatrix},
    \]
    and, provided $1-s(1+u_1)>0$,
    \[
    \gamma(0)=-\log(1-u_1)+\max\!\Big\{\mathbb E[\log(1-u_{2,t})],\,\log(1-s(1+u_1))\Big\}.
    \]
    \end{itemize}
\noindent The eigenvalues of $\mathbf F_t(0)$ are therefore
\[
\lambda_1(t)=\frac{1-u_{2,t}}{1-u_1},\qquad
\lambda_2=\begin{cases}
\dfrac{1-s}{1-u_1}, & \text{haploid},\\[6pt]
\dfrac{1-s(1+u_1)}{1-u_1}, & \text{diploid}.
\end{cases}
\]
\noindent \emph{Regime 1 ($\lambda_1(t)$ term dominates).} Because the resident population is at mutation–selection balance, $\lambda_2<1$, in both haploids and diploids. Hence, with complete linkage ($R=0$), invasion can occur only through $\lambda_1(t)$. The stochastic invasion condition is 
\begin{equation}\label{eq:stochastic_R0_inv_condition}
\gamma(0)>0
\;\Longleftrightarrow\;
\mathbb{E}\!\left[\log \lambda_1(t)\right]>0
\;\Longleftrightarrow\;
\mathbb{E}[\log(1-u_{2,t})]>\log(1-u_1).
\end{equation}
\noindent Equation~\eqref{eq:stochastic_R0_inv_condition} states that $M_2$ increases when the expected log of its per-generation probability of transmitting allele $A$, namely $(1-u_{2,t})$, exceeds that of the resident. Because $\log(1-x)$ is strictly concave on $[0,1)$, Jensen’s inequality implies
\[
\mathbb{E}[\log(1-u_{2,t})]\le \log\!\big(1-\mathbb{E}[u_{2,t}]\big),
\]
with equality only if $u_{2,t}$ is constant. Hence temporal variance in $u_{2,t}$ reduces $\mathbb{E}[\log(1-u_{2,t})]$ relative to a deterministic rate with the same mean, shrinking the parameter region in which invasion is possible. Increasing $u_1$ increases $\gamma(0)$  (since $-\log(1-u_1)$ increases), expanding the invasion region. When \eqref{eq:stochastic_R0_inv_condition} holds, the growth rate is
\[
\gamma(0)=\mathbb{E}[\log(1-u_{2,t})]-\log(1-u_1),
\]
which is independent of $s$ and ploidy; invasion then depends solely on the difference in long-run effective mutation rates. The $R{=}0$ invasion criterion can be evaluated analytically or by a single numerical integral for standard distributions; Appendix~\ref{appendix:approximation} and Table~\ref{tab:lyap_G_summary_landscape}, summarize $\mathbb E[\log(1-u_{2,t})]$ and the relationship between $\mathbb E[\log(1-u_{2,t})]$ and $\log(1-u_1)$ for several cases.

\noindent \emph{Regime 2 ($\lambda_2$ term dominates).} If $\mathbb{E}[\log(1-u_{2,t})]\le \log(1-u_1)$, then $\gamma(0)=\log\lambda_2<0$ and invasion fails. In this regime $\gamma(0)$ is independent of $u_{2,t}$ and its variance; only $u_1$, $s$, and ploidy matter.
\begin{itemize}
    \item \textit{Haploids:} $\lambda_2=(1-s)/(1-u_1)$. For fixed $u_1$, $\partial \lambda_2/\partial s=-1/(1-u_1)<0$, so increasing $s$ decreases $\lambda_2$ and makes $\log \lambda_2$ more negative. For fixed $s<1$, $\partial \lambda_2/\partial u_1=(1-s)/(1-u_1)^2>0$, so increasing $u_1$ increases $\lambda_2$ (moving $\log \lambda_2$ toward $0$).
    \item \textit{Diploids:} $\lambda_2=\{1-s(1+u_1)\}/(1-u_1)$, assumed positive (i.e., $1-s(1+u_1)>0$) so that $\log \lambda_2$ is defined. For fixed $u_1$, $\partial \lambda_2/\partial s=-(1+u_1)/(1-u_1)<0$, so increasing $s$ decreases $\lambda_2$. Also,
    \[
    \frac{\partial \lambda_2}{\partial u_1}
    =\frac{1-2s}{(1-u_1)^2},
    \]
$\lambda_2$ increases with $u_1$ when $s<\tfrac12$ and decreases with $u_1$ when $s>\tfrac12$, and is locally insensitive to $u_1$ at $s=\tfrac12$ (subject to $1-s(1+u_1)>0$). Accordingly, $\log \lambda_2$ becomes less negative or more negative in these cases.
\end{itemize}

\noindent In summary, when the $\lambda_1$ term dominates, invasion is determined by differences in long-run effective mutation rates: it is promoted by higher $u_1$ but hindered by temporal variance in $u_{2,t}$. When the $\lambda_2$ term dominates, invasion cannot occur, and the negative Lyapunov exponent $\gamma(0)=\log\lambda_2<0$ quantifies the strength of this constraint on the growth rate of $M_2$'s frequency. Stronger selection decreases $\lambda_2$, making $\gamma(0)$ more negative and thus reinforcing the constraint. The effect of the resident mutation rate $u_1$ is monotonic in haploids but depends on $s$ in diploids, decreasing $\lambda_2$ when $s>\tfrac{1}{2}$ and increasing it when $s<\frac12$.


\noindent \subparagraph{Simulation Results.} We examine how temporal variation in the mutation rate $u_2$ alters invasion when its mean is held constant and $R=0$. The rare modifier allele generates an i.i.d.\ sequence $u_{2,t}\sim\mathcal D(u_2,\sigma^2)$ with mean $\mathbb E[u_{2,t}]=u_2$ and variance $\operatorname{Var}(u_{2,t})=\sigma^2$, supported between $0$ and $1$. Four distributions are considered for $\mathcal D$—uniform, beta, truncated log-normal, and truncated gamma—with parameters fitted to match $(u_2,\sigma^2)$ exactly (Appendix~\ref{appendix:approximation}).

\noindent In the deterministic case ($u_{2,t}\equiv u_2$), the modifier invades if $u_2<u_1$. With stochastic variation in $u_2$ and complete linkage ($R=0$), the leading Lyapunov exponent satisfies
\[
\gamma(0)\;=\;\log\!\frac{1}{1-u_1}\;+\;\mathbb E\big[\log(1-u_{2,t})\big],
\]
so invasion occurs if $\mathbb E[\log(1-u_{2,t})]>\log(1-u_1)$ (Eq.~\eqref{eq:stochastic_R0_inv_condition}). Because $\log(1-x)$ is strictly concave on $[0,1)$,
\[
\mathbb E[\log(1-u_{2,t})]\;\le\;\log\!\big(1-\mathbb E[u_{2,t}]\big)=\log(1-u_2),
\]
with strict inequality when $\sigma^2>0$ (by Jensen’s inequality). Temporal variability in $u_{2,t}$ therefore lowers $\gamma(0)$ relative to the deterministic case, with the reduction driven mainly by infrequent large values of $u_{2,t}$, which strongly affect the logarithmic mean. Figure~\ref{fig:distribution_comparison} shows that, for a constant mean mutation rate $\mathbb{E}[u_{2,t}] = u_2 = 0.06$, increasing the variance $\sigma^2$ changes the shape of $u_{2,t}$ under different distributional assumptions.

\noindent The distributions are parameterized to have the same mean but different higher moments. The uniform expands symmetrically with variance, while the beta shifts probability mass toward both boundaries—especially near $u_{2,t}=1$—as $\sigma^2$ increases. The truncated log-normal and gamma distributions are right-skewed; the former retains a relatively thin tail after truncation, whereas the latter has a heavier upper tail with comparable variance. These differences in shape determine how much weight is placed on infrequent, large $u_{2,t}$ values that disproportionately reduce $\mathbb E[\log(1-u_{2,t})]$ making invasion less likely.

\begin{figure}[h]
    \centering
    \includegraphics[width=\textwidth]{figures/f2_distributions.png}
    \caption{\textbf{Mutation–rate probability densities on $[0,1]$ with fixed mean $u_2=0.06$.}
    Columns show distributions (uniform, beta, truncated log-normal, truncated gamma); rows correspond to “low’’ (top) and “high’’ (bottom) variance parameterizations. Dashed vertical line: common mean. Parameters used: 
    uniform $[a,b]=[0.045,0.075]$ (low), $[0.001,0.119]$ (high); 
    beta $(\alpha,\beta)=(12.0,188.0)$ (low), $(0.15,2.35)$ (high);
    log-normal $(\mu_{\ln},\sigma_{\ln})=(-3.0,0.25)$ (low), $(-3.8,0.83)$ (high); 
    gamma $(k,\theta)=(4.0,0.015)$ (low), $(2.0,0.030)$ (high).
    Log-normal and gamma densities are truncated and renormalized on $[0,1]$; realized variances (annotated in each panel) are computed numerically.}
    \label{fig:distribution_comparison}
\end{figure}
\noindent For each simulation, the population is initialized at the resident equilibrium under \(M_1\) (haploid example: \(u_1\approx 0.048796\)), set \(s=0.2\) and \(R=0\), and the full two-locus recursion is iterated for \(5{,}000\) generations. Fixation is declared if \((x_3+x_4)_t\ge 0.999\) (modifier \(M_2\)) or \((x_1+x_2)_t\ge 0.999\) (resident \(M_1\)); otherwise the outcome is “no fixation.” For each distributional family \(\mathcal D\) we generate an i.i.d.\ sequence \(u_{2,t}\sim\mathcal D(u_2,\sigma^2)\) with prescribed mean \(u_2\) and variance \(\sigma^2\).

\noindent Parameterization is as follows. (i) \emph{uniform:} choose \([a,b]\) with \(a=u_2-\sqrt{3\sigma^2}\), \(b=u_2+\sqrt{3\sigma^2}\), and \(\sigma^2=(b-a)^2/12\). (ii) \emph{beta:} set \(\alpha=u_2 m\), \(\beta=(1-u_2)m\) with \(m=u_2(1-u_2)/\sigma^2-1\) (feasible for \(\sigma^2<u_2(1-u_2)\)); this matches \((u_2,\sigma^2)\) exactly on \([0,1]\). (iii) \emph{truncated log-normal} and (iv) \emph{truncated gamma:} choose \((\mu_{\ln},\sigma_{\ln})\) or \((k,\theta)\) so that the \([0,1]\)-truncated density has mean \(u_2\) and variance \(\sigma^2\); parameters are obtained by a two-dimensional root-finding routine, and the realized \(\sigma^2\) is computed by numerical quadrature. The feasible variance range \([\min\sigma^2,\max\sigma^2]\) depends on the family: beta admits the largest \(\sigma^2\) (probability near boundaries), uniform is limited by interval width, and truncation restricts log-normal and gamma to have moderate \(\sigma^2\).

\noindent Figure~\ref{fig:distribution-fixation-all} summarizes simulation outcomes in terms of \((u_2,\sigma^2)\) for each \(\mathcal D\). Boundaries align with the geometric-mean criterion at \(R=0\): distributions placing little mass at high mutation (uniform; truncated log-normal) maintain larger \(\mathbb E[\log(1-u_{2,t})]\) and favor invasion, whereas distributions allowing frequent large \(u_{2,t}\) (beta; high-variance truncated gamma) reduce the logarithmic mean and disfavor invasion—even when the arithmetic mean \(u_2<u_1\). Quantitative comparisons across distributions are provided in Appendix~\ref{app: compare_derivations}, Table~\ref{tab:u2star_comparison}, which reports how the invasion boundary $u_2^*$ varies with constant variance for different distributions. These results illustrate how the location of probability mass relative to $u_1$ affects invasion through its effect on $\mathbb E[\log(1-u_{2,t})]$.

\begin{figure}[h]
    \centering
    \includegraphics[width=\linewidth]{figures/f3_haploid_simulation_results.png}
    \caption{\textbf{Fixation outcomes for alleles $M_1$ and \(M_2\) across \((u_2,\sigma^2)\) at \(R=0\) (haploids).} 
    Settings: \(5{,}000\) generations; \(M_2\) initially rare; resident \(M_1\) fixed at \(u_1=0.048796\); \(s=0.2\). Columns correspond to distributional families (uniform, beta, truncated log-normal, truncated gamma). The variance axis in each panel reflects the feasible range at constant \(u_2\): uniform \(\sigma^2\in[0,\,2\times10^{-3}]\), beta \([0,\,5\times10^{-2}]\), truncated log-normal \([0,\,2\times10^{-3}]\), truncated gamma \([0,\,10^{-2}]\) (panel limits used in the simulations). Parameterization for each sampled point: uniform \([a,b]=[u_2-\sqrt{3\sigma^2},\,u_2+\sqrt{3\sigma^2}]\) clipped to \([0,1]\); beta \((\alpha,\beta)=(u_2 m,\,(1-u_2)m)\) with \(m=u_2(1-u_2)/\sigma^2-1\); truncated log-normal and truncated gamma fitted so their truncated moments equal \((u_2,\sigma^2)\) (realized \(\sigma^2\) obtained by numerical quadrature). Colors denote outcomes: \(M_1\) fixation (blue), \(M_2\) fixation (orange), no fixation within $5{,}000$ generations (gray).}
    \label{fig:distribution-fixation-all}
\end{figure}
\begin{table}[h]
\centering
\renewcommand{\arraystretch}{1.12}
\begin{tabular}{p{0.23\textwidth} p{0.34\textwidth} p{0.34\textwidth}}
\hline
\textbf{Distribution} & \textbf{Variance–shape relationship at constant $u_2$ (Fig.~\ref{fig:distribution_comparison})} & \textbf{Fixation pattern (Fig.~\ref{fig:distribution-fixation-all})} \\
\hline
Uniform & Flat density; increasing variance widens the support symmetrically around the mean.& Nearly horizontal fixation boundary; weak dependence on $\sigma^2$. $M_2$ invasion remains likely whenever $u_2<u_1$. \\[3pt]
Beta & Variance increases by shifting probability mass toward both boundaries, especially near $u_{2,t}=1$, producing a heavy upper tail. & Strong sensitivity to $\sigma^2$: the $M_2$ fixation region contracts rapidly with increasing variance, and $M_1$ frequently fixes even when $u_2<u_1$. \\[3pt]
Truncated log-normal & Right-skewed with most mass concentrated near $u_{2,t}=0$; truncation sharply limits the upper tail. & $M_2$ invasion robust to stochasticity; the $M_2$ fixation region declines only gradually as $\sigma^2$ rises. \\[3pt]
Truncated gamma & Moderately right-skewed with greater weight at intermediate-to-large values than the log-normal; truncation reduces extreme tails but retains moderate upper mass. & Intermediate sensitivity: $M_2$ invades for small $\sigma^2$ but its fixation region contracts steadily as $\sigma^2$ grows. \\[3pt]
\hline
\end{tabular}
\caption{\textbf{Summary of distributional effects on modifier invasion at $R=0$.} 
The shape of the mutation–rate distribution determines $\mathbb E[\log(1-u_{2,t})]$ and thus the long-term growth rate of the invading modifier. 
Distributions placing more probability mass near high mutation values ($u_{2,t}\!\to\!1$) yield smaller geometric means and therefore hinder invasion, even when the arithmetic mean $u_2$ is held constant. 
Conversely, distributions concentrated at low $u_{2,t}$ maintain larger $\mathbb E[\log(1-u_{2,t})]$ and favor invasion.}
\label{tab:distribution-summary}
\end{table}
\noindent These results highlight two key insights. First, invasion depends on $\mathbb{E}[\log(1 - u_{2,t})]$ rather than on $\mathbb{E}[u_{2,t}]$. For a constant mean of $u_{2,t}$, greater variance $\sigma^2$ decreases $\mathbb{E}[\log(1 - u_{2,t})]$. More generally, when two distributions with the same mean are ordered by convexity (i.e., one is more variable in the sense of second-order stochastic dominance), the expectation of the concave function $\log(1 - x)$ is smaller for the more variable distribution. Consequently, modifier alleles with identical mean mutation rates may differ in their invasion outcomes solely due to differences in temporal variance. That is, if $u_{2,t}^{(A)}$ and $u_{2,t}^{(B)}$ are two i.i.d.\ sequences with $\mathbb E[u_{2,t}^{(A)}]=\mathbb E[u_{2,t}^{(B)}]=u_2$ and
\[
\mathbb E\!\big[\log(1-u_{2,t}^{(A)})\big]\;>\;\log(1-u_1)\;\ge\;\mathbb E\!\big[\log(1-u_{2,t}^{(B)})\big],
\]
then $M_2^{(A)}$ invades at $R=0$ while $M_2^{(B)}$ does not, despite identical means. 

\noindent \paragraph{\textbf{Stochastic mutation rate with Recombination $R\in(0,\tfrac12]$.}} For $R>0$, $\mathbf F_t(R)$ has positive off–diagonal entries whenever $u_{2,t}>0$; products of $\mathbf F_t(R)$ do not commute, and no closed form for $\gamma(R)$ is available. However, the qualitative effect of $R$ derives from the linear structure of \eqref{eq: A_hap} and \eqref{eq: B_dip}: recombination has no direct fitness effect at the modifier alleles, it mediates the indirect response by redistributing the associations between $M_2$ and the selected background that are generated each generation by selection and mutation. 

\noindent In deterministic settings (constant $u_2$), this association brings the dominant eigenvalue of the linearized near $(\hat x_1, \hat x_2, 0, 0)$ towards $1$, changing the \emph{magnitude} but not the \emph{sign} of initial frequency change.  With stochastic $u_{2,t}$, multiplicative averaging emphasizes the geometric mean of per–generation growth factors, and the distribution of $u_{2,t}$ interacts with $s$ and $u_1$, allowing recombination to \emph{reverse} the sign of $\gamma(R)$ as $R$ increases—an effect not seen in deterministic models.

\noindent\subparagraph{Numerical analysis.} To assess the effect of recombination with stochastically varying mutation induced by $M_2$, let \(u_{2,t}\stackrel{\text{i.i.d.}}{\sim}\mathrm{Beta}(\alpha,\beta)\) with mean \(u_2=0.04\) and variance $\sigma^2$ $\in$ $\{0,\,0.25,\,0.5,\,0.75,\,0.95\}$ $\times u_2(1-u_2)$, where \(u_2(1-u_2)=0.0384\). The factor \(u_2(1-u_2)\) is the maximum possible variance for a variable on \([0,1]\) with mean \(u_2\) and scaling by $u_2(1-u_2)$ expresses \(\sigma^2\) as a fraction of this upper bound for comparability. For each \(\sigma^2\), we draw one sequence \(\{u_{2,t}\}\), and reuse it for both ploidies to isolate ploidy effects. For \(R\) on a grid in \([0,\tfrac12]\), we iterate \(\bm v_{t+1}=\mathbf F_t(R)\,\bm v_t\) with \(\ell^1\)-normalization at each step; by the Furstenberg–Kesten theorem, the time average of the log rescalings converges almost surely to the top Lyapunov exponent \(\gamma(R)\)~\cite{furstenberg1960}. Unless stated otherwise, \(u_1=0.05\) and \(s\in\{0.20,\,0.06\}\). Figure~\ref{fig:gamma-lyap} plots \(\gamma(R)\) for both ploidies; the left and right endpoints of each curve correspond to \(R=0\) and \(R=\tfrac12\), respectively.
\begin{itemize}
    \item \textit{Weak selection (\(s=0.06\)).} With low variance (\(0.25\times u_2(1-u_2)\)), \(\gamma(0)>0\) while \(\gamma(\tfrac12)<0\) (e.g., haploid: \(\gamma(0)\approx 8\times10^{-4}\to \gamma(\frac12)\approx-2\times10^{-4}\); diploid: \(\gamma(0)\approx 8\times10^{-4}\to \gamma(\frac12)\approx-1\times10^{-4}\)). Thus if recombination is frequent enough, it suppresses invasion.
    \item     \textit{Strong selection (\(s=0.20\)).} With high variance (\(0.95\times u_2(1-u_2)\)), \(\gamma(0)<0\) while \(\gamma(\tfrac12)>0\) in both ploidies (e.g., haploid: \(\gamma(0)\approx-1.72\times10^{-1}\to \gamma(\frac12)=2\times10^{-4}\); diploid: \(\gamma(0)\approx-1.85\times10^{-1}\to \gamma(\frac12)=1\times10^{-4}\)). Thus sufficiently frequent recombination allows invasion.
\end{itemize}
\noindent Fig.~\ref{fig:gamma-lyap} shows that sign changes in $\gamma(R)$ with $R$ occur only when variability in $u_{2,t}$ is present; with $\sigma^2=0$ (deterministic $u_{2,t}\equiv u_2$), $R$ changes the magnitude but not the sign of $\gamma(R)$ on $[0,\tfrac12]$.

\noindent \emph{Recombination cutoff.} Define the cutoff \(R^\ast:=\inf\{\,R\in[0,\tfrac12]:\gamma(R)=0\,\}\), the smallest recombination fraction at which the long–run growth rate of \(M_2\) changes invasion direction ($\gamma(R) \lessgtr 0$). This quantity identifies the minimum recombination needed to reverse the direction of invasion for a given parameter set. We estimate \(R^\ast\) by evaluating \(\gamma(R)\) on a fine grid in \(R\in[0,\tfrac12]\) using a common realization of \(\{u_{2,t}\}\) for all \(R\) (variance reduction via common random numbers). We then bracket a sign change, selecting an interval \([R_L,R_U]\) with \(\gamma(R_L)\gamma(R_U)<0\), and solve \(\gamma(R)=0\) on \([R_L,R_U]\) by a one–dimensional root finder (bisection or Brent). Tolerances are \(|\gamma(R^\ast)|<10^{-5}\) or interval width \(<10^{-3}\). Results are stable to numerical error across independent replicates to numerical error.
\begin{itemize}
\item \textit{Weak selection example.} For \(s=0.06\) and a beta distribution for \(u_{2,t}\) calibrated to mean \(u_2\) and variance \(\sigma^2=0.25 \times \,u_2(1-u_2)\), we obtain
\[
R^\ast \approx 0.27 \ \text{(haploid)},\qquad R^\ast \approx 0.28 \ \text{(diploid)},
\]
consistent with the zero crossing near \(R\in[0.25,0.30]\) in Fig.~\ref{fig:gamma-lyap}. In this setting, moderate recombination reverses the sign of \(\gamma(R)\) because it disrupts associations between \(M_2\) and high–mutation backgrounds that disproportionately reduces the geometric mean growth rate. When the variance is increased (e.g., \(0.50\times u_2(1-u_2)\) and \(0.75 \times u_2(1-u_2)\) for the same beta distribution), \(\gamma(R)\) can become non-monotone in \(R\). Intuitively, for small \(R\), recombination primarily breaks associations formed in generations with larger \(u_{2,t}\), raising \(\gamma(R)\); at higher \(R\), it also breaks beneficial associations formed during low–\(u_{2,t}\) generations, lowering \(\gamma(R)\). This can yield two zero crossings in \(R\); by definition, \(R^\ast\) denotes the smaller root. The presence and locations of these roots depend on \((s,u_1)\) and the full distribution of \(u_{2,t}\) (not just its mean and variance).
\item \textit{Strong selection example.} For \(s=0.20\) and high variance \(\sigma^2=0.95 \times \,u_2(1-u_2)\),
\[
R^\ast \approx 0.46 \ \text{(haploid)},\qquad R^\ast \approx 0.49 \ \text{(diploid)}.
\]
Here, reversal requires recombination to be close to free (\(R=\tfrac12\)): strong selection amplifies the cost of low-fitness associations, so more frequent reshuffling is needed to offset them. In these high-variance cases, \(\gamma(R)\) typically increases with \(R\) for small \(R\) (breaking high–\(u_{2,t}\) associations) and may decrease for larger \(R\) once the loss of favorable associations during low–\(u_{2,t}\) episodes dominates.
\end{itemize}
\noindent The above variance multipliers were chosen because, among the tested set \(\{0,0.25,0.5,0.75,0.95\}\), they were the lowest and highest values for which \(\gamma(R)\) changed sign under weak and strong selection, respectively.

\noindent \emph{Parameter effects.} For $R>0$, recombination affects associations between the modifier and the selected background without directly changing modifier fitness. Consequently, the sensitivity of $\gamma(R)$ to $R$ is governed by (i) how selection strength $s$ and the resident mutation rate $u_1$ are associated at each generation, and (ii) how temporal variability in $u_{2,t}$ enters through geometric (multiplicative) averaging.
\noindent Throughout, we use the linear decomposition $\mathbf F_t(R)=\mathbf A_t+R\,\mathbf B_t$ with
\begin{equation}\label{eq:A_hap_repeat}
\mathbf A_t^{\mathrm{hap}}=\frac{1}{1-u_1}
\begin{pmatrix}
1-u_{2,t} & 0\\[3pt]
u_{2,t} & 1-s
\end{pmatrix},
\qquad
\mathbf B_t^{\mathrm{hap}}=\frac{(1-s)(1-u_{2,t})}{s(1-u_1)^2}
\begin{pmatrix}
-\,u_1 & \;s-u_1\\[6pt]
u_1 & -\,(s-u_1)
\end{pmatrix},
\end{equation}
for haploids, and
\begin{equation}\label{eq:A_B_dip_repeat}
\mathbf A_t^{\mathrm{dip}}=\frac{1}{1-u_1}
\begin{pmatrix}
1-u_{2,t} & 0\\[3pt]
u_{2,t} & 1-s(1+u_1)
\end{pmatrix},
\qquad
\mathbf B_t^{\mathrm{dip}}=\frac{(1-s)(1-u_{2,t})}{s(1-u_1)}
\begin{pmatrix}
-\,u_1 & \;s-u_1(1-s)\\[6pt]
u_1 & -\,[\,s-u_1(1-s)\,]
\end{pmatrix},
\end{equation}
for diploids. These make explicit that $R$ enters \emph{linearly} in each generation via $\mathbf B_t$. However, $\gamma(R)$ itself is not generally a linear (or monotone) function of $R$ because it depends on products of non-commuting random matrices.

\begin{itemize}
\item \textit{Selection strength $s$.} In both ploidies, $\mathbf B_t$ scales as $(1-s)/s$ times a matrix whose off–diagonal elements increase linearly with $s$ (specifically $s-u_1$ in haploids and $s-u_1(1-s)$ in diploids). Because $\mathrm d[(1-s)/s]/\mathrm ds=-1/s^2<0$, increasing $s$ reduces the overall multiplicative factor governing the per–generation sensitivity to $R$, while simultaneously amplifying certain matrix entries that mediate background exchange. The resulting effect of $s$ on the long–run growth rate $\gamma(R)$ is therefore non-monotonic: stronger selection diminishes the effect of recombination yet enhances direct coupling between backgrounds. At $R=0$, where $\mathbf F_t=\mathbf A_t$ is lower–triangular, the eigenvalue $\lambda_2$ equals $1-s$ (haploid) or $1-s(1+u_1)$ (diploid), with $\partial\lambda_2/\partial s<0$, so $\log\lambda_2$ decreases strictly with $s$. For $R>0$, however, $\gamma(R)$ depends on products of non-commuting random matrices, and its response to increasing $s$ cannot be determined in general.

\item \textit{Resident mutation rate $u_1$.} Assume $u_1\in[0,1)$ so that expectations of logs exist. At $R=0$, the product is lower–triangular and $\gamma(0)=\mathbb{E}[\log(1-u_{2,t})]-\log(1-u_1)$, hence
\[
\frac{\partial\gamma(0)}{\partial u_1}=\frac{1}{1-u_1}>0,
\]
so increasing $u_1$ increases $\gamma(0)$. For $R>0$, $u_1$ enters $\mathbf B_t$ through global factors $(1-u_1)^{-2}$ (haploid) or $(1-u_1)^{-1}$ (diploid), which amplify recombination’s per–generation effect as $u_1$ increases, and through off–diagonals $s-u_1$ (haploid) and $s-u_1(1-s)$ (diploid), which \emph{decrease} with $u_1$; these opposing influences, combined with temporal covariances between $u_{2,t}$ and selection–generated modifier–background associations, entail that the effect of $u_1$ on $\gamma(R)$ for $R>0$ has no general sign and is parameter–dependent.

\item \textit{Temporal variability of $u_{2,t}$.}  For constant mean $u_2$, concavity of $\log(1-x)$ on $[0,1)$ gives $\mathbb E[\log(1-u_{2,t})]\le \log\!\big(1-\mathbb E[u_{2,t}]\big)$, with strict inequality when $\mathrm{Var}(u_{2,t})>0$. Thus higher variance lowers the $R{=}0$ component of $\gamma(R)$.  When $R>0$, $u_{2,t}$ enters $\mathbf B_t$ through the factor $(1-u_{2,t})$, so higher temporal variance increases fluctuations in the per–generation growth rate and modifies how recombination influences the geometric mean. The overall influence of variability on $\gamma(R)$ depends on the joint dynamics of $u_{2,t}$ and the modifier–background associations and cannot be determined in general.

\item \textit{Comparing ploidies.}  The recombination prefactor is $(1-u_1)^{-2}$ in haploids and $(1-u_1)^{-1}$ in diploids, so—holding other terms fixed—the single–step $R$–perturbation is larger in haploids. However, the $s$–dependent off–diagonal terms also differ (e.g., $s-u_1$ vs.\ $s-u_1(1-s)$), and $\lambda_2$ differs between ploidies at $R=0$. Consequently, there is no uniform ordering by ploidy; differences in $\gamma(R)$ arise from the interaction of these factors with the distribution of $u_{2,t}$.
\end{itemize}

\begin{figure}[t]
  \centering
  \includegraphics[width=1.1
\linewidth]{figures/f1_gamma_lyap_composite.png}
  \caption{Top Lyapunov exponent \(\gamma(R)\) for the two-locus modifier system. Panels are arranged in a \(2\times2\) grid: rows indicate selection strength \(s\in\{0.06,\,0.20\}\); columns indicate ploidy (haploid vs.\ diploid). For each \((s,\text{ploidy})\), curves correspond to variances \(\sigma^2\in\{0,\,0.25,\,0.50,\,0.75,\,0.95\}\times u_2(1-u_2)\) with \(u_2=0.04\). The deterministic baseline (\(\sigma^2=0\), labeled \emph{\(u_{2,t}=u_2\)}) uses \(\gamma(R)=\log\rho\!\big(\mathbf F(u_2,R)\big)\). Stochastic curves plot the sample Lyapunov exponent from normalized matrix products using the vector \(\ell^1\) norm at each iteration (i.e., \(\|\cdot\|_1\) re-scaling). Annotated points display \(\gamma(0)\) and \(\gamma(\tfrac12)\) only for selected variance levels: in the weak-selection panels (\(s=0.06\)), annotations are shown for \(\sigma^2=0\) and \(\sigma^2=0.25\,u_2(1-u_2)\); in the strong-selection panels (\(s=0.20\)), for \(\sigma^2=0\) and \(\sigma^2=0.95\,u_2(1-u_2)\).}
  \label{fig:gamma-lyap}
\end{figure}

\noindent Stochastic transmission replaces arithmetic by geometric averaging of per–generation growth factors, so invasion depends on the \emph{full distribution} of $u_{2,t}$ rather than only its mean. At $R=0$, the condition $\gamma(0)>0$ is equivalent to $\mathbb E[\log(1-u_{2,t})]>\log(1-u_1)$; by concavity of $\log(1-x)$, greater variance in $u_{2,t}$ strictly lowers $\mathbb E[\log(1-u_{2,t})]$ at constant mean, reducing $\gamma(0)$. For $R>0$, recombination has no direct fitness effect at the modifier, but it affects the time–varying associations between $M_1/M_1$ and $A/a$ generated each generation by selection and mutation. Because $\gamma(R)$ is a Lyapunov exponent of products of random matrices, its dependence on $R$ need not be linear or monotone: recombination can increase $\gamma(R)$ when it disproportionately breaks associations formed during high–$u_{2,t}$ generations, and decrease $\gamma(R)$ when it breaks associations formed during low–$u_{2,t}$ generations. The joint effects of the distribution of $u_{2,t}$ (particularly its variance), the selection coefficient $s$, and the resident mutation rate $u_1$ can therefore induce sign reversals of $\gamma(R)$ that are absent in deterministic models with constant $u_2$. In sum, stochastic mutation at the modifier creates regimes in which recombination alters not only the \emph{magnitude} of the growth rate but also the \emph{direction} of invasion.
\newpage

\section*{Discussion}
\noindent Most analyses of genetic evolution in large populations introduce stochasticity through fluctuating selection while holding transmission parameters fixed. In this study, selection is constant and randomness enters through the transmission process itself. A neutral modifier locus ($M_1/M_2$) changes the forward mutation rate at a linked, selected locus ($A/a$): the resident $M_1$ has constant rate $u_1$, whereas the invader $M_2$ has a generation–to–generation rate $u_{2,t}\in[0,1)$ with mean $u_2$ and variance $\sigma^2$. We ask how $\sigma^2>0$ alters $M_2$ invasion.

\noindent In the deterministic setting ($u_{2,t}\equiv u_2$), both haploid and diploid models yield an invasion condition that is independent of $R$: the modifier $M_2$ increases when rare if $u_2<u_1$, consistent with the \emph{Reduction Principle} (Eq.~\eqref{eq:deterministic_reduction_principle}). Recombination shifts the dominant eigenvalue of the linearized system toward unity, reducing the magnitude of indirect selection without altering its sign; hence, recombination affects the rate but not the direction of invasion.  In this case, the direction of modifier evolution is statistically aligned with the equilibrium structure of the resident system. 

\noindent With stochastic mutation ($\mathrm{Var}(u_{2,t})>0$) and complete linkage ($R=0$), the invasion recursion is lower triangular and the criterion becomes geometric (Eq.~\eqref{eq:stochastic_R0_inv_condition}):
\[
\gamma(0)=-\log(1-u_1)+\mathbb E[\log(1-u_{2,t})],\qquad
\gamma(0)>0\ \Longleftrightarrow\ \mathbb E[\log(1-u_{2,t})]>\log(1-u_1).
\]
\noindent Because $\log(1-x)$ is strictly concave on $[0,1)$, $\mathbb E[\log(1-u_{2,t})]\le \log(1-u_2)$, with strict inequality when $\sigma^2>0$; randomness in $u_{2,t}$ therefore reduces $\gamma(0)$ relative to the deterministic case. Occasional large $u_{2,t}$ values near $1$ can substantially reduce $\mathbb E[\log(1-u_{2,t})]$. Simulations at $R=0$ with uniform, beta, truncated log–normal, and truncated gamma families support this: with constant $u_2$, heavier upper tails yield smaller geometric means and lower invasion probabilities.

\noindent Variability at $R=0$ also modifies the deterministic \emph{Reduction Principle}. When $u_{2,t}\equiv u_2$, any reduction ($u_2<u_1$) gives $\gamma(0)=\log\!\big((1-u_2)/(1-u_1)\big)>0$. If $u_{2,t}$ fluctuates with small variance $\sigma^2$, a Taylor expansion yields
\[
\mathbb E[\log(1-u_{2,t})]=\log(1-u_2)-\frac{\sigma^2}{2(1-u_2)^2}+o(\sigma^2),
\]
so invasion requires
\[
\log\!\Big(\frac{1-u_2}{1-u_1}\Big)>\frac{\sigma^2}{2(1-u_2)^2}.
\]
\noindent Thus, for sufficiently large $\sigma^2$, invasion can fail even when $\mathbb E[u_{2,t}]<u_1$. Variability in transmission can therefore reduce—or overturn—the deterministic \emph{Reduction Principle}.

\noindent For $0<R\le\tfrac12$, $\gamma(R)$ is continuous in $R$ under the regularity conditions assumed for the product of random matrices. At $R=0$, the geometric–mean condition defines the baseline, with $\gamma(0)$ decreasing monotonically in $\mathrm{Var}(u_{2,t})$ with constant mean. When $R>0$, recombination reshuffles haplotypes, averaging fitness effects over the distribution of associations produced each generation by selection and mutation. This averaging can either increase or decrease the Lyapunov exponent: $\gamma(R)$ increases when recombination disproportionately breaks associations formed during high–$u_{2,t}$ generations (which decreases multiplicative growth), and decreases when it breaks associations formed during low–$u_{2,t}$ generations. Consequently, with $\mathrm{Var}(u_{2,t})>0$ the direction of modifier evolution is not necessarily aligned with the deterministic equilibrium and can be reversed by recombination —a phenomenon absent when $u_{2,t}$ is deterministic. The direction of this stochastic reversal depends on $u_1$, $s$, ploidy, and the distribution of $u_{2,t}$; modifier alleles with the same mean $u_2$ can therefore differ in invasion solely because temporal variability in $u_{2,t}$ induces different responses to recombination.

\noindent Two conclusions follow. First, the \emph{Reduction Principle} emerges as the deterministic limit: with constant $u_2$, the invasion condition depends only on the ordering of $u_2$ and $u_1$; under stochastic $u_{2,t}$, the relevant quantity is the geometric mean of $(1-u_{2,t})$, which can diverge markedly from predictions based on $u_2$. Second, recombination neither uniformly facilitates nor uniformly hinders invasion; its effect depends jointly on $u_1$, $s$, ploidy, and the distribution of $u_{2,t}$. Mean mutation rates alone are therefore insufficient to predict outcomes: modifiers alleles with identical $u_2$ can differ in invasion outcome purely because of differences in temporal variance or tail behavior, through their impact on $\mathbb E[\log(1-u_{2,t})]$ and on products of the $R$–dependent $\mathbf F_t$ matrices.

 \noindent Several extensions follow naturally. Introducing weak temporal correlation in $u_{2,t}$ might affect whether clustering of high–mutation generations changes invasion probabilities and how this interacts with small $R>0$. Allowing recombination itself to vary randomly ($R_t\in[0,\tfrac12]$), possibly jointly with $u_{2,t}$, would generalize the framework to products of matrices with time–varying $R_t$; the effect on $\gamma$ would depend on the joint distribution of $(u_{2,t},R_t)$ and their covariation with selection–generated associations. \emph{Empirically}, these results motivate estimating not only $\mathbb E[u_{2,t}]$ and $\mathrm{Var}(u_{2,t})$ but also $\mathbb E[\log(1-u_{2,t})]$ and the temporal covariance between $u_{2,t}$ and measures of linkage or background fitness, using pedigree–based recombination maps, mutation–accumulation lines, or experimental evolution data.

\noindent In summary, stochastic transmission replaces the arithmetic–mean criterion of deterministic models with a geometric–mean criterion. Rare but extreme realizations of $u_{2,t}$ disproportionately reduce the expected logarithmic growth rate, while recombination determines how this variability is averaged across genetic backgrounds. Invasion therefore depends jointly on the distribution of $u_{2,t}$, the recombination rate, selection strength, and ploidy. Stochastic variation in $u_{2,t}$ can reverse the deterministic \emph{Reduction Principle}, allowing recombination to change not only the \emph{magnitude} but also the \emph{direction} of modifier invasion.

\newpage
\appendix
\section{Estimation of Invasion Regions from Lyapunov Exponents: Examples} \label{appendix:approximation}
\noindent With $R=0$, the invasion matrices are lower triangular for both ploidies, so the top Lyapunov exponent equals the larger of the two diagonal log–growth rates. Fix $u_1\in(0,1)$ and a rare modifier allele $M_2$ with i.i.d.\ mutation rates $u_{2,t}\in[0,L]$ ($L\le1$), mean $u_2$ and variance $\sigma^2$. Then
\begin{equation}\label{eq:lyap_clean}
\gamma(0)\;=\;-\log(1-u_1)\;+\;\max\!\Big\{\ \mathbb E\big[\log(1-u_{2,t})\big]\ ,\ \log\Lambda_{\mathrm{res}}\ \Big\},
\end{equation}
where
\[
\Lambda_{\mathrm{res}}=
\begin{cases}
\dfrac{1-s}{\,1-u_1\,},& \text{haploids},\\[6pt]
\dfrac{1-s(1+u_1)}{\,1-u_1\,},& \text{diploids}.
\end{cases}
\]
\noindent Under the resident–equilibrium constraints ($u_1<s$ in haploids; $u_1<s/(1-s)$ in diploids), $\log\Lambda_{\mathrm{res}}<\log(1-u_1)$, and \eqref{eq:lyap_clean} yields the invasion criterion
\begin{equation}\label{eq:inv_unified}
\gamma(0)>0\quad\Longleftrightarrow\quad \mathbb E\!\big[\log(1-u_{2,t})\big]\;>\;\log(1-u_1).
\end{equation}
\noindent For any distribution \(\mathcal D\) on \([0,L]\) with density \(f(u)\), the expected log term is
\[
\mathbb E[\log(1-u_{2,t})]\;=\;\int_0^L \log(1-u)\,f(u)\,du.
\]
\noindent This quantity can be evaluated directly once the distribution is parameterized by its mean \(u_2\) and variance \(\sigma^2\), either in closed form or by one–dimensional numerical integration. The invasion boundary then follows from solving
\[
\mathbb E[\log(1-u_{2,t})]\;=\;\log(1-u_1),
\]
subject to the feasible support of \(\mathcal D\). Table~\ref{tab:lyap_G_summary_landscape} summarizes analytical and numerical expressions for \(\mathbb E[\log(1-u_{2,t})]\) and the corresponding invasion criterion across the four distribution families considered.

\begin{landscape}
\begin{table}[p]
\centering
\small
\renewcommand{\arraystretch}{1.08}
\setlength{\tabcolsep}{5pt}

\vspace{6pt}
\resizebox{\linewidth}{!}{%
\begin{tabular}{>{\centering\arraybackslash}p{0.12\linewidth} >{\centering\arraybackslash}p{0.34\linewidth} >{\centering\arraybackslash}p{0.30\linewidth} >{\centering\arraybackslash}p{0.20\linewidth}}
\toprule
\textbf{Family on $[0,L]$} &
\textbf{Moment parameterization \& feasibility} &
\(\displaystyle \boldsymbol{\mathbb E[\log(1-u)]}\) &
\textbf{Invasion check at $R{=}0$} \\
\midrule

\textbf{Uniform} \(\ \mathrm{Unif}[a,b]\) &
\(u_2=\tfrac{a+b}{2},\ \sigma^2=\tfrac{(b-a)^2}{12}\).
Set \(a=u_2-\sqrt{3\,\sigma^2}\), \(b=u_2+\sqrt{3\,\sigma^2}\);
feasible if \(0\le a<b\le L\).&
\[
\frac{1}{b-a}\!\int_a^b\!\log(1-u)\,du
=\frac{H(b)-H(a)}{b-a},
\]
\(H(x)=(x-1)\log(1-x)-x\). &
Solve \(\mathbb E[\log(1-u)] = \log(1-u_1)\). \\[6pt]

\textbf{Beta} \(\ u=L X,\ X\sim\mathrm{Beta}(\alpha,\beta)\) &
\[
u_2=L\frac{\alpha}{\alpha+\beta},\quad
\sigma^2=L^2\frac{\alpha\beta}{(\alpha+\beta)^2(\alpha+\beta+1)}.
\]
Let \(m=u_2/L\) and \(v=\sigma^2/L^2\).
Then \(\alpha=m\,t,\ \beta=(1-m)\,t,\ t=\frac{m(1-m)}{v}-1\).
Feasible if \(0<u_2<L\) and \(0<\sigma^2<u_2(L-u_2)\). &
\[
\int_{0}^{1}\!\log(1-Lx)\,
\frac{x^{\alpha-1}(1-x)^{\beta-1}}{B(\alpha,\beta)}\,dx,
\]
Where \(B(\alpha,\beta)=\Gamma(\alpha)\Gamma(\beta)/\Gamma(\alpha+\beta)\). &
Map \((u_2,\sigma^2)\!\mapsto\!(\alpha,\beta)\), evaluate \(\mathbb E[\log(1-u)]\), and test \(>\log(1-u_1)\). \\[6pt]

\textbf{Truncated log–normal} \(\ \mathrm{LN}_{(0,L]}(\mu,\tau^2)\), \(\log u\sim\mathcal N(\mu,\tau^2)\) &
Any \((\mu,\tau^2)\) feasible on \((0,L]\).
Given \((u_2,\sigma^2)\), recover \((\mu,\tau^2)\) by matching truncated moments:
$u_2=\frac{\int_0^L u\,f_{\mathrm{LN}}(u;\mu,\tau^2)\,du}{\Phi\!\big(\tfrac{\log L-\mu}{\tau}\big)},$
$\sigma^2=\frac{\int_0^L u^2 f_{\mathrm{LN}}(u;\mu,\tau^2)\,du}{\Phi\!\big(\tfrac{\log L-\mu}{\tau}\big)}-u_2^2,$
where \(f_{\mathrm{LN}}(u)=\frac{1}{u\,\tau\sqrt{2\pi}}e^{-(\log u-\mu)^2/(2\tau^2)}\), \(\Phi=\) standard normal CDF. &
\[
\frac{1}{\Phi\!\big(\tfrac{\log L-\mu}{\tau}\big)}
\int_{0}^{L}\!\log(1-u)\,f_{\mathrm{LN}}(u;\mu,\tau^2)\,du.
\] &
Compute \(\mathbb E[\log(1-u)]\) (1D quadrature) and test \(>\log(1-u_1)\). \\[6pt]

\textbf{Truncated gamma} \(\ \mathrm{Ga}_{(0,L]}(\alpha,\beta)\) &
\(\alpha,\beta>0\) feasible. Truncated moments:
$u_2=\frac{1}{\beta}\frac{\gamma(\alpha+1,\beta L)}{\gamma(\alpha,\beta L)},$
$\sigma^2=\frac{1}{\beta^2}\frac{\gamma(\alpha+2,\beta L)}{\gamma(\alpha,\beta L)}-u_2^{\,2},$
$\gamma(a,z)=\int_0^z t^{a-1}e^{-t}\,dt.$ &
\[
\frac{1}{F_G(L;\alpha,\beta)}
\int_{0}^{L}\!\log(1-u)\,
\frac{\beta^\alpha}{\Gamma(\alpha)}u^{\alpha-1}e^{-\beta u}\,du,
\]
\(F_G(L;\alpha,\beta)=\gamma(\alpha,\beta L)/\Gamma(\alpha)\). &
Map \((u_2,\sigma^2)\!\mapsto\!(\alpha,\beta)\), evaluate \(\mathbb E[\log(1-u)]\), and test \(>\log(1-u_1)\). \\
\bottomrule
\end{tabular}%
} % end resizebox

\caption{Summary of \(\mathbb E[\log(1-u_{2,t})]\) and the invasion test \(\mathbb E[\log(1-u_{2,t})]>\log(1-u_1)\) at \(R=0\) for four distribution families on \([0,L]\) (in simulations \(L=1\)). Means and variances refer to the distribution on \([0,L]\). Closed forms are available for some special cases (e.g., Beta with \(L{=}1\)); otherwise a single one–dimensional quadrature suffices.}
\label{tab:lyap_G_summary_landscape}
\end{table}
\end{landscape}


\subsection{\textbf{Sensitivity of $\mathbb{E}[\log(1 - u_{2,t})]$ to Distributional Shape}} \label{app: compare_derivations}
\noindent Fix the variance $\sigma^2\approx0.00159$, support $(0,1)$, and resident mutation rate $u_1\approx0.0954$. For each distribution family, let $u_2^*\in(0,1)$ denote the \emph{largest} mean such that, under complete linkage ($R=0$),
\[
\gamma \;=\; -\log(1-u_1)+\mathbb E\!\big[\log(1-u_{2,t})\big]\;=\;0,
\]
so that $u_2\le u_2^*$ implies invasion at $R=0$. The function $\log(1-u_{2,t})$ is strictly decreasing and strictly concave on $(0,1)$, with derivatives $(1-u_{2,t})^{-1}$, $(1-u_{2,t})^{-2}$, and $(1-u_{2,t})^{-3}$ increasing as $u_{2,t}\!\uparrow1$. Consequently, $\mathbb E[\log(1-u_{2,t})]$ is especially sensitive to probability mass near the upper boundary. To quantify how probability mass relative to $u_1$ influences $\mathbb E[\log(1-u_{2,t})]$, define
\[
\mathsf C(u_1):=\Pr(u_{2,t}\le u_1),\qquad
\mathcal T(u_1):=\mathbb E\!\big[-\log(1-u_{2,t})\,\mathbf 1\{u_{2,t}>u_1\}\big].
\]
\noindent Here, $\mathbf 1\{\cdot\}$ denotes the indicator function, equal to $1$ when the condition inside the bracket is true and $0$ otherwise. The quantity $\mathcal T(u_1)$ measures the expected contribution to $-\log(1-u_{2,t})$ from realizations where $u_{2,t}>u_1$, thus capturing the influence of the right tail of the mutation–rate distribution on the overall mean. Larger $\mathsf C(u_1)$ (greater probability mass at or below $u_1$) increases $\mathbb E[\log(1-u_{2,t})]$, whereas larger $\mathcal T(u_1)$ (heavier right tail above $u_1$) decreases it. For the fixed $\sigma^2$ and $u_1$ given above, Table~\ref{tab:u2star_comparison} reports $u_2^*$ together with $\mathsf C(u_1)$ and $\mathcal T(u_1)$ for the four distribution families.
\begin{table}
\centering
\renewcommand{\arraystretch}{1.12}
\setlength{\tabcolsep}{10pt}
\caption{Largest mean $u_2^*$ (at fixed $\sigma^2$) consistent with $\gamma=0$, together with the $u_1$ cdf and upper–tail penalty.}
\begin{tabular}{lccc}
\toprule
\textbf{Distribution on $[0,1]$} & $\boldsymbol{u_2^*}$ & $\boldsymbol{\mathsf C(u_1)}$ & $\boldsymbol{\mathcal T(u_1)}$\\
\midrule
Uniform                & 0.09427879 & 0.507983 & 0.068280 \\
Beta                   & 0.09448161 & 0.559617 & 0.061754 \\
Truncated log-normal   & 0.09432226 & 0.591052 & 0.057932 \\
Truncated gamma        & 0.09427113 & 0.567150 & 0.060784 \\
\bottomrule
\end{tabular}
\label{tab:u2star_comparison}
\end{table}
\noindent The $u^*_2$ ordering in Table~\ref{tab:u2star_comparison} (beta $>$ truncated log–normal $>$ uniform $>$ truncated gamma) follows from the combined effect of $\mathsf C(u_1)$ and $\mathcal T(u_1)$ together with where mass lies below $u_1$. The beta case has the largest $u_2^*$ by pairing relatively high $\mathsf C$ with moderate $\mathcal T$. The truncated log–normal has the highest $\mathsf C$ and the lowest $\mathcal T$ but yields a slightly smaller $u_2^*$, reflecting diminishing returns from placing additional probability far below $u_1$. The uniform allocates more mass above $u_1$, raising $\mathcal T$ and lowering $u_2^*$. The truncated gamma places comparatively more mass in the upper mid-range than the log–normal, increasing $\mathcal T$ enough to produce the smallest $u_2^*$. Although the numerical differences are small (on the order of $2\times 10^{-4}$), they are systematic: for fixed variance, shifting probability from the right tail toward (and just below) $u_1$ increases $\mathbb E[\log(1-u_{2,t})]$ and thus permits a larger $u_2^*$, while shifting probability toward the boundary $u\approx 1$ has the opposite effect.

\subsection{\textbf{Accuracy of Lyapunov Approximations}}
We compare the Lyapunov invasion criterion with stochastic recursions ($5{,}000$ generations) for four mutation–rate distributions on \([0,L]\) (uniform, beta, truncated log–normal, truncated gamma). With resident \(u_1\approx 0.048796\) and fixed variance \(\sigma^2\), the \(R{=}0\) invasion boundary is the mean \(u_2^\ast\) solving
\[
-\log(1-u_1)+\mathbb E\!\big[\log(1-u_{2,t})\big]=0,
\]
where \(u_{2,t}\) is drawn from the chosen distribution fitted to \(\mathbb E[u_{2,t}]=u_2\) and \(\operatorname{Var}(u_{2,t})=\sigma^2\) (closed form fitting for uniform and beta; moment inversion for truncated distributions). Because \(\log(1-x)\) is strictly decreasing on \([0,1)\), the left-hand side is strictly decreasing in \(u_2\), so any root \(u_2^\ast\) is unique. Numerically, we restrict \(u_2\) to the feasible mean range of each distribution and solve by Brent’s method (tolerance \(10^{-10}\)). Expectations are evaluated exactly for the uniform case and, when available, in closed form for beta on \([0,1]\); otherwise we use one–dimensional adaptive quadrature with an endpoint split near \(L=1\) to handle the logarithmic singularity.

\begin{figure}
\centering
\includegraphics[width=\textwidth]{figures/f4_lyapunov_approx.png}
\caption{\textbf{Fixation outcomes versus Lyapunov thresholds.} Each panel: one mutation–rate distribution. Settings: $5{,}000$ generations, $R=0$, $s=0.2$, $u_1\approx0.048796$. Colors: resident fixation (blue), modifier fixation (orange), no fixation within time range (gray). Red curve: Lyapunov prediction $\gamma(u^*_2)=0$. The predicted boundaries (red) closely tracks the observed transition for all families.}
\label{fig:lyapunov-critical-thresholds}
\end{figure}

\noindent Figure~\ref{fig:lyapunov-critical-thresholds} shows close agreement between simulated fixation outcomes and the Lyapunov threshold. The prediction is slightly conservative: $u_2$ values below the curve lead to reliable invasion. Differences across distributions arise from how fixed variance reallocates probability mass relative to $u_1$ and toward the upper boundary $1$, where $\log(1-u_{2,t})$ is highly sensitive. When most probability lies below $u_1$ and away from $1$ (e.g., truncated log–normal with a thin upper tail, or uniform once its interval includes $u_1$), $\mathbb E[\log(1-u_{2,t})]$ changes little with $\sigma^2$, yielding an almost horizontal boundary in $(u_2,\sigma^2)$.  

\noindent In contrast, distributions that shift mass into the right tail as variance increases (beta, truncated gamma) produce curved Lyapunov boundaries (red): larger $\sigma^2$ raises the upper–tail contribution, lowers $\mathbb E[\log(1-u_{2,t})]$, and reduces $u_2^\ast$. At $\sigma^2\approx8\times10^{-4}$, for example, the beta distribution gives $u_2^\ast\approx0.0484<u_1$, consistent with a modest right tail. The uniform is infeasible at this variance (its interval cannot match the target mean and contain $u_1$). The truncated log–normal and truncated gamma yield $u_2^\ast$ close to $u_1$ ($|u_2^\ast-u_1|\lesssim10^{-3}$), with the gamma slightly lower due to heavier midrange weight. In all cases, shifting probability from below $u_1$ toward $1$ decreases $\mathbb E[\log(1-u_{2,t})]$ and moves the boundary downward, as predicted by the Lyapunov condition.

\section{Recombination effect on Invasion Condition Across Models}
\label{appendix:comparison-models}
\noindent Table~\ref{tab:comparison-grouped} summarizes invasion conditions for a rare modifier $M_2$ under deterministic and stochastic transmission, distinguishing $R=0$ from $0<R\le\tfrac12$. It indicates when recombination can alter the \emph{direction} of selection on $M_2$—that is, whether varying $R$ can switch the outcome between invasion and non–invasion.

\noindent \textit{Deterministic transmission ($u_{2,t}\equiv u_2$).} For both ploidies, the invasion threshold is independent of $R$: $M_2$ increases when rare if $u_2<u_1$ (the \emph{Reduction Principle}; Eq.~\eqref{eq:deterministic_reduction_principle}). For $0<R\le\tfrac12$, recombination moves the dominant eigenvalue toward $1$ in magnitude but cannot change its sign. Thus, under deterministic transmission, recombination affects the \emph{rate} but not the \emph{direction} of invasion.

\noindent \textit{Stochastic transmission ($\mathrm{Var}(u_{2,t})>0$).} With complete linkage ($R=0$), both haploid and diploid models satisfy the geometric–mean criterion
\[
\gamma(0)>0 \;\Longleftrightarrow\; \mathbb{E}[\log(1-u_{2,t})]>\log(1-u_1).
\]
\noindent Because $\log(1-x)$ is strictly concave on $[0,1)$, Jensen’s inequality implies
\[
\mathbb{E}[\log(1-u_{2,t})]\;\le\;\log\!\big(1-\mathbb{E}[u_{2,t}]\big)=\log(1-u_2),
\]
with strict inequality whenever $\mathrm{Var}(u_{2,t})>0$ and $\mathbb{P}(u_{2,t}\in(0,1))>0$. Temporal variability therefore lowers the effective (geometric–mean) growth rate and can prevent invasion even when $u_2<u_1$. Hence, the deterministic \emph{Reduction Principle} may fail under stochastic transmission.

\noindent For $R>0$, products of noncommuting random matrices preclude a closed–form expression for $\gamma(R)$. Recombination has no direct fitness effect but affects associations between the modifier and the selected background that are created each generation by selection and mutation. Depending on how $R$ modulates these associations across the realized distribution of $u_{2,t}$, $\gamma(R)$ can increase or decrease with $R$, allowing \emph{reversals in invasion direction} under a fixed parameter set—an effect absent when $\mathrm{Var}(u_{2,t})=0$. 

\begin{table}[t]
\centering
\footnotesize
\setlength{\tabcolsep}{4pt}
\renewcommand{\arraystretch}{1.15}
\begin{tabularx}{\linewidth}{lYYYY}
\toprule
& \multicolumn{2}{c}{\textbf{Haploid}} & \multicolumn{2}{c}{\textbf{Diploid (additive)}}\\
\cmidrule(lr){2-3}\cmidrule(lr){4-5}
& \textbf{Deterministic} & \textbf{Stochastic} & \textbf{Deterministic} & \textbf{Stochastic}\\
\midrule
\textbf{$R=0$}
&
$\lambda_{+}>1 \iff u_2<u_1$ (\emph{Reduction Principle}; \eqref{eq:deterministic_reduction_principle}). &
$\gamma(0)>0 \iff \mathbb E[\log(1-u_{2,t})]>\log(1-u_1)$ (\eqref{eq:stochastic_R0_inv_condition}); variability strictly lowers $\mathbb E[\log(1-u_{2,t})]$ at constant mean, so invasion may fail even if $u_2<u_1$. &
$\lambda_{+}>1 \iff u_2<u_1$ (\emph{Reduction Principle}; \eqref{eq:deterministic_reduction_principle}). &
$\gamma(0)>0 \iff \mathbb E[\log(1-u_{2,t})]>\log(1-u_1)$ (\eqref{eq:stochastic_R0_inv_condition}); same conclusion. \\

\addlinespace[2pt]
\textbf{$0<R\le \tfrac12$}
&
Sign independent of $R$; $R$ pulls $\lambda_{+}$ toward $1$ (magnitude only). &
$\gamma(R)$ exists (under standard moment/positivity conditions) and can change sign with $R$; \emph{reversals observed in both directions}. &
Sign independent of $R$; $R$ pulls $\lambda_{+}$ toward $1$ (magnitude only). &
Same qualitative behavior as haploids; reversals possible, with the $R$–response typically smaller for comparable parameters. \\
\bottomrule
\end{tabularx}
\caption{\textbf{Invasion conditions and recombination across deterministic and stochastic transmission.} 
Deterministic transmission: the \emph{Reduction Principle} holds for all $R$; recombination alters the magnitude but not the sign of initial growth. 
Stochastic transmission: at $R{=}0$, invasion depends on the geometric mean $\mathbb E[\log(1-u_{2,t})]$; temporal variability can overturn the deterministic prediction even without recombination. For $R>0$, $\gamma(R)$ may change sign as $R$ varies, producing reversals that do not occur when $\mathrm{Var}(u_{2,t}){=}0$.}
\label{tab:comparison-grouped}
\end{table}


\section{Empirical Analysis - \textit{Arabidopsis thaliana} } \label{appendix:empirical}
We examine transmission–related determinants of mutation for genes annotated as essential or lethal in \textit{A.~thaliana}. These loci are expected to experience consistently strong purifying selection, reducing confounding from fluctuating selective regimes. Within this constrained functional class, residual variation in mutation rates should more plausibly reflect transmission mechanisms---chromatin configuration, local DNA accessibility, and DNA–repair efficacy---rather than changing selection pressures \citep{monroe2022}.

\paragraph{Predictors of Mutation Rate.}
Following Monroe et al.\ \cite{monroe2022}, the response for gene $i$ is the per–base mutation rate
\[
\text{mut\_rate}_{i}=\frac{\text{de novo mutations in gene }i}{\text{CDS length}_{i}},
\]
modeled as a function of sequence composition (GC content), cytosine methylation (CG/CHG/CHH), chromatin accessibility (ATAC–seq), and histone marks (H3K4me1/2/3, H3K27ac, H3K14ac, H3K27me1, H3K36ac/3, H3K56ac, H3K9ac/me1/me2, H3K23ac). Variables reported on a $0$–$100$ scale were rescaled to $[0,1]$; all predictors were standardized to $z$–scores. Thus, in linear models, coefficients represent the change in \emph{expected} per–base mutation rate associated with a $+1$\,SD change in a predictor, conditional on others at their means. Gene–level observations are treated as independent; heteroskedasticity and mean–variance coupling with gene length are addressed below.

\noindent We constructed a stable predictor set in three steps: \emph{(i) Multicollinearity control:} we iteratively removed predictors with variance–inflation factor (VIF) $>10$, a conventional threshold that guards against unstable coefficient estimation. \emph{(ii) Multiple–testing control across models:} for each predictor, we computed Benjamini–Hochberg FDR $q$–values within each model and averaged these $q$–values across models to obtain a cross–model significance ranking. \emph{(iii) Model diversity:} we fit three multivariable specifications to the same predictors:
\begin{align*}
\text{(a) OLS with HC3:}\quad 
&\text{mut\_rate}_i=\beta_0+\mathbf x_i^\top\boldsymbol\beta+\varepsilon_i,\ \ \text{HC3 standard errors,} \\
\text{(b) WLS (length–weighted):}\quad 
&\text{mut\_rate}_i=\beta_0+\mathbf x_i^\top\boldsymbol\beta+\varepsilon_i,\ \ \text{weights } w_i=\text{CDS length}_i, \\
\text{(c) Poisson GLM (counts with offset):}\quad 
& C_i\sim\text{Poisson}(\mu_i),\ \ \log \mu_i=\log(\text{CDS length}_i)+\beta_0+\mathbf x_i^\top\boldsymbol\beta,
\end{align*}
where $C_i$ denotes the raw mutation count. In (c), $\exp(\beta_j)$ is a rate ratio per $+1$\,SD in predictor $j$, holding others fixed; in (a,b), $\beta_j$ is a marginal slope for the per–base rate. HC3 ensures valid Wald tests under heteroskedasticity (given independence), while WLS stabilizes variance when the error variance scales with gene length.

\noindent To focus inference on effects that are stable across specifications and adjusted for multiplicity, we defined the \emph{predictor set} by retaining the five predictors with the smallest mean BH–$q$ across the three models (a pre–specified cap to promote parsimony and curb residual collinearity). The resulting set was \{H3K4me1, H3K4me3, H3K36ac, GC\_content\_pct, CG\_pct\}. We then refit the OLS model on this reduced set (after a second VIF screen to verify $\text{VIF}\le10$ for all retained predictors) and summarized partial effects by varying one predictor at a time while fixing others at their standardized means (Fig.~\ref{fig:regression_lines}). This sequence—screen for collinearity, control FDR across complementary models, then refit on the selected subset—yields coefficient estimates and uncertainty that are interpretable, comparably scaled, and robust to reasonable changes in distributional assumptions.

\noindent In the joint model, three histone marks—H3K4me1, H3K4me3, and H3K36ac—showed strong negative associations with per–base mutation rate:

\[\widehat\beta_{\text{H3K4me1}}=-1.98\times 10^{-6}\ (q_{\text{BH}}=9.19\times10^{-9}),\]
\[\widehat\beta_{\text{H3K4me3}}=-9.99\times 10^{-7}\ (q_{\text{BH}}=1.08\times10^{-3}),\]
\[\widehat\beta_{\text{H3K36ac}}=-5.74\times 10^{-7}\ (q_{\text{BH}}=1.08\times10^{-3}).\]
\noindent GC content showed a small, non–significant positive effect ($\widehat\beta=2.24\times10^{-7}$, $q_{\text{BH}}=0.278$), while CG methylation had a small, non–significant negative effect ($\widehat\beta=-1.24\times10^{-7}$, $q_{\text{BH}}=0.409$). The signs and relative magnitudes of all coefficients were consistent across the WLS and Poisson–GLM models used for sensitivity analysis. The negative coefficients for H3K4me1, H3K4me3, and H3K36ac indicate that regions marked by these histone modifications are associated with lower mutation rates, consistent with enhanced DNA repair activity in transcriptionally active chromatin. In contrast, the weak methylation effect likely reflects reduction of its apparent influence once correlated chromatin features are included \citep{monroe2022}.

\noindent Two caveats apply. First, restricting analysis to essential or lethal genes reduces variation in several predictors (e.g., GC content, chromatin accessibility), lowering statistical power and thereby reducing detectable marginal effects. Second, residual correlations among histone marks remain even after VIF pruning. Coefficients should therefore be interpreted as \emph{partial associations conditional on the other included predictors}, not as independent causal effects.
\begin{figure}[h!]
    \centering
    \includegraphics[width=0.85\textwidth]{figures/empirical/af1_reg_lines_mutations.png}
    \caption{\textbf{Rigor set—partial effects with 95\% CI.} 
    OLS (HC3) partial–effect curves for the five rigor–set predictors. Each curve varies one standardized predictor across its 1st–99th percentile while holding the others at their means (0). Shaded bands are 95\% confidence intervals; coefficients and FDR $q$–values for the same model are reported in the text.}
    \label{fig:regression_lines}
\end{figure}

\paragraph{Predictors and Mutation Moments.}
To examine distributional properties in addition to the mean, we extended the regression to higher moments using the same standardized predictor set (\emph{i.e.}, each predictor $X_j$ was centered and scaled as $X_j^{\ast}=(X_j-\bar X_j)/\mathrm{SD}(X_j)$ to have mean~0 and unit variance: H3K4me1, H3K4me3, H3K36ac, GC content, CG methylation). After fitting the mean model, we modeled dispersion and shape from the residuals: (i) a log–variance regression on squared residuals (reporting percent change in residual variance per $+1$\,SD in a predictor), (ii) a skewness regression on cubed standardized residuals, and (iii) a kurtosis regression on the fourth standardized residual moment centered at~3. All specifications used heteroskedasticity–robust covariance and Benjamini–Hochberg adjustment.

\noindent The variance models indicate strong stabilizing associations for active chromatin marks:

$$\text{H3K4me1: } -84.9\%\ (q_{\text{BH}}\ll 10^{-10}),$$
$$\text{H3K4me3: } -58.9\%\ (q_{\text{BH}}\ll 10^{-10}),$$
$$\text{H3K36ac: } -36.2\%\ (q_{\text{BH}}\ll 10^{-10}),$$

\noindent consistent with their negative mean effects (Fig.~\ref{fig:regression_lines}) and with reduced variance across genes (Fig.~\ref{fig:moment_heatmap}). In contrast, GC content shows a positive variance association ($+38.7\%$, $q_{\text{BH}}<10^{-11}$), indicating greater mutation–rate variability in GC–rich regions. CG methylation exhibits a modest stabilizing association ($-18.2\%$, $q_{\text{BH}}\approx 5\times 10^{-6}$).

\noindent For skewness and kurtosis, estimates are imprecise and not significant after FDR correction (all $q_{\text{BH}}>0.8$). Point estimates suggest that GC content and CG methylation, while associated with higher mean and (for GC) higher variance, are simultaneously associated with lower skewness and kurtosis, whereas H3K4me1/3/36ac display the opposite qualitative pattern. Given the uncertainty of higher–moment regressions, these patterns should be interpreted as descriptive. They are nevertheless consistent with a potential trade–off in which predictors that increase overall dispersion (mean/variance) may correspondingly reduce tail weight (skewness/kurtosis), and vice versa.

\noindent Overall, H3K4me1, H3K4me3, and H3K36ac show a coherent stabilizing profile—lower mean and substantially lower variance—within these essential/lethal genes, while GC content increases variability despite a weak mean effect. This moment–based analysis complements the mean regression by indicating that epigenomic context is associated not only with the \emph{level} of mutation but also with its \emph{dispersion} and, tentatively, its tail shape across genes.

\begin{figure}[h!]
    \centering
    \includegraphics[width=0.7\textwidth]{figures/empirical/af2_predictors_moments.png}
    \caption{\textbf{Predictors $\times$ moments of the per–bp mutation rate (rigor set, essential/lethal genes).} 
    Cells report regression coefficients for the mean (per–bp mutation rate), variance (percent change in residual variance per +1~SD), skewness, and kurtosis. Stars denote FDR significance. The color scale indicates column–wise standardized effect size (diverging, centered at~0).}
    \label{fig:moment_heatmap}
\end{figure}

\paragraph{Trends Across Tajima's D Deciles.}
We examined whether a proxy for purifying selection covaries with (i) the epigenomic predictor set and (ii) the first four moments of the per–base mutation–rate distribution. For each essential/lethal gene $i$, let $D_i$ denote Tajima’s $D$. With $\bar D$ and $\mathrm{sd}(D)$ the sample mean and standard deviation over the analyzed genes, define a standardized selection score
\[
S_i=-\frac{D_i-\bar D}{\mathrm{sd}(D)},
\]
so larger $S_i$ indicates more negative $D_i$ (stronger putative purifying selection). All predictors were $z$–scored, and inference used heteroskedasticity–robust covariance with Benjamini–Hochberg FDR control.

\noindent To assess associations with the epigenomic predictors, we regressed each predictor $X_j$ (H3K4me1, H3K4me3, H3K36ac, GC\_content\_pct, CG\_pct) on $S$ while controlling for the other four predictors. The coefficient $\gamma_j$ measures the change in $X_j$ (SD units) per $+1$\,SD increase in $S$. Only GC\_content\_pct increased significantly with $S$ ($\hat\gamma=0.048$, 95\% CI $[0.015,0.081]$, $q=0.020$). Partial slopes for the remaining predictors were small and non–significant: H3K4me1 ($\hat\gamma=0.022$, $q=0.142$), CG\_pct ($0.017$, $q=0.432$), H3K4me3 ($0.010$, $q=0.432$), H3K36ac ($-0.009$, $q=0.494$). Raw Spearman correlations between $S$ and each predictor were near zero (e.g., GC\_content\_pct $\rho\approx0.04$), indicating that the GC trend arises only after conditioning on the other marks.

\noindent We next asked whether $S$ predicts the mean and higher moments of the per–base mutation rate after adjusting for the predictor set. Specifically, we fit (i) a mean model for the mutation rate, (ii) a log–variance regression on squared residuals, (iii) a regression for skewness based on cubed standardized residuals, and (iv) a regression for kurtosis based on the fourth standardized residual moment centered at 3. Estimated effects of $S$ per $+1$\,SD were uniformly negative—mean $\hat\beta_S=-5.56\times10^{-8}$ ($q=0.652$), variance $\Delta=-5.2\%$ ($q=0.185$), skewness $\hat\eta_S=-13.7$ ($q=0.266$), kurtosis $\hat\kappa_S=-276$ ($q=0.201$)—but none met the FDR threshold.

\noindent The GC content result indicates that, within essential/lethal genes, stronger purifying selection is associated with modestly higher GC even after conditioning on chromatin marks. Possible mechanisms include GC–biased gene conversion or codon–usage constraints; the analysis is correlative and does not distinguish among them. For mutation moments, the negative point estimates are directionally consistent with lower means and reduced dispersion under stronger selection, but wide robust intervals indicate limited precision.

\noindent Tajima’s $D$ aggregates effects of demography and linked selection and is noisy for low–diversity, constraint–rich loci. Together with the essential/lethal restriction—which compresses both epigenomic and diversity variation—these factors limit power. Accordingly, all results should be interpreted as partial associations rather than causal effects (Fig.~\ref{fig:tajimas_trends}).

\begin{figure}
  \centering
  \includegraphics[width=\textwidth]{figures/empirical/af3_tajimas_trends.png}
   \caption{\textbf{Selection intensity ($S=-z($Tajima’s $D)$) versus predictors and mutation moments in essential/lethal genes.}
  \emph{Left (A):} Adjusted trends for each rigor predictor from $X_j\!\sim\!S+X_{-j}$ with robust 95\% CIs; predictors are standardized ($z$; i.e., expressed in standard deviation units).. 
  \emph{Right (B):} Predicted mean, variance (reported as \% change relative to $S{=}0$ and then standardized for display), skewness, and kurtosis from the moment models with robust 95\% CIs; rigor predictors are held at their means. Larger $S$ denotes stronger purifying selection.}
  \label{fig:tajimas_trends}
\end{figure}

\paragraph{Interpretations.} All analyses are restricted to essential/lethal genes, which reduces confounding from fluctuating selection but also compresses variation in predictors and outcomes, thereby lowering power. Results are therefore interpreted as partial associations, not causal effects.

\noindent In the joint linear model with standardized predictors, variance–inflation–factor (VIF) screening, and heteroskedasticity–robust inference, three histone marks linked to active chromatin show negative associations with the per–base mutation rate: H3K4me1 ($-1.98\times10^{-6}$ per $+1$\,SD; $q=9.2\times10^{-9}$), H3K4me3 ($-9.99\times10^{-7}$; $q\approx1.1\times10^{-3}$), and H3K36ac ($-5.74\times10^{-7}$; $q\approx1.1\times10^{-3}$). GC\% and CG\% exhibit small, non–significant mean effects ($q\approx0.28$ and $q\approx0.41$). Here, $q$ denotes the Benjamini–Hochberg false–discovery–rate–adjusted $p$–value used for multiple–testing control. Thus, conditional on other marks, active chromatin is associated with lower average mutation rates, whereas base composition and CG methylation are not.

\noindent Extending the framework to residual dispersion reveals strong associations with variance. Per $+1$\,SD in the predictor, residual variance decreases by $84.9\%$ for H3K4me1, $58.9\%$ for H3K4me3, and $36.2\%$ for H3K36ac (all $q\ll10^{-10}$), whereas GC\% increases variance by $38.7\%$ ($q\approx1.3\times10^{-12}$) and CG\% reduces it by $18.2\%$ ($q\approx4.9\times10^{-6}$). These results indicate that epigenomic context is associated not only with the mean level of mutation but also with the degree of \emph{between–gene variability} (i.e., heterogeneity in mutation rates across genes). Regressions for skewness and kurtosis of variance–standardized residuals yield imprecise estimates (all $q\approx0.82$–$0.90$). Point estimates suggest opposite directions for active marks versus CG methylation, but the lack of FDR significance precludes inference about asymmetry or tail thickness.

\noindent Using $S=-z(D)$ as a standardized proxy for stronger purifying selection (more negative Tajima’s $D$), we regressed each predictor on $S$ while adjusting for the remainder. Only GC\% increases with $S$ ($\hat\gamma=0.048$ SD per $+1$\,SD in $S$; 95\% CI $[0.015,0.081]$; $q=0.020$), where $\hat\gamma$ denotes the standardized partial regression slope. H3K4me1, H3K4me3, H3K36ac, and CG\% show small, non–significant partial slopes ($q\in[0.14,0.49]$). When the mean and higher moments of the mutation rate are modeled as functions of $S$ (conditional on the chromatin predictors), estimated slopes are uniformly negative—mean $-5.6\times10^{-8}$ ($q=0.65$), variance $-5.2\%$ ($q=0.19$), skewness $-13.7$ ($q=0.27$), kurtosis $-276$ ($q=0.20$)—but none meet the FDR threshold.

\noindent Two implications follow. First, H3K4me1, H3K4me3, and H3K36ac are jointly associated with lower mutation means and substantially lower variance, whereas GC\% increases variance despite a weak mean effect. Second, although estimates for each predictor versus $S$ are imprecise, their consistently negative signs are compatible with stronger long–term purifying selection being associated with lower average mutation rates and reduced dispersion. Given limited power in the essential/lethal subset and the composite nature of Tajima’s $D$, these patterns should be viewed as suggestive and subject to confirmation in broader gene sets.

\newpage
\section*{Code and supplementary material}

All code and supplementary material used for the analyses in this paper are available at
\url{https://github.com/ElisaHeinrich/Evo_Stochastic_Transmission_Mut_Modifiers}.


\bibliography{report} 
\bibliographystyle{abbrv}
\end{document}
