## Supplementary material for "Evolution under Stochastic Transmission: Mutation-Rate Modifiers": Full Project Package: report.bib

%% report.bib - contains BiBtex input for bibliography

@phdthesis{altenberg1984,
  author = {Altenberg, Lee},
  title = {A Generalization of Theory on the Evolution of Modifier Genes},
  school = {Stanford University},
  year = {1984},
  type = {PhD dissertation}
}

@article{altenberg2017unified,
  title={Unified reduction principle for the evolution of mutation, migration, and recombination},
  author={Altenberg, Lee and Liberman, Uri and Feldman, Marcus W},
  journal={Proceedings of the National Academy of Sciences},
  volume={114},
  number={12},
  pages={E2392--E2400},
  year={2017},
  publisher={National Acad Sciences}
}

@article{balkau1973selection,
  title={Selection for migration modification},
  author={Balkau, Beverley J and Feldman, Marcus W},
  journal={Genetics},
  volume={74},
  number={1},
  pages={171--174},
  year={1973},
  publisher={Oxford University Press}
}

@article{carja2014evolution,
  title={Evolution in changing environments: Modifiers of mutation, recombination, and migration},
  author={Carja, Oana and Liberman, Uri and Feldman, Marcus W},
  journal={Proceedings of the National Academy of Sciences},
  volume={111},
  number={50},
  pages={17935--17940},
  year={2014},
  publisher={National Acad Sciences}
}

@article{charlesworth1976recombination,
  author       = {Charlesworth, Brian},
  title        = {Recombination modification in a fluctuating environment},
  journal      = {Genetics},
  year         = {1976},
  month        = {May},
  volume       = {83},
  number       = {1},
  pages        = {181--195},
  doi          = {10.1093/genetics/83.1.181},
  pmid         = {1269919},
  pmcid        = {PMC1213499}
}

@article{cookHartl1974,
  author  = {Cook, R. Dennis and Hartl, Daniel L.},
  title   = {Uncorrelated Random Environments and Their Effects on Gene Frequency},
  journal = {Evolution},
  year    = {1974},
  volume  = {28},
  number  = {2},
  pages   = {265--274},
  doi     = {10.1111/j.1558-5646.1974.tb00746.x}
}

@article{feldman1976krakauer,
  title={Genetic modification and modifier polymorphisms.},
  author={Feldman, MW and Krakauer, J},
  year={1976}
}

@article{feldman1972selection,
  title={Selection for linkage modification. I. Random mating populations},
  author={Feldman, Marcus W},
  journal={Theoretical Population Biology},
  volume={3},
  number={3},
  pages={324--346},
  year={1972},
  month={September},
  doi={10.1016/0040-5809(72)90007-x},
  PMID={4667090}
}

@article{feldman1986evolutionary,
  title={An evolutionary reduction principle for genetic modifiers.},
  author={Feldman, Marcus W and Liberman, Uri},
  journal={Proceedings of the National Academy of Sciences},
  volume={83},
  number={13},
  pages={4824--4827},
  year={1986}
}

@article{feldman1996population,
  title={Population genetic perspectives on the evolution of recombination},
  author={Feldman, Marcus W and Otto, Sarah P and Christiansen, Freddy B},
  journal={Annual review of genetics},
  volume={30},
  number={1},
  pages={261--295},
  year={1996},
  publisher={Annual Reviews 4139 El Camino Way, PO Box 10139, Palo Alto, CA 94303-0139, USA}
}

@article{FledelAlon2011,
  author  = {Fledel-Alon, A. and Leffler, E.~M. and Guan, Y. and Stephens, M. and Coop, G. and Przeworski, M.},
  title   = {Variation in Human Recombination Rates and Its Genetic Determinants},
  journal = {PLoS ONE},
  year    = {2011},
  volume  = {6},
  number  = {6},
  pages   = {e20321},
  doi     = {10.1371/journal.pone.0020321}
}

@article{frobenius1908_1909,
  author       = {Frobenius, G.},
  title        = {Über Matrizen aus positiven Elementen},
  journal      = {Sitzungsberichte der Königlich Preussischen Akademie der Wissenschaften zu Berlin},
  year         = {1908--1909},
  pages        = {471--476, 514--518},
}

@article{furstenberg1960,
  author  = {Furstenberg, Harry and Kesten, Harry},
  title   = {Products of Random Matrices},
  journal = {Annals of Mathematical Statistics},
  volume  = {31},
  number  = {2},
  pages   = {457--469},
  year    = {1960},
  month   = jun,
  doi     = {10.1214/aoms/1177705909}
}

@article{gillespie1972,
  author  = {Gillespie, John H.},
  title   = {The Effects of Stochastic Environments on Allele Frequencies in Natural Populations},
  journal = {Theoretical Population Biology},
  year    = {1972},
  volume  = {3},
  number  = {3},
  pages   = {241--248},
  doi     = {10.1016/0040-5809(72)90001-9}
}

@article{gillespie1973,
  author  = {Gillespie, John H.},
  title   = {Natural Selection with Varying Selection Coefficients—A Haploid Model},
  journal = {Genetical Research},
  year    = {1973},
  volume  = {21},
  number  = {2},
  pages   = {115--120},
  doi     = {10.1017/S001667230001329X}
}

@article{karlin1974random,
  title={Random temporal variation in selection intensities: case of large population size.},
  author={Karlin, Samuel and Lieberman, Uri},
  journal = {Theoretical Population Biology},
  year={1974}
}

@article{karlinLiberman1975,
  author  = {Karlin, Samuel and Liberman, Uri},
  title   = {Random Temporal Variation in Selection Intensities: One-Locus Two-Allele Model},
  journal = {Journal of Mathematical Biology},
  year    = {1975},
  volume  = {2},
  number  = {1},
  pages   = {1--17},
  doi     = {10.1007/BF00276012}
}

@article{karlin1974towards,
  title={Towards a theory of the evolution of modifier genes},
  author={Karlin, Samuel and McGregor, James},
  journal={Theoretical population biology},
  volume={5},
  number={1},
  pages={59--103},
  year={1974},
  publisher={Elsevier}
}

@article{lewontin1969population,
  title={On population growth in a randomly varying environment},
  author={Lewontin, Richard C and Cohen, Daniel},
  journal={Proceedings of the National Academy of sciences},
  volume={62},
  number={4},
  pages={1056--1060},
  year={1969}
}

@article{liberman1986modifiers,
  title={Modifiers of mutation rate: a general reduction principle},
  author={Liberman, Uri and Feldman, Marcus W},
  journal={Theoretical population biology},
  volume={30},
  number={1},
  pages={125--142},
  year={1986},
  publisher={Elsevier}
}

@article{liberman2011evolution,
  title={On the evolution of mutation in changing environments: recombination and phenotypic switching},
  author={Liberman, Uri and Van Cleve, Jeremy and Feldman, Marcus W},
  journal={Genetics},
  volume={187},
  number={3},
  pages={837--851},
  year={2011},
  publisher={Oxford University Press}
}

@article{monroe2022,
  author       = {Monroe, J. G. and Srikant, T. and Carbonell-Bejerano, P. and others},
  title        = {Mutation bias reflects natural selection in \textit{Arabidopsis thaliana}},
  journal      = {Nature},
  year         = {2022},
  volume       = {602},
  pages        = {101--105},
  doi          = {10.1038/s41586-021-04269-6}
}

@article{rozhok2021silico,
  title={In silico experiments uncover a novel mechanism underlying mutation rate evolution in sexually reproducing populations},
  author={Rozhok, Andrii and Eldredge, Niles and DeGregori, James},
  journal={bioRxiv},
  pages={2021--09},
  year={2021},
  publisher={Cold Spring Harbor Laboratory}
}

@article{teague1976result,
  title={A result on the selection of recombination altering mechanisms.},
  author={Teague, R},
  year={1976}
}

@article{TWOMEY1990320,
	abstract = {Two diallelic loci in an infinite panmictic population of diploid individuals are modelled. The Aa locus is subject to unidirectional mutation and either multiplicative fertility selection or, equivalently, sex-asymmetric viability selection. The Mm locus acts as a selectively neutral modifier of the mutation rate at Aa. The loci recombine at rate R. If the Mm locus is initially monomorphic, and the Aa locus has reached equilibrium, the fate of a new modifier allele is found to depend not just on its relative effect on mutation but also upon the linkage, R. Each initial equilibrium may be characterized by a critical value of the recombination rate, R∗. If 0 < R∗ < 0.5, a sufficiently small ``down'' modifier of the mutation rate will invade the population when R < R∗ whereas a sufficiently small ``up'' modifier will succeed when R > R∗. If R∗ < 0 or R∗ > 0.5, only mutation reduction may occur. Numerical analysis of 56,000 sample equilibria indicates that mutation rates may be increased, but only when the selection regime is such that the Aa locus would remain polymorphic in the absence of mutation.},
	author = {Michael J. Twomey and Marcus W. Feldman},
	doi = {https://doi.org/10.1016/0040-5809(90)90042-T},
	issn = {0040-5809},
	journal = {Theoretical Population Biology},
	number = {2},
	pages = {320-342},
	title = {Mutation modification with multiplicative fertility selection},
	url = {https://www.sciencedirect.com/science/article/pii/004058099090042T},
	volume = {37},
	year = {1990},
	bdsk-url-1 = {https://www.sciencedirect.com/science/article/pii/004058099090042T},
	bdsk-url-2 = {https://doi.org/10.1016/0040-5809(90)90042-T}}

@article{wei2022,
  author       = {Wei, W. and Ho, W.-C. and Behringer, M. G. and others},
  title        = {Rapid evolution of mutation rate and spectrum in response to environmental and population-genetic challenges},
  journal      = {Nature Communications},
  year         = {2022},
  volume       = {13},
  pages        = {4752},
  doi          = {10.1038/s41467-022-32353-6}
}


%%%%%%%%%%%%%%%%%%%%%%%%%%%%%%%%%%%%%%%%%%%%%%%%%%%%%%%%%%%%%%%%%%%%%%%%%%%%%%%%%%%%

@book{loeve1977,
  author    = {Lo{\`e}ve, Michel},
  title     = {Probability Theory I},
  edition   = {4th},
  year      = {1977},
  publisher = {Springer},
  address   = {New York},
  series    = {Graduate Texts in Mathematics},
  volume    = {45},
  isbn      = {978-0387902780}
}
@article{oseledets1968,
  author  = {Oseledets, V. I.},
  title   = {A Multiplicative Ergodic Theorem: Lyapunov Characteristic Numbers for Dynamical Systems},
  journal = {Transactions of the Moscow Mathematical Society},
  volume  = {19},
  year    = {1968},
  pages   = {197--231},
  note    = {Russian original in: \emph{Moscow Mathematical Society}, 19 (1968), 179--210}
}

@article{furstenberg1960products,
  title={Products of random matrices},
  author={Furstenberg, Harry and Kesten, Harry},
  journal={The Annals of Mathematical Statistics},
  volume={31},
  number={2},
  pages={457--469},
  year={1960},
  publisher={JSTOR}
}

@article{bennett1954theory,
  title={On the theory of random mating},
  author={Bennett, JH},
  journal={Annals of Eugenics},
  volume={18},
  number={4},
  pages={311--317},
  year={1954},
  month={March},
  doi={10.1111/j.1469-1809.1952.tb02522.x},
  PMID={13148997}
}

@article{kimura1964diffusion,
  title={Diffusion models in population genetics},
  author={Kimura, Motoo},
  journal={Journal of Applied Probability},
  volume={1},
  number={2},
  pages={177--232},
  year={1964},
  publisher={Cambridge University Press}
} 

@article{bodmer1967linkage,
  title={Linkage and selection: theoretical analysis of the deterministic two locus random mating model},
  author={Bodmer, Walter F and Felsenstein, Joseph},
  journal={Genetics},
  volume={57},
  number={2},
  pages={237--265},
  year={1967},
  month={October},
  doi={10.1093/genetics/57.2.237},
  url={https://doi.org/10.1093/genetics/57.2.237}
}

@article{Karlin_Feldman_1970, title={Convergence to equilibrium of the two locus additive viability model}, volume={7}, DOI={10.2307/3211963}, number={2}, journal={Journal of Applied Probability}, author={Karlin, Samuel and Feldman, Marcus W.}, year={1970}, pages={262–271}} <div></div>

@article{karlin1970linkage,
  title={Linkage and selection: two locus symmetric viability model},
  author={Karlin, Samuel and Feldman, Marcus W},
  journal={Theoretical Population Biology},
  volume={1},
  number={1},
  pages={39--71},
  year={1970},
  month={May},
  doi={10.1016/0040-5809(70)90041-9},
  PMID={5527625}
}

@article{feldman1973selection,
  title={Selection for linkage modification II. A recombination balance for neutral modifiers},
  author={Feldman, Marcus W and Balkau, Beverley},
  journal={Genetics},
  volume={74},
  number={4},
  pages={713--726},
  year={1973},
  publisher={Oxford University Press}
}

@article{feldman1986evolutionary,
  title={An evolutionary reduction principle for genetic modifiers.},
  author={Feldman, Marcus W and Liberman, Uri},
  journal={Proceedings of the National Academy of Sciences},
  volume={83},
  number={13},
  pages={4824--4827},
  year={1986},
  publisher={National Acad Sciences}
}


@article{Swain2002,
  author = {Swain, P. S. and Elowitz, M. B. and Siggia, E. D.},
  title = {Intrinsic and Extrinsic Contributions to Stochasticity in Gene Expression},
  journal = {Proceedings of the National Academy of Sciences},
  volume = {99},
  number = {20},
  pages = {12795--12800},
  year = {2002},
  doi = {10.1073/pnas.162041399},
  url = {https://doi.org/10.1073/pnas.162041399}
}

@article{Barnett2025,
  author = {Barnett, Michael and Meister, Lena and Rainey, Paul B.},
  title = {Experimental Evolution of Evolvability},
  journal = {Science},
  volume = {387},
  number = {6736},
  pages = {eadr2756},
  year = {2025},
  doi = {10.1126/science.adr2756},
  url = {https://www.science.org/doi/10.1126/science.adr2756}
}
