## Supplementary figures and images for "Evolution under Stochastic Transmission: Mutation-Rate Modifiers"

### af1_reg_lines_mutations.png

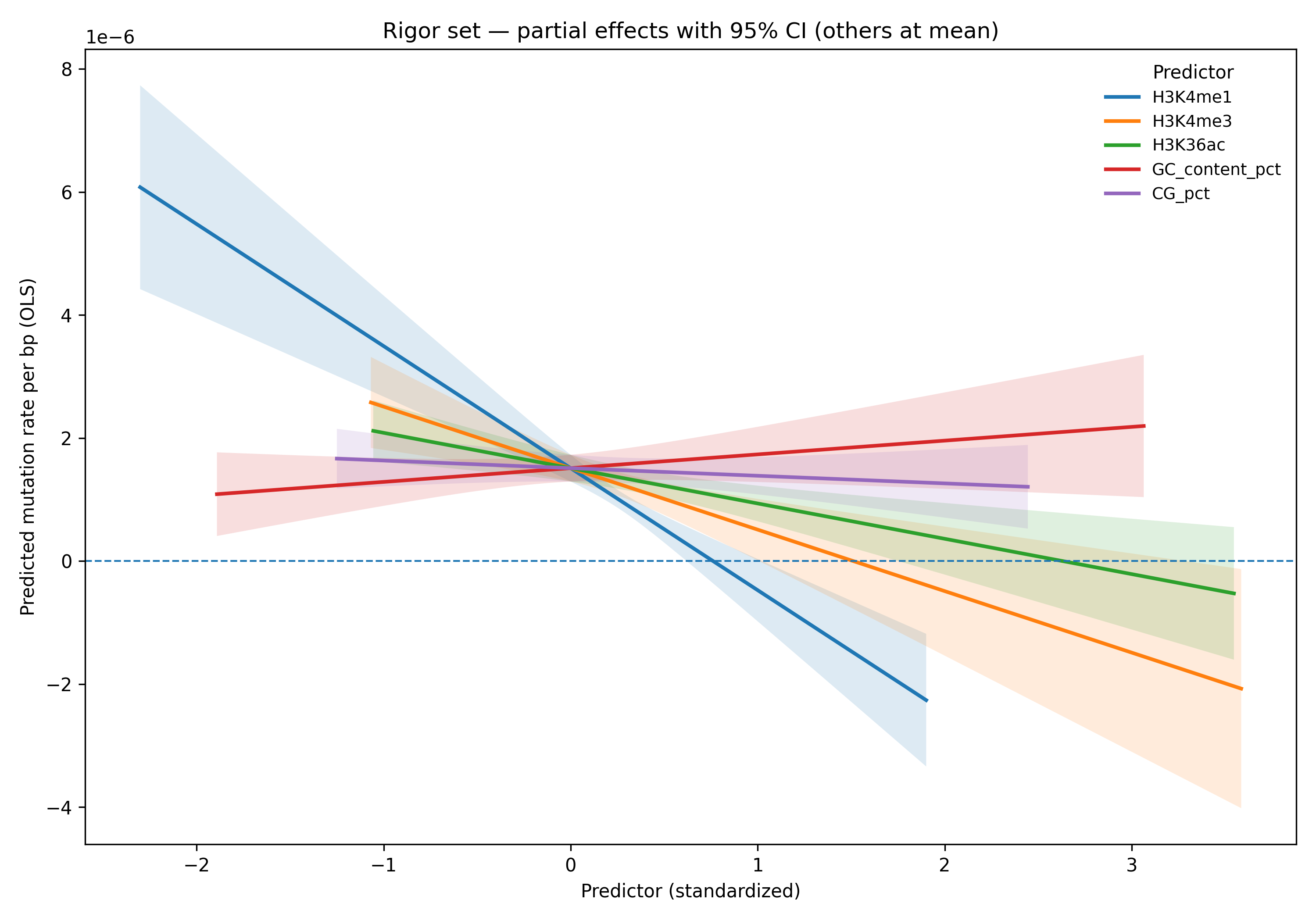

### af2_predictors_moments.png

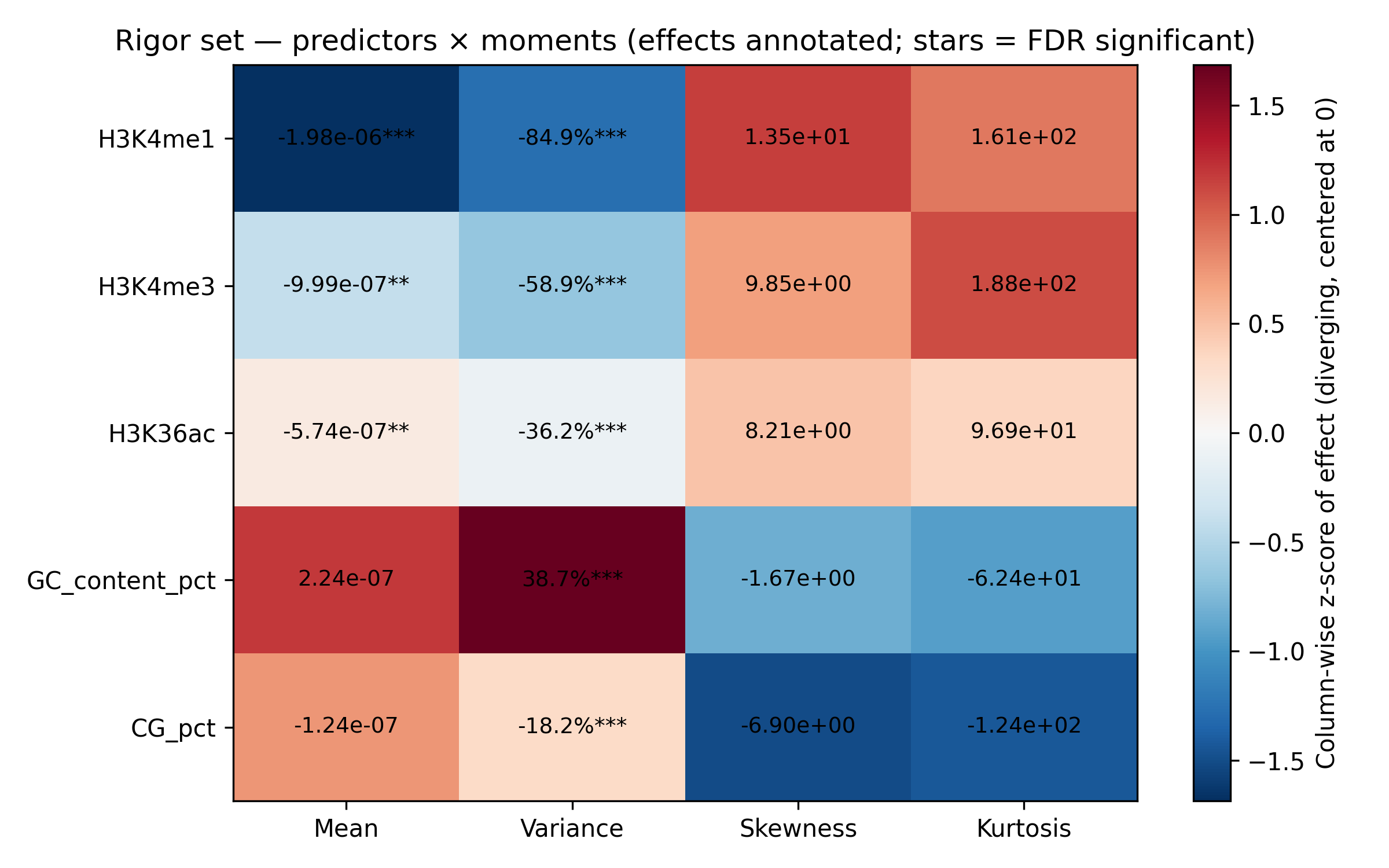

### af3_tajimas_trends.png

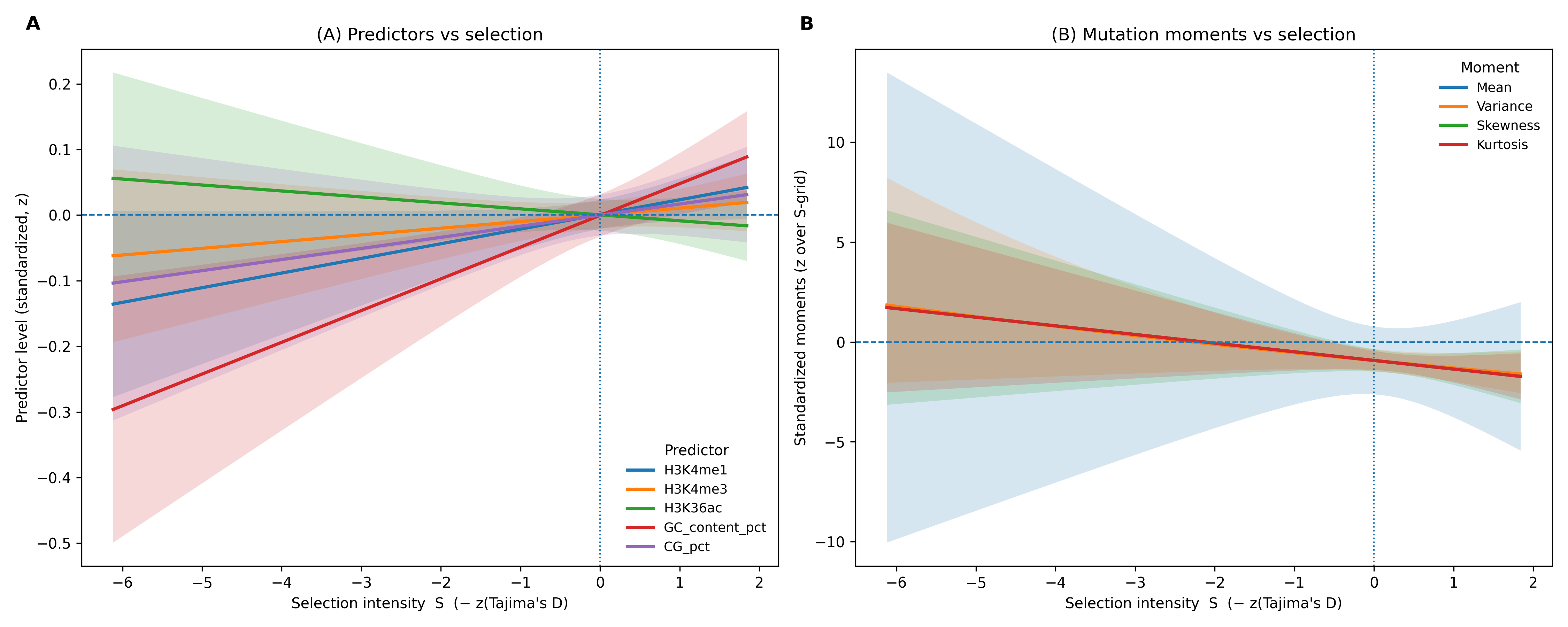

### f1_gamma_lyap_composite.png

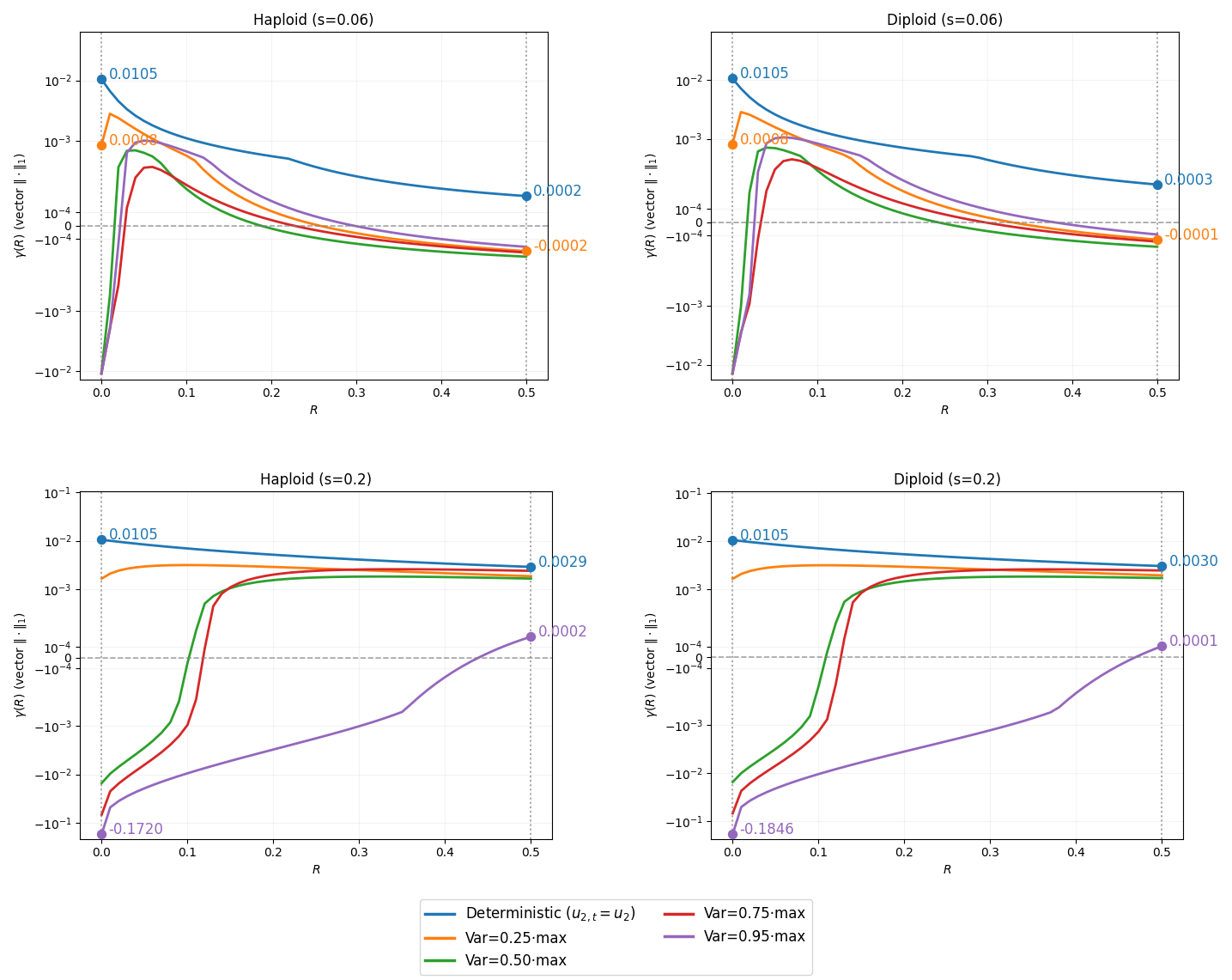

### f2_distributions.png

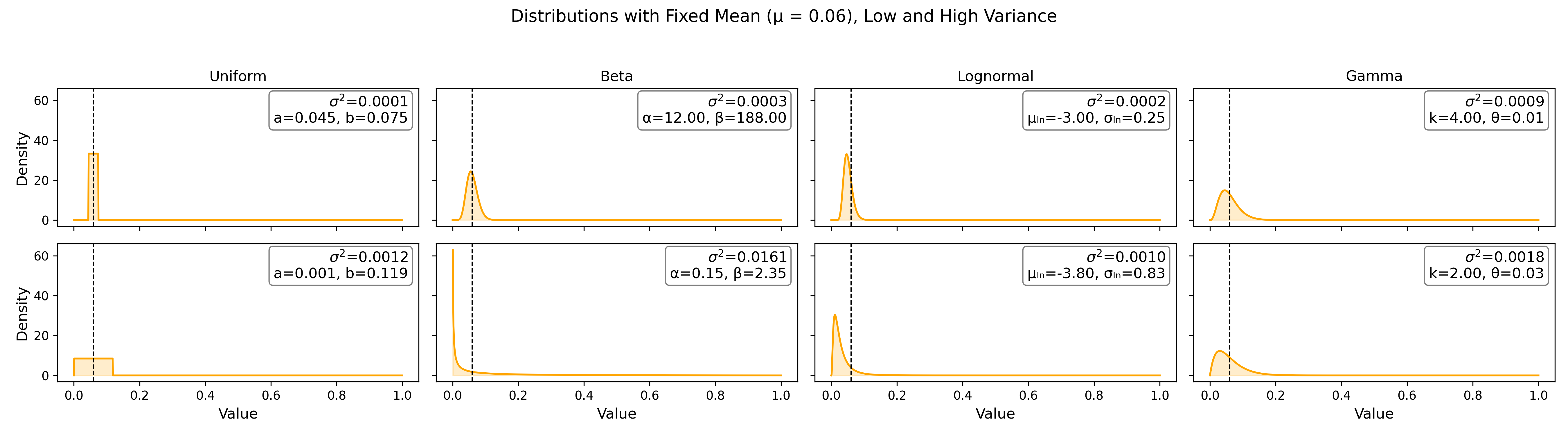

### f3_haploid_simulation_results.png

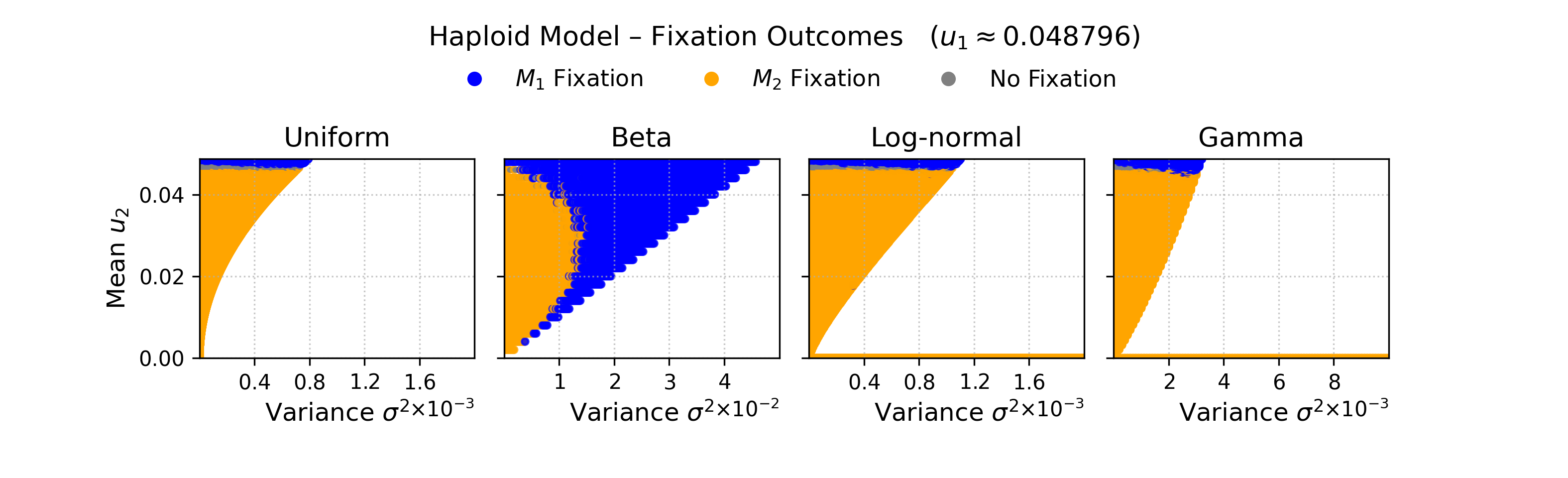

### f4_lyapunov_approx.png

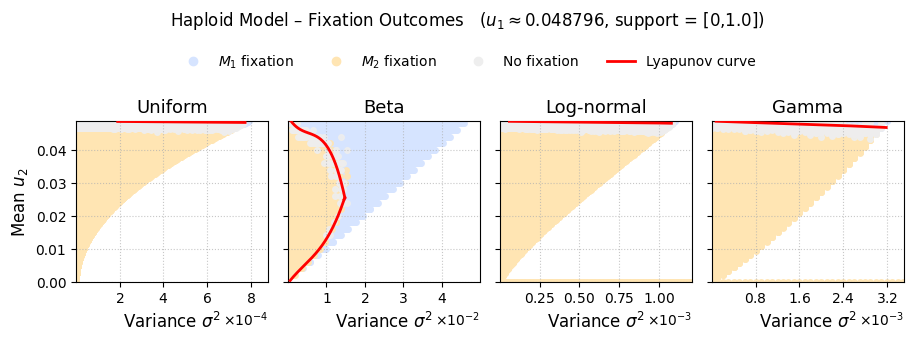
